# RESM: Capturing sequence and structure encoding of RNAs by mapped transfer learning from ESM (evolutionary scale modeling) protein language model

**DOI:** 10.1101/2025.08.09.669469

**Authors:** Yikun Zhang, Hao Zhang, Guo-Wei Li, He Wang, Xing Zhang, Xu Hong, Tingting Zhang, Liangsheng Wen, Yu Zhao, Jiuhong Jiang, Jie Chen, Yanjun Chen, Liwei Liu, Jian Zhan, Yaoqi Zhou

**Affiliations:** Institute of Systems and Physical Biology, Shenzhen Bay Laboratory, Shenzhen 518107, China; School of Electronic and Computer Engineering, Peking University, Shenzhen 518055, China; China Mobile Research Institute, 32 Xuanwumen West Street, Xicheng District, Beijing 100053, China; Advanced Computing and Storage Laboratory, Central Research Institute, 2012 Laboratories. Huawei Technologies Co., Ltd., China; School of Life Science and Technology, ShanghaiTech University, Shanghai, 201210, China; Guangzhou Laboratory, Guangzhou International Bio Island, Guangzhou 510005, China; Ribopeutic (Shenzhen) Co., Ltd., Futian, Shenzhen, Guangdong Province, 518000, China; Ribopeutic Inc., Qiantang, Hangzhou, Zhejiang Province, 310018, China

## Abstract

RNA sequences exhibit lower evolutionary conservation than proteins due to their informationally constrained four-letter alphabet, compared to the 20-letter code of proteins. More limited information makes unsupervised learning of structural and functional evolutionary patterns more challenging from single RNA sequences. We overcame this limitation by mapping RNA sequences to pseudo-protein sequences to allow effective transfer training from a protein language model (protein Evolution-Scale Model 2, protESM-2). The resulting RNA ESM (RESM) outperforms 12 existing RNA language models in zero-shot prediction, not only in sequence classification but also in RNA secondary structure and RNA-RNA interaction prediction. Further supervised fine-tuning demonstrates RESM’s generalizability and superior performance over the existing models compared across multiple downstream tasks, including mRNA ribosome loading efficiency and gene expression prediction, despite RESM being trained exclusively on noncoding RNAs. Moreover, RESM can generalize to unseen sequences beyond its 1,024-nucleotide training limit, achieving 81.3% improvement over state-of-the-art methods in supervised secondary structure prediction for RNAs up to 4,000 nucleotides, limited only by the available GPU memory, while providing >1000-fold speedup compared to MSA-based approaches. RESM provides a robust foundation for deciphering RNA sequence-structure-function relationships, with broad implications for RNA biology.

## Introduction

RNA molecules are central to numerous cellular processes, serving not only as messengers in protein synthesis but also executing critical regulatory, catalytic, and structural functions across all domains of life. Their fundamental biological significance has spurred intense efforts to computationally elucidate complex relationships between RNA sequences, structures, and functions^1,2^. Despite decades of advances, accurately modeling these relationships remains challenging due to intrinsic physio-chemical properties of RNAs. Foremost among these challenges is that 4-letter-coded RNAs have much lower sequence information density than 20-letter-coded proteins. Moreover, most RNA regions possess dynamic, flexible structures and only a few RNA structures (about 3% in the Protein Data Bank^3^) were experimentally solved, making gold-standard training data for secondary and tertiary-structure prediction quite limited^4^.

Traditionally, accurate RNA structure prediction has heavily relied on comparative modeling involving multiple sequence alignments (MSAs), exploiting evolutionary covariation signals to identify conserved base-pairing interactions^5,6^. However, the dependence on extensive, high-quality alignments confines these methods primarily to well-studied RNA families, posing considerable challenges for novel RNA sequences whose homologous data are difficult to locate. For example, Rfam, a standard reference database, which is specifically constructed to catalog RNA families and their homologous sequences, currently encompasses approximately 4,000 RNA families, each with a median of only ∼45 homologous sequences^7^. Furthermore, homology search methods, such as RNAcmap^8^, RNAcmap3^9^, and rMSA^10^, which utilize covariance models implemented in Infernal^11^, become computationally prohibitive for long RNAs, creating significant practical barriers to scaling analyses of large-scale RNA datasets with long RNAs. Moreover, these methods were limited to RNA sequences with accurately known or predicted secondary structures.

Recent breakthroughs in machine learning, particularly transformer-based large language models (LLMs)^12,13,14,15^, have shown transformative potential in biological sequence modeling. Initially developed for natural language processing, these models have demonstrated unparalleled success in capturing deep sequence patterns and contextual dependencies across diverse domains, including protein sequence modeling^16^. The extension of transformer architectures to protein sequences, exemplified by models such as ESM-2 models^17^(ProtESM-2), leverages large-scale pretraining on evolutionary diverse datasets, enabling models to internalize complex evolutionary, structural, and functional constraints inherent in protein families^18,19,20,21,22,23,24,25,26,27,28^.

However, directly applying similar LLM strategies to RNAs faces distinct challenges. RNA sequences typically have much lower complexity, and experimentally validated RNA structures are significantly fewer than proteins. Moreover, RNA structural flexibility and limited conserved evolutionary signals further complicate the direct transfer of the methods from proteins to RNAs. Indeed, a recent zero-shot benchmarking test of 13 RNA language models (LMs)^29^ revealed biased learning on either secondary structure or sequences, regardless if these RNA LMs were trained for specific purposes^30,31^, noncoding RNAs^32,33,34,35,36,37,38,39,40,41^, or universal models for DNA, RNA, and proteins^42,43,44^. In other words, most models performing well on secondary structure prediction would perform poorly in sequence classification and vice versa.

Currently, the majority of RNA-specific LMs such as RNA-FM^32^ and UniRNA^33^ have relied on training from scratch on existing RNA databases, while the model like RNA-MSM^34^ incorporates homologous information from MSAs to enhance structural prediction but remain dependent on the quality and quantity of available MSAs. Unlike RNA, proteins demonstrate significantly higher sequence conservation, facilitating efficient learning of evolutionary and co-evolutionary relationships through Bidirectional Encoder Representations from Transformers (BERT)^15^ architectures. Indeed, unsupervised prediction of protein contact maps from protein LMs significantly surpasses RNA base-pair predictions^29^ from RNA LMs underscoring the potential benefits of transferring learned insights directly from protein models to RNA modeling. Previous efforts, including approaches by Jian et al.^49^ and ProtRNA^38^, attempted to utilize pretrained protein models through partial adaptation strategies, such as freezing protein model backbones or selective layer fine-tuning.

In this study, we introduced RESM (RNA Evolutionary-Scale Model) by employing a direct mapping strategy that converts RNA sequences into pseudo-protein representations so that all ProtESM-2’s parameters can be updated and refined for a comprehensive dataset of 23.51 million noncoding RNA sequences curated from RNAcentral^50^ database. The mapping transformation from a 4-letter to a 20-letter alphabet expands the information-theoretic capacity^51^ from log₂(4)=2 bits to log₂(20)≈4.3 bits per position, enabling the encoding of subtle sequence features including structural propensities and evolutionary constraints that remain compressed in native RNA representations. Indeed, we showed that RESM provided improved capture of both functional classes and base pairing in zero-shot prediction, in constrast to the fact that most previous RNA LMs bias toward one task. Moreover, RESM efficiently handles long RNA sequences beyond its training limit, overcoming a major bottleneck where MSA-based approaches become computationally prohibitive. Further evaluations on various downstream applications, including on mRNAs unseen by RESM, confirmed RESM’s robust and generalizable performance across multiple RNA-centric tasks.

## Results

### Mapped transfer learning from protein ESM-2 to RNAs

As illustrated in Fig. 1a, we employed the same network architecture as ProtESM-2 for continuous training with 23.51 million de-redundant noncoding RNA sequences from RNAcentral database^50^(Supplementary Table 1). To monitor the behavior of the model during training (Fig, 1b), we implemented an attention-based diagnostic pipeline in which self-attention maps extracted from the model were utilized as the input features for RNA secondary structure prediction via a lightweight logistic regression classifier. Model performance was evaluated using F1 scores on a validation set of 40 RNA sequences with experimentally determined three-dimensional structures (VL40). The diagnostic pipeline was applied during pretraining on a small subset of 20 noncoding RNAs selected from the TR405 dataset, which consists of 405 RNAs curated in RNA-MSM^34^. The model is subjected to various zero-shot and supervised benchmark tests as summarized in Fig. 1c.

**Fig. 1.**
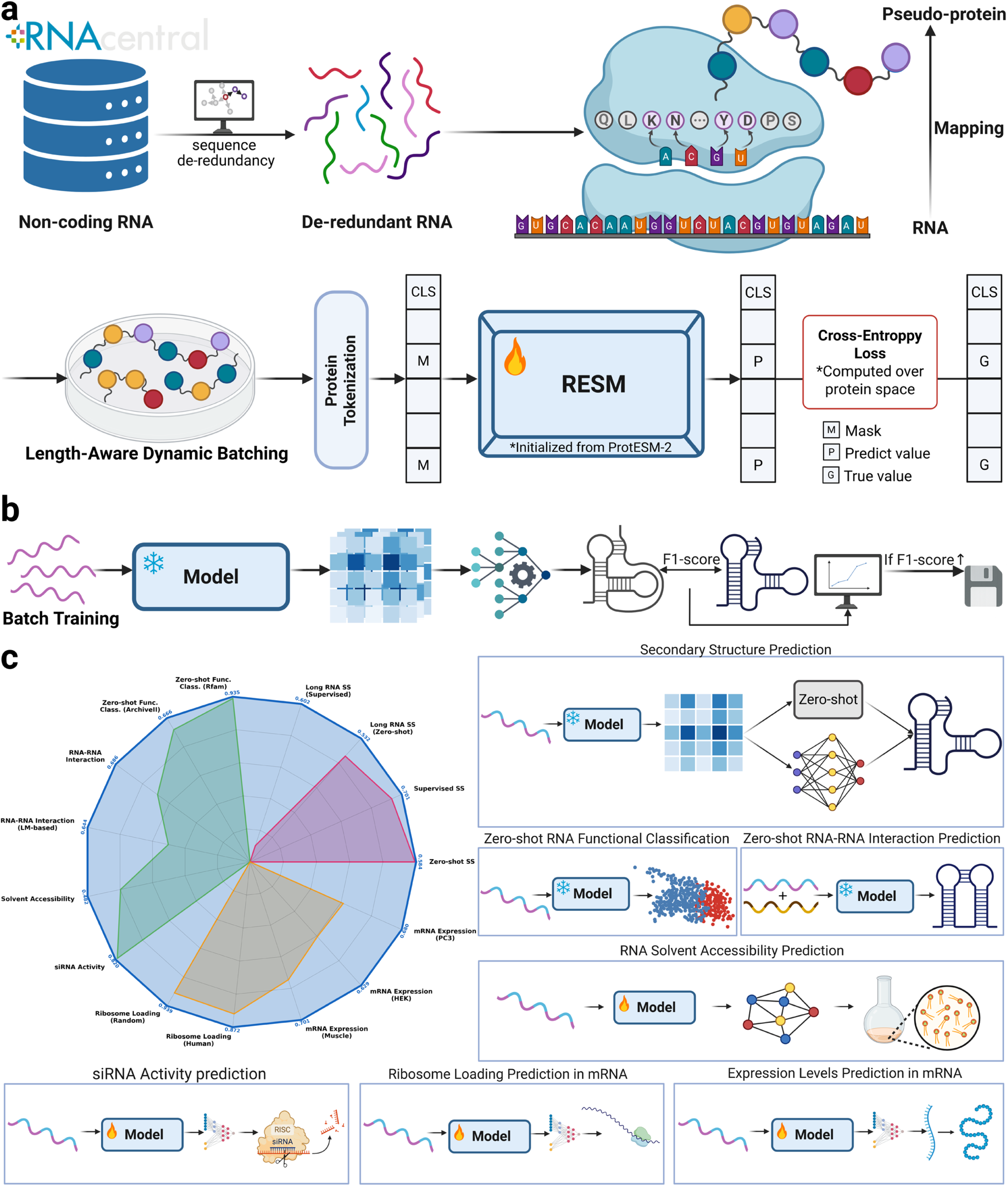
Overview of RESM training framework and its applications to diverse RNA-level structural and functional tasks. **a.** RNA sequences are mapped to pseudo-protein sequence using a defined RNA-to-protein mapping module, enabling full, initial reuse of the parameters from the protein language model ProtESM-2. The RESM model is re-trained on 23.51 million de-redundant noncoding RNA sequences from the RNAcentral database using the same architecture as ProtESM-2, with masked token prediction performed over the protein vocabulary. **b.** To monitor training progress, the structure-aware evaluation module is introduced. Self-attention maps from the model are passed into a lightweight classifier for RNA secondary structure prediction, and performance is tracked via F1 scores on a validation set of 40 RNAs with experimentally determined 3D structures. **c.** Performance comparison of RESM against baseline methods across 15 RNA-related subtasks spanning 8 task categories. RESM (blue line) consistently outperforms or matches the best baselines including MSA-based methods, which are color-coded by task type: secondary structure (SS) prediction tasks (red), function-related tasks (green), and mRNA-related tasks (yellow). The numerical values around the perimeter indicate the best performance achieved for each task, with the color of each value showing which method (RESM in blue or the corresponding baseline) performs best. The resulting RESM representations are evaluated in both zero-shot and supervised settings, covering structural and functional domains: zero-shot and supervised secondary structure prediction, zero-shot RNA functional classification, zero-shot RNA–RNA interaction prediction, and supervised prediction of solvent accessibility, ribosome loading, mRNA expression level, and siRNA activity.

Fig. 2a examined several possible mappings of AUCG to KDNY, KEDR, and VLER as well as without mapping (random initial weights with the same network architecture). The result shows that it was challenging to train without mapping. Meanwhile, we tested a few possible biophysical mappings (see Methods) and found reasonably similar performance, consistent with a previous work when mapping protein-to-RNA in co-evolution analysis^49^. The mapping from AUCG to KDNY was selected as the final choice, according to diagnostic F1 scores.

**Fig. 2.**
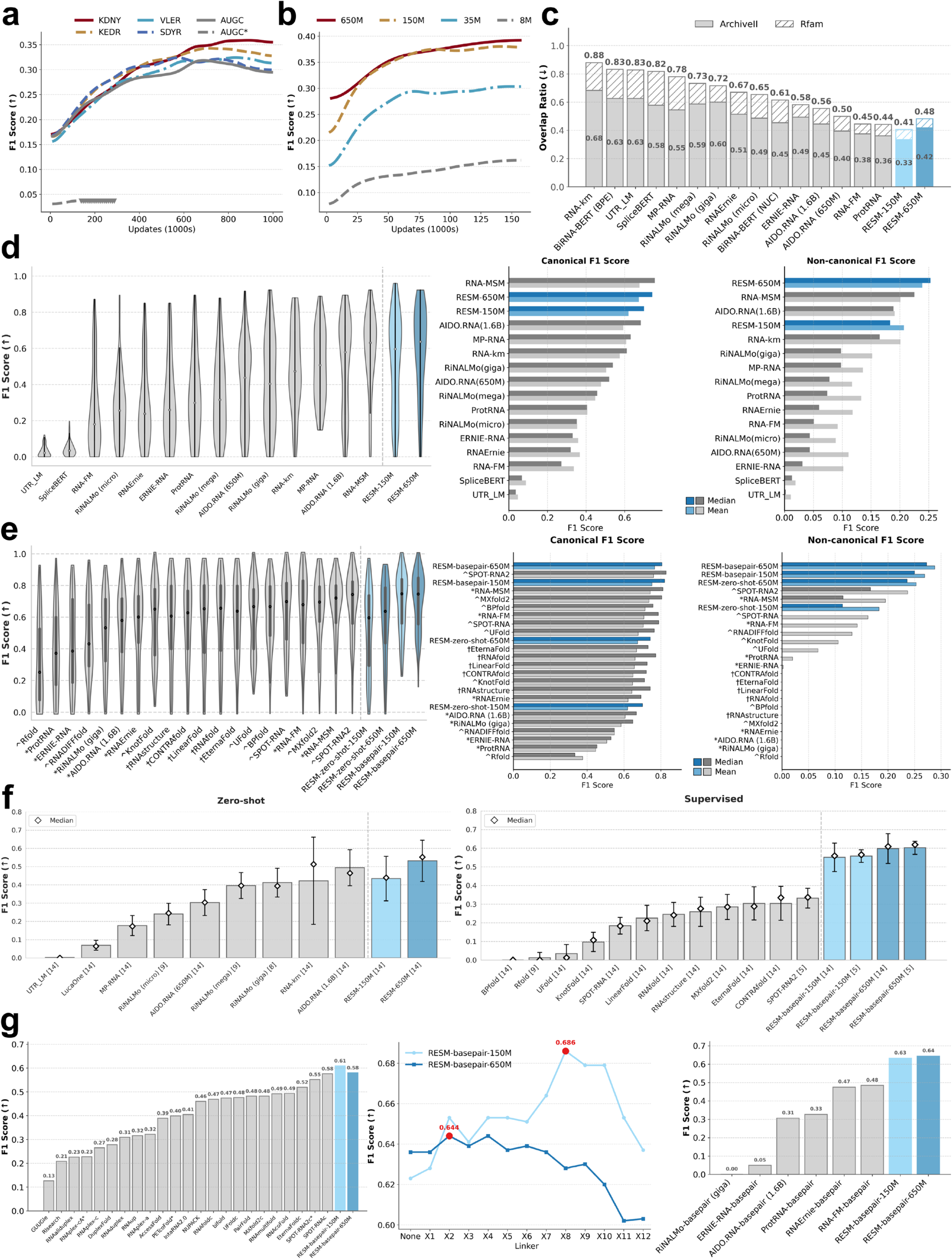
RNA-protein mapping and zero-shot and supervised comparison of RNA language models for structural and functional interference. **a.** Performance of diagonostic secondary structure prediction during pre-training, measured by F1 score on a set of 40 RNAs employing different mapping strategies from AUCG to KDNY, KEDR, VLER, SDYR, and AUCG, along with a randomly initialized baseline (AUCG*), under identical network architectures. The triangle symbol indicates NaN values encountered during training. **b.** Same evaluation as in panel **a.,** but comparing RESM models of different sizes (from 8M to 650M) under the KDNY mapping. **c.** Zero-shot functional classification using RESM embeddings, benchmarked against other RNA language models by using the overlap ratio between family and non-family members (the smaller, the better performance). **d.** The accuracy of zero-shot prediction of RNA secondary structure according to F1 scores directly from attention maps by various RNA language models. The left boxplot summarizes F1 scores between attention-derived contact maps and experimentally determined structures across benchmark RNAs. The right grouped bar charts further separate this analysis into canonical and non-canonical base-pairing types. **e.** Supervised base-pairing structure prediction with RESM-basepair using attention maps. The left boxplot compares F1 scores of different supervised models. RESM-basepair-650 and RESM-basepair-150M achieved the highest median and mean F1 scores, respectively, outperforming RNA-MSM_ResNet and SPOT-RNA2 using multiple sequence alignment. The right bar charts further separate performance into canonical and non-canonical F1 scores.**f.** Performance of RNA secondary structure prediction methods on long RNA sequences (1,256-3,705 nucleotides). Bar plots showing F1 scores for zero-shot (left) and supervised (right) prediction methods on a curated dataset of seven non-redundant long RNA structures. Error bars represent standard deviation, with white diamonds indicating median values. **g.** Zero-shot RNA–RNA interaction prediction with RESM. Left: Comparison of 23 inter-RNA base-pair predictors on 64 RNA complexes unseen by SPOT-RNA/SPOT-RNA2. Middle: Performance of RESM-basepair (150M and 650M) with different linker configurations. Right: Supervised RNA language models fine-tuned for secondary structure prediction (“-basepair” suffix), evaluated directly on 62 RNA-RNA complexes without additional training (excluding two sequences >510 nt).

We further examined the dependence of the performance on secondary structure prediction on the model size ranging from 8M, 35M, 150M to 650M (Fig. 2b). The improvement from the 8M model to the 35M model is the largest and the smallest from the 150M model to the 650M model, suggesting that the gain from beyond the 650M model may not be cost effective at least in term of secondary structure. Given the huge training cost, we limited the model size to 650M in this study.

### Zero-shot RNA functional classification

To evaluate whether RESM can capture biologically meaningful RNA features for discriminating homologous from nonhomologous relationships, we conducted zero-shot functional classification experiments by direct use of the output embeddings of RESM (Fig. 2c and Supplementary Table 2). We evaluated model performance on two benchmark datasets: the Rfam database^7^ containing 24,607 sequences of 4,170 families and ArchiveII^52^ comprising 3,864 RNA structures of 9 RNA types (see Methods). Classification performance was quantified through the overlapping ratio (OR) between the family and non-family members, where lower OR values indicate better separation between homologous and non-homologous sequence distributions with 0 for no overlap and 1 for 100% overlap. We conducted comprehensive evaluations against 12 distinct RNA LMs. For models available in multiple settings (BiRNA-BERT: BPE/NUC; RiNALMo: micro/mega/giga; AIDO.RNA: 650M/1.6B), we tested all variants to ensure robustness of our results across different model capacities.

On the Rfam dataset for Rfam classification, RESM-650M achieved a benchmark OR of 0.065 ± 0.000, representing a 7.1% improvement over the next best RNA-FM^32^ (OR=0.070 ± 0.000), which has a similar performance to RESM-150M (OR=0.072 ± 0.000). Interestingly, on the smaller yet more challenging ArchiveII dataset, it is RESM-150M having the best performance (OR = 0.334 ± 0.002) and 7.5% improvement over the next best ProtRNA^38^ (OR=0.361 ± 0.000). ArchiveII test dataset is more challenging because the average sequence identity between homologous sequences is significantly lower (54.5%) than that in the Rfam dataset (77.1%). The decreased classification performance of RESM-650M on the ArchiveII set (OR=0.417 ± 0.000) may be also due to the small size of the set (3,864 RNAs of only 9 RNA types), also highlighting the challenge of functional classification from sequence information alone.

### Zero-shot secondary structure prediction

While RESM’s training process involved monitoring the performance in RNA secondary structure prediction, the model itself was never directly trained on secondary structure data. Nonetheless, inherent structural information might be implicitly captured within the model’s attention layers. We selected the best-performing attention map based on the validation set (VL40) and assessed its predictive capability on an independent test set comprising 70 RNAs (TS70). In this zero-shot scenario, we benchmarked RESM against 11 other RNA LM architectures across 14 parameter configurations, ranging from 1.2M to 1.6B parameters to evaluate its ability to infer RNA secondary structures from single sequences without explicit structural inputs.

RESM-650M achieved state-of-the-art performance (median F1 = 0.637; mean F1 = 0.584 ± 0.248; Fig. 2d and Supplementary Table 2), matching MSA-based RNA-MSM^34^ (median F1 = 0.631; mean F1 = 0.582 ± 0.217; P = 0.9505). A detailed evaluation, distinguishing canonical and non-canonical base pairs across 11 RNA LMs, further demonstrated RESM’s robustness (Supplementary Table 3). Although RESM variants exhibited a modest performance shortfall relative to RNA-MSM in canonical base-pair predictions (median F1: 0.743 vs. 0.756), they substantially outperformed RNA-MSM in non-canonical base-pair predictions by 18.0% (median F1: 0.236 vs. 0.200).

When compared exclusively against single-sequence-based RNA LMs, RESM-650M demonstrated superior prediction performance. Specifically, for canonical base pairs, RESM-650M improved median (mean) F1 scores by 6.1% (8.7%) over RESM-150M and by 8.6% (13.7%) over the next best, a larger AIDO.RNA^35^ model (1.6B parameters). The advantage was even more pronounced for non-canonical base pairs, where RESM-650M surpassed RESM-150M by 107.0% (38.3%) and AIDO.RNA by 12.9% (33.9%) in median (mean) F1 scores. Overall, RESM-650M achieved a median (mean) F1 improvement of 6.8% (12.1%) relative to RESM-150M and 10.0% (15.0%) compared to AIDO.RNA for base-pair prediction, highlighting its effectiveness in capturing RNA structural information from single sequence data alone.

### Supervised secondary structure prediction

To further improve secondary structure prediction, we developed a deep neural network model (RESM-basepair) leveraging attention maps from RESM for supervised RNA secondary structure prediction. Our training framework and protocol followed SPOT-RNA^53^ augmented with attention map inputs (see Methods). The model underwent pre-training on the TR0 dataset filtered from bpRNA-1m^54^, transfer learning on the TR405 dataset, and was evaluated on the TS70 dataset, consistent with the zero-shot evaluation benchmark. Notably, RESM-basepair-150M (median F1: 0.748; mean F1: 0.686 ± 0.190) and RESM-basepair-650M (median F1: 0.746; mean F1: 0.701 ± 0.180) achieved the best results (Fig. 2e and Supplementary Table 4), that improves over MSA-based, supervised RNA-MSM model (median F1: 0.721; mean F1: 0.662 ± 0.211). In the TS70 dataset, the supervised RESM-basepair-150M and RESM-basepair-650M models improved upon their zero-shot counterparts by 31.7% and 20.0% in mean F1 score, respectively (Fig. 2e and Supplementary Table 4).

Further analysis revealed distinct performance differences between canonical and non-canonical base pairs (Supplementary Table 5). RESM-basepair-650M achieved a strong performance on canonical pairs (median F1: 0.806; mean F1: 0.766 ± 0.180), compared to the MSA-based model, SPOT-RNA2 (median F1: 0.814; mean F1: 0.740 ± 0.218) on mean F1 score, despite some overlap between the latter’s training set and TS70. RESM-basepair-150M exhibited competitive performance on canonical pairs (median F1: 0.820; mean F1: 0.752 ± 0.192). Remarkably, RESM-basepair-650M demonstrated superior performance on non-canonical pairs (median F1: 0.273; mean F1: 0.288 ± 0.275), significantly outperforming other methods, including MSA-based supervised RNA-MSM (median F1: 0.115; mean F1: 0.195 ± 0.225) and MSA-based SPOT-RNA2 (median F1: 0.167; mean F1: 0.237 ± 0.246). These results underscore the RESM-basepair model’s enhanced ability to accurately predict non-canonical RNA structures, even without using MSAs.

To evaluate the generalizability of RESM-basepair beyond its training length limit, we assessed its performance on a long RNA Dataset. The dataset spans lengths from 1,256 to 3,705 nucleotides with a resolution better than 5 Å (Supplementary Table 6). In zero-shot prediction, RESM-650M achieved a mean F1 score of 0.532 ± 0.113, outperforming the second-best baseline method by 7.7%. Among RNA language models, AIDO.RNA (1.6B) showed moderate performance (0.494 ± 0.099), while others such as RNA-FM are limited to sequences shorter than 512 nucleotides.

For supervised prediction, RESM-basepair-650M demonstrated even more improvements with a mean F1 score of 0.602 ± 0.035, representing an 81.3% improvement over the best competing method, SPOT-RNA2 (0.332 ± 0.053) on the 5 long RNAs that SPOT-RNA2 could process. Importantly, SPOT-RNA2, despite being MSA-based and computationally intensive, could only be evaluated on 5 RNAs shorter than 2,000 nucleotides due to prohibitive computational costs of multiple sequence alignment for longer sequences. In contrast, RESM-basepair processed all 14 sequences efficiently and achieved 96.7% improvement over the second-best method CONTRAfold (0.304 ± 0.091) across all 14 long RNAs (Fig. 2f and Supplementary Table 7). Notably, many established methods showed severe performance degradation on these long sequences: thermodynamic approaches (BPfold, Rfold, UFold) failed completely with F1 scores near zero, while probabilistic methods such as KnotFold and SPOT-RNA achieved only 0.097 ± 0.052 and 0.184 ± 0.045, respectively. This capability addresses a critical computational bottleneck, as homology searches for sequences exceeding 1,000 nucleotides using RNAcmap3 or similar methods require prohibitive computational resources, often taking days to complete for a single sequence.

### Zero-shot RNA**-**RNA interaction prediction

To further assess the generalizability of RESM-basepair, we applied it to RNA–RNA interaction (RRI) prediction by concatenating two RNA chains into a single sequence (see Methods). Following the benchmarking protocol established by Lang et al.^56^, we utilized a rigorously curated gold-standard dataset of 64 non-redundant RNA–RNA interaction pairs from high-resolution 3D RNA structures, explicitly chosen to minimize structural redundancy with SPOT-RNA and SPOT-RNA2 training datasets. This approach leverages the secondary structure prediction capabilities of supervised models to infer inter-molecular base pairs without explicit training on RNA-RNA interaction data.

We first compared RESM-basepair models (150M and 650M) against 23 non-LM methods across multiple synthetic linker configurations using precision, recall, overall F1 score, and Matthews correlation coefficient (MCC). Incorporation of synthetic linkers boosted prediction accuracy; RESM-basepair (150M and 650M) achieved the highest median individual F1 scores of 0.686 ± 0.355 (𝑋^8^ linker) and 0.644 ± 0.368 (𝑋^2^ linker), respectively (Fig. 2g and Supplementary Table 8). Although removing linkers led to slight performance degradation (Supplementary Table 9 and Supplementary Table 10), RESM-basepair-150M consistently outperformed SPOT-RNA^53^, the second best among 23 methods compared. Overall, RESM-basepair-150M surpassed SPOT-RNA’s median individual F1 score (0.579 ± 0.323) by 18.5%, highlighting the superior ability of RESM in RNA-RNA interaction prediction.

Additionally, to ensure fair comparison, we evaluated RESM-basepair against other RNA language models that were similarly fine-tuned for supervised secondary structure prediction, all tested without synthetic linker sequences (Supplementary Table 11). Among these LM-based methods, RESM-basepair-150M achieved the highest precision (0.705), overall F1 score (0.607), and Matthews correlation coefficient (MCC: 0.613). The larger RESM-basepair-650M model demonstrated robust performance with an overall F1 score of 0.599, MCC of 0.604, and the highest median individual F1 score (0.644 ± 0.360). Notably, RESM-basepair-150M outperformed the next best RNA foundation model (RNA-FM) by 13.0% in overall F1 score (0.607 vs 0.537). These results demonstrate the superior capability of RESM-basepair models in accurately capturing RNA–RNA interactions, highlighting their potential for diverse applications in RNA structure prediction and functional analysis.

### Supervised RNA solvent accessibility prediction

To further validate RESM’s ability to capture RNA structural features, we assessed its performance in predicting RNA solvent accessibility, a critical measure of nucleotide solvent exposure in 3D structures (Fig. 3a). We employed the RNA-MSM benchmark dataset^34^, consisting of experimentally determined RNA structures (405 training, 40 validation, and 70 test RNAs), carefully filtered for structural and sequence redundancy (see Methods). Comparative analysis demonstrated that the trained model called RESM-ASA-650M achieved the highest Pearson Correlation Coefficient (PCC = 0.482) and the lowest Mean Absolute Error (MAE = 31.0 Å^2^), significantly outperforming RESM-ASA-150M, RNA-MSM, RNAsnap2-pro^57^, and RNA-FM LMs (Supplementary Table 12). The statistical significance of improvement over MSA-based RNA-MSM (t = 3.090, P = 0.003) confirms RESM-ASA-650M’s superior accuracy and consistency in modeling RNA solvent accessibility, even without MSAs as the input.

**Fig. 3.**
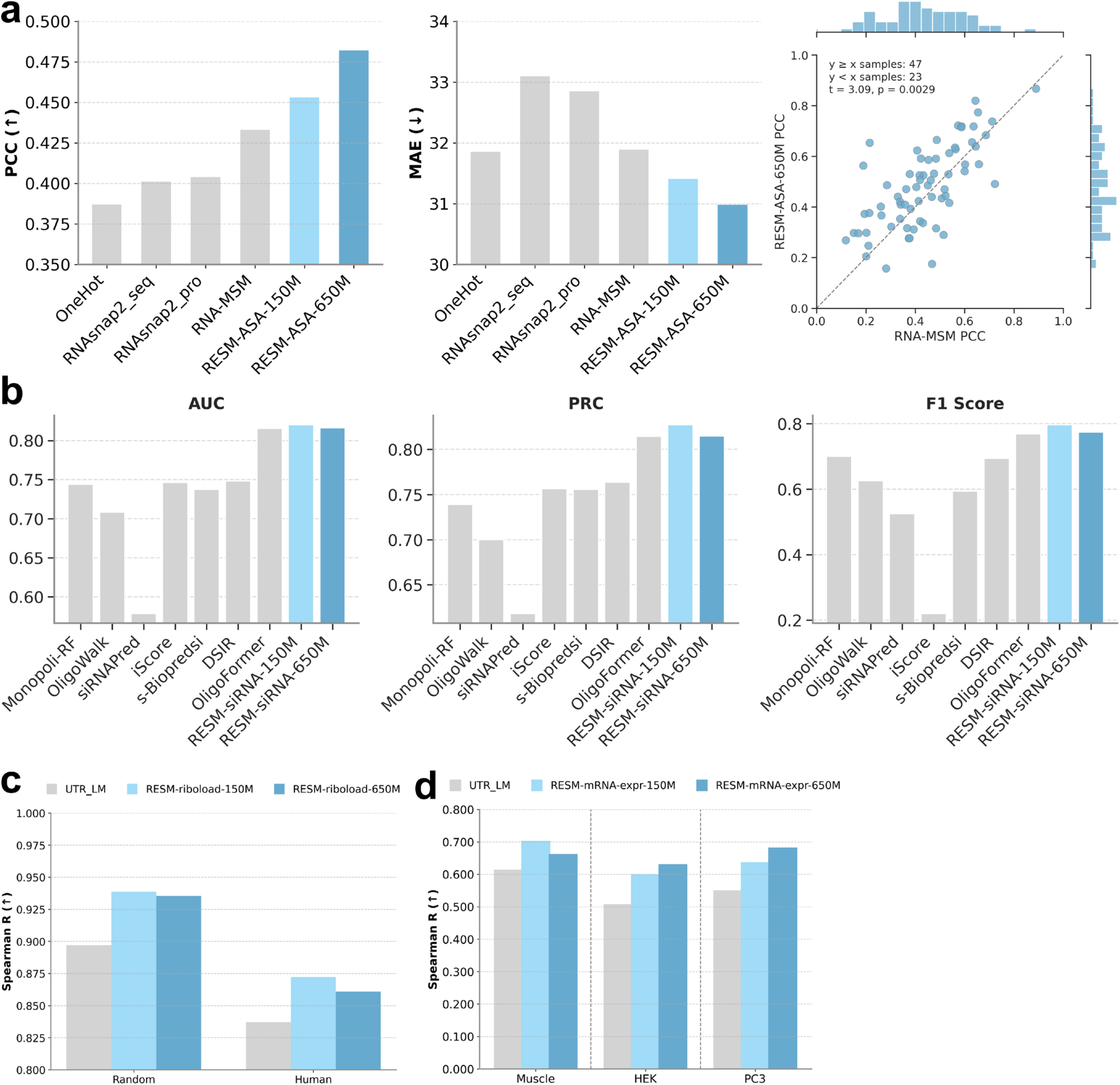
RESM performance across diverse RNA-level prediction tasks. **a.** RNA solvent accessibility prediction on the RNA-MSM benchmark. RESM-ASA (650M) outperforms RNA-MSM, RNAsnap2, and one-hot encoding baselines according to Pearson’s correlation coefficient (PCC), Mean Absolute Error (MAE), confirming RESM’s enhanced structural awareness. **b.** siRNA activity prediction under cross-dataset evaluation (trained on Huesken, tested on Mixset). RESM-siRNA (150M) achieves state-of-the-art AUC_ROC, AUC_PRC, and F1 scores, highlighting strong generalization across domains. **c.** Ribosome loading prediction evaluated according to Spearman’s correlation coefficient on synthetic random sequences and human-derived 5’UTRs under the fine-tuning setting using the non-redundant dataset. **d.** mRNA expression level prediction (RESM-mRNAexpr) evaluated according to Spearman’s correlation coefficient across human muscle tissue, PC3 cells, and HEK293T cells using full-length sequence input. RESM task variants (150M: light blue; 650M: dark blue) consistently outperform all models compared including the specialized UTR-LM model (gray).

### Supervised siRNA activity prediction

Predicting siRNA activity is essential for optimizing RNA interference strategies and developing siRNA drugs. To evaluate RESM’s capability in predicting siRNA activity, we conducted inter-dataset validation, training the model on the Huesken dataset^58^ and validating it on the Mixset dataset^59^ (Fig. 3b). RESM’s performance on this task (RESM-siRNA) was evaluated across several models. RESM-siRNA-150M achieved superior performance with an AUC_ROC of 0.820, AUC_PRC of 0.827, and F1 score of 0.797, outperforming existing methods including Monopoli-RF^60^ (AUC_ROC: 0.744), DSIR^61^ (AUC_ROC: 0.748), and transformer-based OligoFormer^59^ (AUC_ROC: 0.815) (Supplementary Table 13).

### Beyond noncoding RNAs: prediction of supervised ribosome loading and expression levels in mRNA

Given RESM’s ability to capture both sequential (homology detection and classification) and structural (base-pairing) encoding, we further evaluated RESM’s generalization capability by going beyond noncoding RNAs employed in training and assessed its performance in coding mRNAs unseen by RESM: prediction of ribosome loading quantification and expression levels. For both tasks, we employed the same downstream task prediction head (see Methods) for predicting ribosome loading and expression levels.

For ribosome loading prediction, we utilized a synthetic dataset composed of 76,319 randomized 5’UTR sequences spanning 25 to 100 nucleotides upstream of the start codon^62^. The RESM model was fine-tuned on this dataset, and its generalization performance was tested on two held-out sets: 7,600 random sequences and 7,600 human-derived 5’UTRs curated from published benchmarks^62^. We also included a comparison with six representative models, including UTR-LM^30^, MTtrans^63^, Optimus^62^, Framepool^64^, RNABERT^65^ and RNA-FM^32^. During analysis, we observed substantial sequence similarity between the training and test sets, which raised concerns about potential data leakage. To address this, we applied an 80% sequence identity threshold using the CD-HIT-EST^66^ tool to eliminate redundant sequences between while maintaining the original distribution of the two test sets. This resulted in a reduced training set of 70,372 sequences, representing a 7.8% reduction with fixed test sets. Results from both training settings are reported in Supplementary Table 14 to facilitate comparison to previous studies. Based on the previous redundant training set, RESM-riboload-150M achieved a Spearman R of 0.941 on the random test set and 0.875 on the human test set, outperforming the best-performing baseline UTR-LM model, which reached 0.911 and 0.847, respectively. Based on the new non-redundant training set, the RESM-riboload-150M model achieved Spearman R of 0.939 (random dataset) and 0.872 (human dataset), surpassing the specialized UTR-LM model by 4.7% and 4.2%, respectively (Fig. 3c). The larger RESM-riboload (650M) model has essentially the same performance as RESM-riboload (150M) model with Spearman R values at 0.936 and 0.865, respectively. Notably, while both models experienced a slight performance decline when trained on the new filtered dataset, RESM-riboload-150M model showed better robustness. The decrease in Spearman R for RESM-riboload-150M was only 0.2% (from 0.941 to 0.939) on the random test set and 0.3% (from 0.875 to 0.872) on the human set. In contrast, UTR-LM’s performance dropped by 1.5% (from 0.911 to 0.897) and 1.2% (from 0.847 to 0.837), respectively. These results suggesting that RESM-riboload is less prone to overfitting and better generalized to unseen data, even when sequence redundancy is removed.

In mRNA expression level prediction, we utilized three biologically diverse datasets spanning human muscle tissue (muscle), prostate cancer (PC3) cells, and human embryonic kidney (HEK) 293T cells^67^. Following the setup used by UTR-LM and other methods, we initially truncated all mRNA sequences to a fixed input length of 100 nucleotides as done in UTR-LM. However, this approach introduced a substantial number of duplicate sequences with inconsistent expression values due to loss of the full sequence (see Methods). To maintain consistency with previous benchmarks, we first evaluated performance under the fixed-length setting, where RESM-mRNAexpr-150M outperformed UTR-LM across all datasets, achieving Spearman correlation coefficients of 0.692 on the muscle dataset, 0.661 on HEK 293T cells, and 0.645 on PC3 cells, corresponding to absolute improvements of 3.1%, 1.0%, and 1.5%, respectively, over UTR-LM’s results of 0.661, 0.651, and 0.630 (Supplementary Table 15). These results demonstrate the strong generalization ability of RESM-mRNAexpr across distinct tissues, even when constrained to limited input lengths. To better preserve regulatory context, we further conducted experiments using full-length 5’UTR inputs, a departure from previous approaches restricted to the first 100 nucleotides. Under this setting, RESM-mRNAexpr-150M achieved Spearman R of 0.701 for muscle tissue, 0.598 for HEK 293T cells, and 0.634 for PC3 cells, reflecting relative improvements of 14.5%, 18.4%, and 15.5% over UTR-LM (0.612, 0.505, and 0.549). The larger RESM-mRNAexpr-650M model further improved performance in HEK and PC3 cells, reaching 0.629 and 0.680, respectively, although its performance on muscle tissue was reduced slightly to 0.660 (Fig. 3d). Compared to UTR-LM, these results correspond to relative gains of 7.9%, 24.6%, and 23.9%, respectively, highlighting RESM-mRNAexpr’s capacity to scale effectively and leverage extended sequence contexts for more accurate expression-level inference across diverse biological systems.

## Discussion

This study was motivated by a recent survey of RNA languages whose zero-shot performances on sequence classification and secondary structure prediction often go in the opposite direction. This is likely because lacking sequence conservation in RNA homologs makes it more difficult to establish a link between sequence and structure. To overcome this challenge, we present RESM, a multi-scale RNA LM that built upon the pre-trained protein ESM-2 architecture and was further trained on a comprehensive dataset comprising 23.51 million de-redundant noncoding RNA sequences. Two parameter sets of models (RESM-150M and RESM-650M) were trained on ESM-2 150M and ESM-2 650M models, respectively.

These two models were tested by both zero-shot prediction and supervised down-streaming tasks. Zero-shot prediction tasks include sequence classification, RNA base pair prediction, and RNA-RNA interaction prediction. Supervised down-streaming tasks include secondary-structure base-pairs, solvent accessibility, siRNA activity, ribosome loading and expression levels in mRNA. This new LM demonstrated a state-of-the-art performance in all structural and functional tasks in both unsupervised and supervised settings when comparing to 12 RNA LMs in various tasks. It should be noted that the model was wholly trained on noncoding RNA sequences and yet performed better than existing models that were trained on mRNAs for mRNA-related tasks.

One noticeable performance improvement is that supervised RESM models on base-pair and solvent accessibility prediction can perform better than or comparable to those relied on MSAs such as SPOT-RNA2^68^ and RNAsnap2^57^ as well as the LM based on MSAs (RNA-MSM). A particularly significant advantage of RESM is its ability to generalize to RNA sequences far exceeding its training length. While the supervised RESM-basepair models were trained exclusively on sequences ≤1,024 nucleotides, they demonstrated exceptional performance on RNAs of long sequences with the longest RNA at about 4,000 nucleotides (a limit imposed by available GPU memory), outperforming state-of-the-art methods by 7.7% and 81.3% in zero-shot and supervised predictions, respectively. This capability transforms the landscape of RNA structure prediction for long sequences. Traditional MSA-based approaches face computational explosion when processing RNAs longer than 1,000 nucleotides—homology searches using tools like RNAcmap3 can require days of CPU time for a single long RNA, making them impractical for genome-scale analyses or time-sensitive applications. In contrast, RESM completes predictions within minutes on a single NVIDIA GeForce RTX 4090 GPU, with RESM-basepair-150M processing a 1,600-nucleotide RNA in just ∼100 seconds while maintaining superior accuracy. This highlights the superior ability of RESM models to capture co-evolutionary information, eliminating the computational bottleneck of MSA construction while maintaining superior accuracy across RNA lengths that are critical for time-sensitive applications involving large datasets.

The reason for the above superior performance is the transfer learning of co-evolution from proteins to RNAs by direct mapping between amino acids and nucleotides as shown in Fig. 2. There are 116,280 (20 × 19 × 18 × 17) unique ways to map RNA (4 bases) to proteins (20 amino acids). It is not possible to test all possible mapping. Jian et al.^49^ tested a few mappings for their studies of employing a pretrained model of protein contact-map prediction for transfer learning of RNA-contact prediction. They have tested random mapping (e.g. AUCG to HETL) and group-based mapping (AUCG is mapped onto charged, polar uncharged, hydrophobic, and special amino acid residues, randomly: e.g. AUCG to RDSY, AUCG to KDNY). We tested a few their choices (AUCG to SDYR, AUCG to KDNY) as well as our own biophysical mapping (AUCG to KEDR). For example, Watson-Crick base pairings A-U and C-G in RNA were mapped to the salt bridge interactions of K-E and D-R in proteins, respectively, or mapped to hydrophobic interactions (V-L) and salt-bridge (E-R), respectively. As a baseline control, we also evaluated a direct identity mapping, AUCG mapped to AUCG. To assess the interaction between mapping strategies and model initialization, we conducted experiments both with and without loading pretrained weights from ProtESM-2. The AUCG to KDNY mapping was selected based on its performance on diagnostic secondary structure prediction (Fig. 2a). We noticed that this mapping reflects a mapping of all bases to all charged (KD) or hydrophilic (NY) residues. This is consistent with the fact that all nucleotide bases can be considered as hydrophilic given their hydrophilic sugar-phosphate backbone. More interestingly, the stable Watson-Crick AU base pair corresponds well with the salt bridge in proteins between the positively charged lysine (K) and the negatively charged aspartic acid (D). However, the Watson-Crick GC base pair does not have a such neat salt-bridge biophysical correspondence. Other more biophysical mappings such AUCG to K(+)E(-)D(-)R(+) to ensure AU, GC, and the wobble base pair GU as salt bridge or AUCG to V(hydrophobic)L(hydrophobic)E(-)R(+) did not end up in improved performance. Nevertheless, our test is quite limited. It is entirely possible that there is a mapping allowing improved transfer learning. Additional searches for potentially optimal mappings were not pursued due to computational constraints.

It is noted that RNA LM ProtRNA also utilized the pre-trained ESM-2. However, there are key distinctions between ProtRNA and RESM. More specifically, unlike our direct mapping module, which aligns nucleotides with specific amino acids, ProtRNA expands the vocabulary by introducing lowercase nucleotide tokens (“A”, “U”, “G”, “C”, and “X”) to distinguish nucleotides from uppercase amino acid tokens, which were accommodated by extending embedding and bias layers. Moreover, ProtRNA freezed the ESM-2 backbone and fine-tunes only the last four layers, whereas RESM undergoes full-parameter continuous pretraining on a significantly larger noncoding RNA dataset (23.51 million vs. 6 million sequences). This more extensive pretraining allowed RESM to fully internalize the semantic structure of RNA, enabling richer representations and improved generalization across RNA-related tasks. Comparison of method performance indicates that our direct mapping and transfer of all parameters allow improved transfer learning of co-evolution information from proteins to RNAs.

Interestingly, when comparing RESM-150M and RESM-650M across various tasks, we observed that larger models do not always outperform smaller ones, particularly when evaluated on limited datasets. This reflects a well-known challenge in deep learning. Although larger models possess greater representational capacity, they are often more susceptible to overfitting when training data is scarce. For example, in the zero-shot RNA–RNA interaction prediction task, RESM-basepair-150M achieved a higher precision of 0.733 than 0.695 given by RESM-basepair-650M, despite the latter having substantially more parameters. Similarly, in the siRNA activity prediction task, RESM-siRNA-150M slightly outperformed RESM-siRNA-650M in terms of AUC_ROC (0.820 compared to 0.816). This pattern is especially pronounced in the task of predicting the mRNA expression level. On the Muscle dataset, which contains only 1,257 RNA sequences, RESM-mRNAexpr-150M achieved a higher Spearman’s rank correlation coefficient of 0.701, compared to 0.660 given by RESM-mRNAexpr-650M. However, on the larger HEK and PC3 datasets, RESM-mRNAexpr-650M consistently outperformed RESM-mRNAexpr-150M (0.629 versus 0.598 on HEK, and 0.680 versus 0.634 on PC3), demonstrating improved generalization when sufficient training data are available.

While this work establishes a robust foundation for protein-to-RNA transfer learning, it remains an open question whether mapping strategies can be jointly optimized with pretraining objectives in a differentiable framework. Future efforts may also explore cross-modal models capable of jointly encoding RNA sequence, structure, and expression dynamics. Furthermore, inherent and accurate RNA base-pair information contained in RESM makes it an ideal candidate for integration into RNA tertiary structure prediction modules, where it could provide more accurate secondary structure data or serve as an input for end-to-end tertiary structure prediction. These advancements are expected to significantly broaden the utility of RESM, particularly in applications such as RNA tertiary structure prediction, functional annotation, and therapeutic RNA design.

## Methods

### Pre-training data collection and preprocessing

To construct the pretraining dataset for RESM, we downloaded RNA sequences from the RNAcentral Database version 18.0, a globally recognized repository encompassing diverse non-coding RNA (ncRNA) types across a wide taxonomic range. This database integrates sequences from multiple curated resources, ensuring comprehensive coverage of functional and evolutionary variations in ncRNAs, including tRNAs, rRNAs, lncRNAs, and miRNAs.

For quality control, rigorous deduplication was performed using CD-HIT-EST with an 80% sequence identity threshold. This step minimized redundancy while preserving meaningful biological diversity, which is critical for training a generalizable model. After deduplication, the dataset comprised 23,510,939 sequences. The ncRNA sequences included in the dataset are between 11 and 347,561 nucleotides.

Sequences were preprocessed to standardize inputs for tokenization. Thymine (“T”) was replaced with uracil (“U”) to reflect RNA biochemistry, and non-canonical or ambiguous nucleotides (“R”, “Y”, “K”, “M”, “S”, “W”, “B”, “D”, “H”, “V”, “N”) were unified into a single token, “X”, to simplify rare base handling. Lowercase characters were converted to uppercase to ensure consistency. The final RNA token vocabulary included five primary nucleotides (“A, “U, “G, “C, “X) and five special tokens: <CLS> (classification), <PAD> (padding), <EOS> (end-of-sequence), <UNK> (unknown), and <MASK> (masking).

### RNA 3D structure data for base pair prediction and solvent accessibility prediction

We utilized RNA 3D structural data from RNA-MSM, derived from experimentally resolved RNA structures deposited in the Protein Data Bank^3^, to support both base pair and solvent accessibility prediction tasks. All RNA sequences were shorter than 510 nucleotides and were processed using the pre-trained RESM model. For base pair prediction, self-attention maps from each head and layer were extracted, while embeddings from the final transformer layer were used for solvent accessibility prediction.

Dataset construction followed the protocol established by RNA-MSM. All RNA-containing structures available in the PDB as of 9 August 2021 were collected and randomly split into a training set (TR, 80%), a validation set (VL, 10%), and an initial test set (TS1, 10%). To reduce redundancy, sequences in VL and TS1 with more than 80% sequence identity were removed using CD-HIT-EST. Structural redundancy was further minimized by excluding any sequences from the training set with a TM-score greater than 0.45 relative to those in VL or TS1, as computed by RNA-align. The same structural filtering was applied between VL and TS1 to ensure strict separation. Additional RNA structures deposited in the PDB between 9 August 2021 and 14 July 2022 were also included. These additional entries were filtered using identical sequence and structural redundancy criteria against all TR, VL, and TS1 sequences, resulting in 31 non-redundant RNAs designated as TS2. The final test set (TS70) combines TS1 and TS2, totaling 70 RNAs with experimentally determined tertiary structures that are non-redundant with training and validation sets. In total, the dataset consists of 405 RNAs for training (TR405), 40 for validation (VL40), and 70 for testing (TS70). VL40 was used to determine evaluation thresholds, and TS70 served as the benchmark for model performance.

To evaluate the generalizability of RESM-basepair beyond its training length limit, we built a test set called the long RNA Dataset containing RNA structures with lengths >1024 nucleotides from the RNA3DB structure database^55^. The dataset comprises diverse RNA types including ribosomal RNAs (LSUa and SSU rRNAs from bacteria, archaea, and eukarya), viral RNAs (IRES elements from Cripavirus and CcsR1), snoRNAs, and other functional RNAs, spanning lengths from 1256 to 3705 nucleotides (Supplementary Table 6). We applied a resolution requirement of 5 Å or better along with stringent redundancy filters based on sequence and structural similarity (sequence identity <80%, TM-score <0.45) from the training sequences and structures. We excluded one RNA longer than 10,000 because it demands an internal memory greater than what we have for a CPU. We finally obtained 14 non-redundant long RNA structures for evaluation.

### Model architecture

RESM consists of four main components: an RNA-to-protein mapping module, an embedding module, a transformer module^69^, and task-specific heads. The input RNA sequence 𝑆 = (𝑠_1_, 𝑠_2_, . . ., 𝑠_𝑛_), where 𝑠_𝑖_ is the 𝑖-th RNA base chosen from the 4 primary nucleotides is fed into the modules consecutively. We describe the details of input tokenization and modules as follows.

### RNA-to-protein mapping module

The algorithm first converts RNA and protein token vocabularies into index-based dictionaries. For canonical nucleotide tokens (A, U, C, G), predefined mappings override default indices:

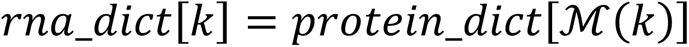

where (ℳ:RNA bases → Amino acids) represents a predefined mapping function. Ambiguous RNA tokens not explicitly mapped (e.g., X, <CLS>, <PAD>, <MASK>) inherit indices directly from the protein vocabulary when present, ensuring semantic alignment across domains. This hybrid approach generates pseudo-protein token sequences 𝑆^′^ = (𝑝_1_, 𝑝_2_, . . ., 𝑝_𝑛_) where 𝑝_𝑖_ corresponds to a mapped RNA token. Given the 116,280 possible one-to-one mappings from four RNA bases to 20 amino acids, we systematically tested both random and biophysically inspired mappings, including those from Jian et al.^49^ as well as our own biophysical mapping and random mapping. As a baseline control, we also evaluated a direct character mapping, AUCG mapped to AUCG. To assess the interaction between mapping strategies and model initialization, we conducted experiments both with and without loading pretrained weights from ProtESM-2 in the case of AUCG character-to-character mapping.

### Embedding layer

The pseudo-protein tokens 𝑆^′^ are projected into a shared embedding space using a trainable embedding layer. Each token 𝑝_𝑖_ is mapped to a 𝐷-dimensional vector ℎ_𝑖_ ∈ 𝑅^𝐷^defined as:

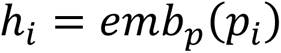

where 𝑝𝑒𝑏_𝑝_ denotes the protein embedding lookup operation. By leveraging RNA-to-protein mapping, RNA sequences inherit latent representations from protein LMs, enabling knowledge transfer of evolutionary and structural patterns. These embeddings are processed by a unified transformer architecture, which integrates cross-domain features to predict RNA secondary structures and functional tasks.

### Transformer module

The model employs a transformer architecture with pre-trained ESM-2-encoder blocks to process RNA sequence embeddings. Each encoder block consists of a multi-head self-attention mechanism followed by a multi-layer perceptron (MLP), with layer normalization applied before each sub-layer and residual connections added after. The self-attention mechanism dynamically weights pairwise interactions between all positions in the sequence, enabling the model to capture base-pairing dependencies and long-range interactions characteristics of RNA structures. The transformer processes input embeddings iteratively through stacked layers:

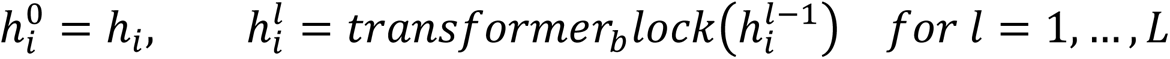

where 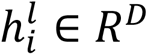 represents the hidden state of the 𝑖-th token at layer 𝑙. The final layer outputs encode contextualized features integrating local and global sequence constraints.

During pre-training, a masked language modeling head projects the output embeddings to predict masked pseudo-protein tokens, leveraging cross-entropy loss to align RNA sequences with protein-like representations. In the fine-tuning phase, the MLP or convolutional layers process the complete output embeddings to predict structure or functional tasks.

### Model pretraining Pretraining objective

The model was pre-trained using a masked language modeling objective, where 15% of pseudo-protein tokens in each RNA sequence were randomly masked. The training loss was defined as the cross-entropy between predicted token probabilities and original tokens, computed exclusively over non-padding positions:

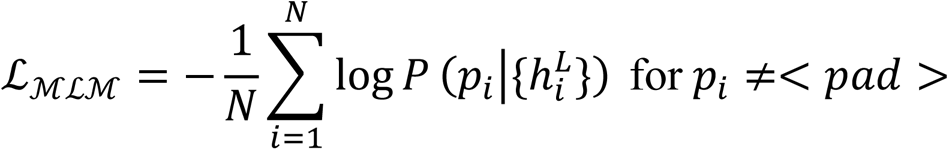

where 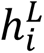 denotes the final-layer transformer embeddings, and 𝑃 is the softmax probability distribution over the protein vocabulary.

### Length-Aware Dynamic Batching Module

To optimize memory usage and computational efficiency during pretraining, we designed a length-aware dynamic batching module tailored for variable-length ncRNA sequences (Fig. 1b). Instead of using a fixed batch-size based on sequence counts, batches are constructed such that the cumulative number of tokens per batch does not exceed a predefined maximum token threshold.

The batching process begins by sorting all RNA sequences by length. The sorted dataset is then partitioned into length-stratified bins. During batch construction, sequences are dynamically sampled from these bins and grouped into batches, ensuring that the total number of tokens in each batch remains within the specified limit. This approach minimizes the amount of padding required within each batch, thereby reducing computational waste and improving GPU memory utilization.

### Structure-Aware Evaluation Module

To monitor model behavior during pretraining, we introduced a structure-aware diagnostic module that evaluates the model’s capacity to capture base-pair information in an unsupervised setting as illustrated in Fig. 1c. Specifically, self-attention maps were extracted from all heads and layers of the pre-trained RESM model and used as input features to a lightweight logistic regression classifier. This classifier was trained to predict RNA secondary structure labels.

Model performance was tracked using F1 scores on a validation set consisting of 40 RNA sequences (VL40) with experimentally determined three-dimensional structures. These sequences were drawn from the RNA-MSM benchmark dataset and filtered to ensure non-redundancy. This evaluation strategy allows for early insight into the model’s ability to encode biologically meaningful structural information, providing a low-cost, efficient proxy for structural awareness during pretraining.

To prepare attention maps for classification, self-attention matrices from intermediate transformer layers were first symmetrized as:

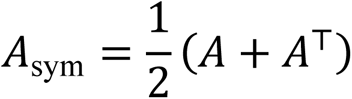

and then normalized via average product correction:

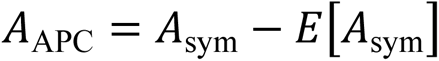

and finally employed as input features for a logistic regression classifier to predict base-pairing contacts.

For pretraining, we utilized the AdamW optimizer with a weight decay of 0.0001 and a warmup linear learning rate scheduler. The RESM-150M model was trained on 8 NVIDIA Tesla A100 GPUs, while RESM-650M scaled to 256 Huawei D910B GPUs to accommodate increased parameter complexity. Checkpoint mechanism is enabled by the training platform to guarantee the training continuity.

### Zero-shot RNA functional classification

We followed Wang et al.^29^ for zero-shot RNA functional classification. Briefly, we employed the Rfam 14.10 of 24,607 sequences in 4,170 families with up to 10 sequences randomly chosen from each family and ArchiveII datasets of 3,864 sequences in 9 RNA types after excluding the sequences longer than 510 nucleotides to facilitate the comparison to those RNA LMs with such limitation. The output embeddings for all sequences in both datasets were extracted and the cosine similarity values were calculated between every pair of homologous and non-homologous sequences. Two cosine similarity distributions for the same-family (RNA type) and different family (RNA type) groups were obtained, and the overlap ratio (OR) between the two distributions was computed to quantify the model’s ability to distinguish between homologous and non-homologous sequences with 0 for zero overlap (complete separation) and 1 for complete overlap (completely indistinguishable). And to ensure robustness, three independent samplings were performed on both the Rfam and ArchiveII datasets when calculating OR, where 100,000 same-family (RNA type) sequence pairs and 100,000 different family (RNA type) were randomly selected for each sampling, and the average of the ORs were shown.

### Zero-shot secondary structure prediction

We also followed Wang et al.^29^ for zero-shot RNA secondary structure prediction. Briefly, each attention map was processed using a series of transformations, including symmetrization, APC correction, and sigmoid function scaling to predict the probability of base-pair formation. For every attention map, a threshold value within the range of [0, 1] was tested at intervals of 0.001. The corresponding F1 score for secondary structure prediction was calculated for each attention map, and the optimal head-layer position and threshold were selected based on their performance on the validation dataset. These parameters were then applied to the independent test set to directly evaluate the model’s zero-shot performance without training. We utilized the validation set VL40 and test set TS70 derived from 3D structures employed in RNA-MSM for this zero-shot prediction. The two sets were ensured non-redundancy by using sequence identity cut off of 80% from CD-HIT-EST and structural similarity cut off at 0.45 according to a TM-score.

### Supervised secondary structure prediction by RESM-basepair

A network architecture and training strategy of SPOT-RNA2 was employed for supervised RNA secondary structure prediction. The model is an ensemble of five deep neural networks, each consisting of ResNet^70^ blocks followed by a bidirectional 2D-LSTM layer and a fully connected (FC) block (Supplementary Fig. 1), the detailed configuration of which can be found in Supplementary Table 16. To avoid overfitting during training, dropout rates of 25% and 50% were utilized in each ResNet block and fully connected (FC) block, respectively. When dilated convolutional layers were used, the dilation factor was set to 2*^i^*^%*n*^, where *i* is the depth of the convolution layer, *n* is a fixed scalar, and % is the modulus operator. The input features include both the one-hot encoding used in SPOT-RNA and the attention maps generated by RESM. The attention tensors have dimensions 𝑟 × ℎ × 𝐿 × 𝐿, where 𝑟 is the number of layers, ℎ is the number of attention heads, and 𝐿 is the sequence length. Prior to integration, the attention maps were corrected using average product correction (APC) and symmetrized to reduce background noise.

The training and evaluation datasets for supervised finetuning during pre-training process were TR0 and VL0, respectively, which were derived from SPOT-RNA and contained 10,814 and 1,300 RNAs extracted from the bpRNA-1m database, respectively. Each of the five deep neural networks was first pre-trained on the TR0 dataset using the AdamW optimizer and an initial learning rate of 5e-4, and the best-performing checkpoints were selected based on validation performance on VL0. These networks are further trained on the TR dataset with VL40 used as the validation set and TS70 as the final test set for evaluation. During the transfer learning stage, all the weights and parameters learned on TR0 are retrained on TR with an initial learning rate of 1e-4.

### Zero-shot RNA-RNA interaction prediction by RESM-basepair

RNA–RNA interaction predictions were evaluated using the benchmark introduced by Lang et al.^56^, which comprises 64 unique interaction pairs derived from experimentally resolved three-dimensional RNA complex structures by excluding sequentially and structurally similar RNAs to SPOT-RNA training sets. Each RNA pair was concatenated into a single sequence, and secondary structure prediction was performed using RESM-basepair-150M and RESM-basepair-650M (Supplementary Fig. 1). Only the predicted base pairs that spanned the two RNA regions were extracted and considered for evaluation. To account for potential chain order bias, predictions were made for both concatenation orders (AB and BA), and an interaction was counted as positive if it was detected in either direction. We conducted experiments with and without the use of synthetic polynucleotide linkers between RNA pairs. Among the tested configurations, the use of 𝑋^8^ and 𝑋^2^ linkers yield the best overall performance for RESM-basepair-150M and RESM-basepair-650M, respectively. Predicting RNA-RNA interaction is considered as zero-shot prediction as the methods tested were not trained for RNA-RNA interactions.

### Supervised RNA solvent accessibility prediction experiments by RESM-ASA

We constructed the solvent accessibility prediction model by integrating a simple 2-layer MLP with the pre-trained model, instead of using the more complex architecture employed in RNA-MSM (Supplementary Fig. 2). The final-layer embeddings from the pre-trained model were used as input to the MLP, which predicted RSA values within the range of 0 to 1. These predicted RSA values were then converted to absolute ASA values using the normalization factors described above with the mean absolute error (MAE) between the predicted and actual RSA values serving as the loss function. To evaluate the model’s ability to infer nucleotide-level solvent exposure, we used the same non-redundant dataset split as described in the zero-shot secondary structure prediction task, ensuring no sequence or structural redundancy across the training (TR405), validation (VL40), and test (TS70) sets. Solvent-accessible surface area (ASA) values were computed from 3D RNA chain structures using the POPS algorithm with a solvent probe radius of 1.4 Å. These values were normalized to relative solvent accessibility (RSA) by dividing each nucleotide’s ASA by its theoretical maximum exposure (adenine and guanine: 400 Å²; uracil and cytosine: 350 Å²), a normalization strategy adapted from prior work^34^ to account for nucleotide-specific size variations.

### Supervised siRNA activity prediction experiments by RESM-siRNA

We adopted the architecture and dataset curation strategy of OligoFormer while replacing RNA-FM with RESM as the foundational embedding generator (Supplementary Fig. 3). The Huesken dataset (2,361 samples) and Mixset (472 samples) were preprocessed using Needleman–Wunsch global alignment to remove sequences with >80% identity, ensuring dataset independence. For inter-dataset validation, models were trained on Huesken and evaluated on Mixset to assess cross-condition robustness.

The model framework comprises three parallel modules. The first module is the RESM Embedding Module, where siRNA sequences (19-nt) and flanking mRNA regions (57-nt centered on target sites) are encoded into latent space via RESM. The second module, the Oligo Encoder, processes these embeddings through independent bidirectional encoders for siRNA and mRNA sequences. The Oligo Encoder is built with a hybrid architecture that includes a 2D convolutional layer to capture spatial dependencies in the embedding matrices, followed by max and average pooling to retain salient features while reducing dimensionality. A BiLSTM layer models the sequential dynamics across nucleotide positions, while a multi-head transformer attends to long-range siRNA-mRNA interaction patterns. Finally, the extracted features are consolidated into a single representation via a flatten layer, producing the final embeddings used for the subsequent prediction task.

For feature integration, the outputs from all modules are concatenated and passed through a multilayer perceptron (MLP) classifier. This classifier consists of two hidden layers with 512 and 128 units, respectively, using ReLU activation and a dropout rate of 0.3. The final output layer generates siRNA efficacy probabilities in the range of 0 to 1, representing the predicted efficacy of each siRNA.

### Supervised ribosome loading and expression levels prediction experiments by RESM-riboload and RESM-mRNAexpr

To further evaluate the generalization capability of the pre-trained model, we conducted experiments on two representative datasets corresponding to ribosome loading and mRNA expression level prediction. Both tasks were formulated as sequence-to-regression problems.

For ribosome loading prediction, we adopted the dataset introduced by Sample et al.^62^, which consists of synthetic 5’ untranslated region (UTR) sequences ranging from 25 to 100 nucleotides in length. A total of 76,319 randomized 5’UTRs were used for training. To rigorously evaluate generalization, two separate test sets were employed: one composed of 7,600 synthetic sequences and the other containing 7,600 human-derived 5’UTRs. To avoid potential data leakage, sequences with over 80% identity between the training and test sets were removed using CD-HIT-EST, and the final de-redundant training set consisted of 70,372 sequences. This ensured that model evaluation on the test sets reflected true generalization.

For mRNA expression level prediction, we used three publicly available datasets^67^ representing distinct biological contexts: human skeletal muscle tissue, prostate cancer (PC3) cells, and human embryonic kidney (HEK) 293T cells. Following prior work^30^, the input sequence for each sample was initially truncated to the first 100 nucleotides of the 5’UTR. However, we observed substantial sequence redundancy across samples due to repeated use of identical truncated sequences with varying expression values, with overlap rates of 46.0%, 47.8%, and 46.7% in the muscle, PC3, and HEK datasets, respectively. To mitigate this, we also evaluated model performance using full-length RNA sequences, enabling the model to access additional regulatory information beyond the fixed-length window.

In both tasks, inputs consisted of one-hot encoded RNA sequences combined with final-layer embeddings from the pre-trained RESM model. The regression targets were ribosome loading efficiency or normalized mRNA expression levels.

We fine-tuned the pre-trained model, maintaining consistency with the downstream prediction head architecture, ResNet1D, used in mRNA regulatory task prediction to demonstrate the generalizability of the pre-trained model across downstream tasks, even when applied to out-of-domain datasets (Supplementary Fig. 4). Utilizing the same downstream network structure for both ribosome loading and expression level prediction ensured that performance improvements stemmed from the model’s inherent adaptability rather than task-specific architectural optimization.

For fine-tuning, we used the pre-trained RESM model with the ResNet1D prediction head. The ResNet1D prediction head comprises six residual blocks, each containing two 1D convolutional layers (kernel size=3) followed by batch normalization and ReLU activation. Final-layer transformer embeddings, along with the one-hot encoding of the input sequence, were first projected into a 64-dimensional space via linear transformation. A 3×3 1D convolutional layer then reduced the feature dimensionality to 32 channels, which were processed sequentially through the residual blocks. The output was pooled globally and passed to a linear layer for regression.

The entire model, consisting of the pre-trained RESM and the downstream ResNet1D prediction head, was trained using the Mean Squared Error (MSE) loss function as the optimization objective. The learning rate for the pre-trained RESM was set to 0.00001, and the learning rate for the downstream ResNet1D prediction head was set to 0.0001.

### Evaluation metrics

Model performance was evaluated using task-specific metrics that reflect biological relevance. Instead of using perplexity, which is commonly adopted in natural language processing to quantify a model’s uncertainty in reconstructing masked tokens during pretraining, we employed the F1 score to better assess the model’s base pair prediction capability. In both zero-shot and supervised RNA secondary structure prediction tasks, the F1 score was utilized to evaluate base-pair prediction performance, capturing the harmonic mean of precision and recall in identifying paired nucleotides. The threshold for a defined base pair was evaluated on the VL40 dataset, where the value enabling the model to obtain the best F1 score on VL40 would be selected as the threshold on the test dataset. Functional classification of RNA families employed the overlap ratio (OR)^29^, defined as the intersection area of predicted and ground-truth probability distributions divided by their union area, with values bounded in [0, 1]; a lower OR value indicates a better separation of different classifications with OR=0 for 0 overlap and 1 for 100% overlap. Here, the predicted probability distribution is the cosine similarity distribution of same-family (RNA type) sequence pairs, while the ground-truth probability distributions is the cosine similarity distribution of different family (RNA type) sequence pairs. The cosine similarity of two sequences in a sample sequence pair was defined as:

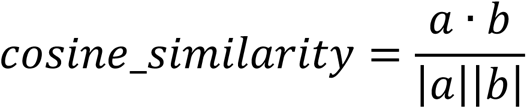

Where *a* and *b* were the representational vectors of sequence A and B, respectively; |𝑥| denotes the modulo operation; the cosine similarity has the value in the range [0, 1].

RNA-RNA interaction prediction was evaluated using multiple metrics, including precision, recall (sensitivity), F1 score (overall and per-RNA), and Matthews correlation coefficient (MCC), offering a comprehensive view of binary interaction detection performance. As with the RNA secondary structure prediction task, the threshold for a defined base pair was evaluated on the VL40 dataset. The median and standard deviation of individual F1 scores were also reported to capture variability across RNA pairs. RNA solvent-accessibility predictions utilized Pearson’s correlation coefficient (PCC) to measure linear agreement and mean absolute error (MAE) to quantify absolute deviation, with higher PCC and lower MAE indicating superior performance. Ribosome loading and gene expression level predictions were assessed using Spearman’s rank correlation coefficient, which evaluates monotonic relationships between predicted and observed values. For siRNA activity prediction, model robustness was evaluated through the areas under the receiver operating characteristic curve (AUC_ROC) and precision-recall curve (AUC_PRC) as well as F1 scores, capturing both class imbalance resilience and discriminative power.

## Supporting information

https://github.com/yikunpku/RESM

## Code availability

The source code for the RESM model and inference scripts are freely available at GitHub (https://github.com/yikunpku/RESM) under an MIT license.

## Data availability

All data used in this study are publicly available.

For pre-training RESM, we obtained 23.51 million non-redundant non-coding RNA sequences from RNAcentral (https://rnacentral.org/). The datasets for downstream tasks were obtained as follows: RNA functional classification data from Rfam (https://rfam.xfam.org) and ArchiveII (https://huggingface.co/datasets/multimolecule/archiveii); RNA secondary structure prediction data from the RNA-MSM dataset (https://github.com/yikunpku/RNA-MSM) and bpRNA-1m (https://bprna.cgrb.oregonstate.edu/); RNA–RNA interaction data from https://github.com/meilanglang/RNA-RNA-Interaction; siRNA activity prediction data (Huesken and Mixset datasets) from https://github.com/lulab/OligoFormer; ribosome loading data (GSM4084997) from Gene Expression Omnibus (https://www.ncbi.nlm.nih.gov/geo/query/acc.cgi?acc=GSE114002); and mRNA expression data (muscle, HEK293T, and PC3 datasets) from https://github.com/a96123155/UTR-LM. Detailed statistics for these datasets are provided in Supplementary Table 1. All datasets used in this study, including the RESM pre-training dataset, benchmark datasets for RNA downstream tasks, and trained weights (RESM-150M and RESM-650M), are publicly available at Zenodo (https://doi.org/10.5281/zenodo.15980875).

## Acknowledgments

This research received support from the National Natural Science Foundation of China (Grant # 22350710182) and benefited from access to the supercomputing resources at China Mobile, Shenzhen Bay Laboratory and Shenzhen Medical Academy of Research and Translation.

## Author contributions

YiZ developed the neural network architecture and training for the pretraining model. HZ, GL, TZ, & LW developed and supported the model training on the Huawei Ascend computing platform. YiZ & HW collected and processed the data for the downstream tasks, developed the downstream training and inference systems, and analyzed and interpreted the experimental results. XZ, XH, JJ, JC, JZ, & YaZ contributed technical advice and ideas. JC, YC, LL, JZ & YaZ co-supervised the project. YaZ and JZ initiated and provided the funding support for the project. YiZ, HW and YaZ wrote the initial draft. All authors were involved in subsequent manuscript improvement and approved the final manuscript.

## Supplementary Information

### Data details

**Supplementary Table 1.**
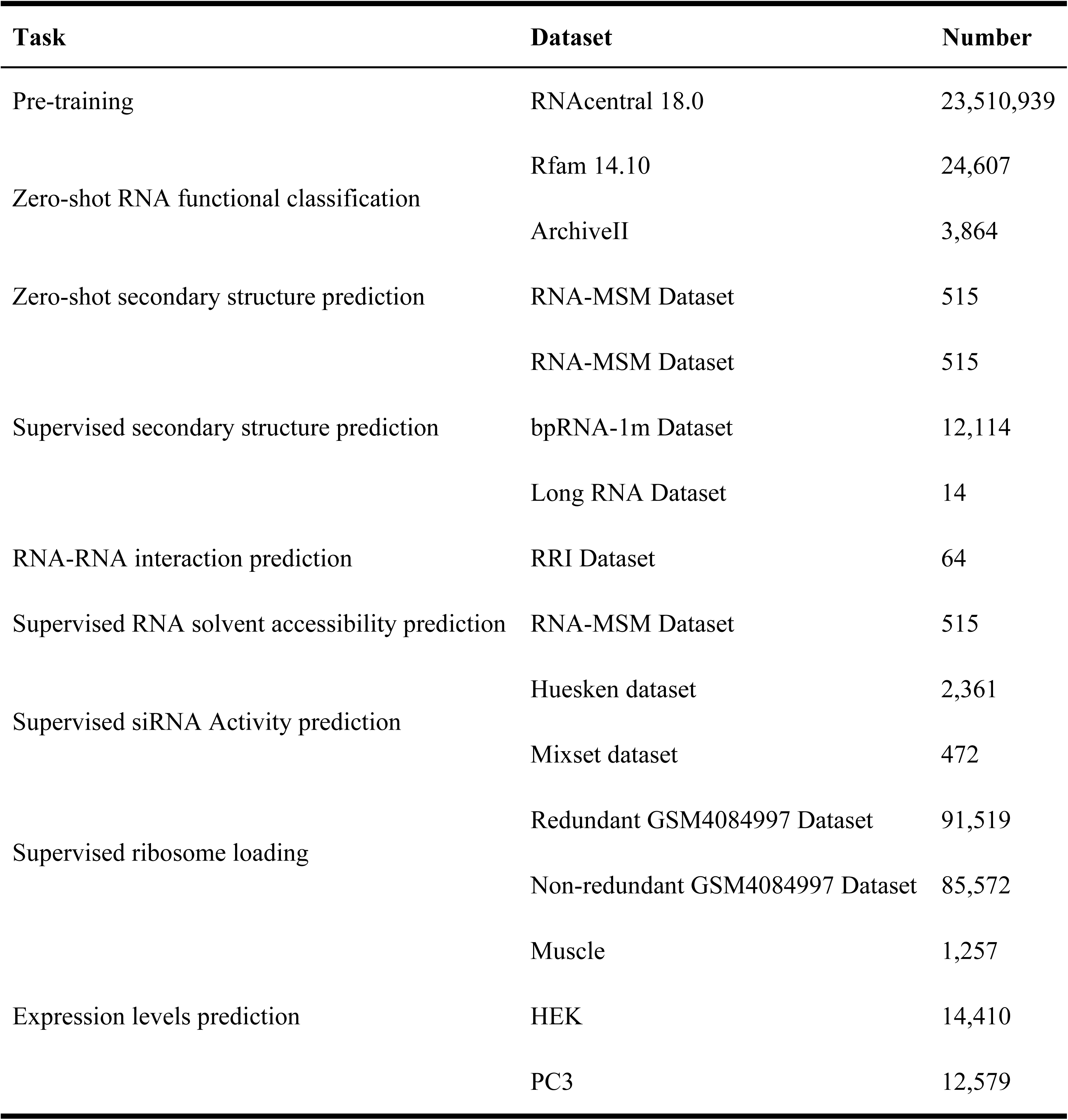
Datasets used for RESM pre-training and downstream tasks. Overview of datasets and sample sizes across all tasks. RESM was pretrained on RNAcentral and evaluated on zero-shot and supervised benchmarks covering structure, function, interaction, and regulatory prediction tasks.

### Results

**Supplementary Table 2.**
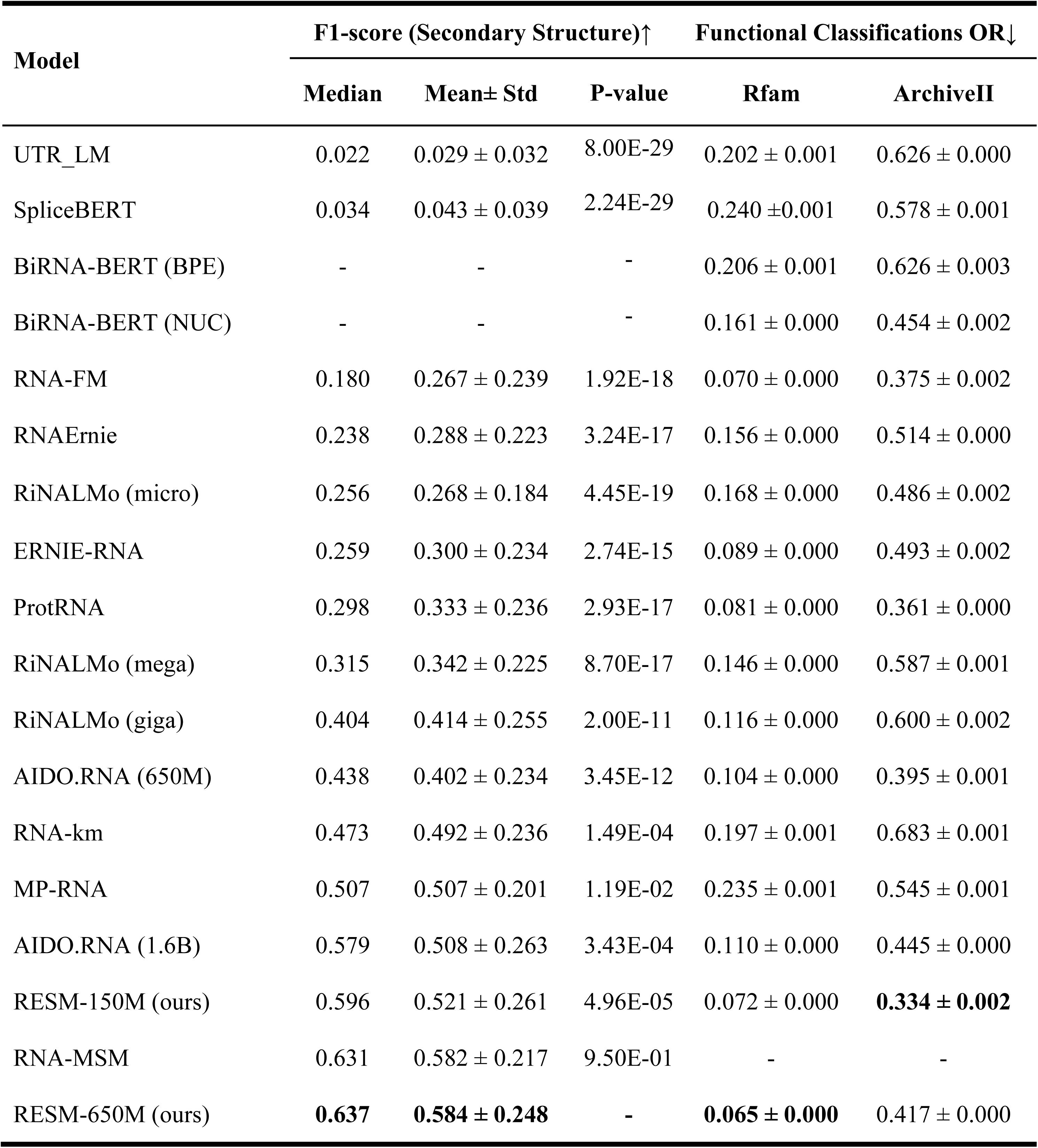
Zero-shot evaluation on RNA secondary structure and functional classification tasks. Models were evaluated under a unified zero-shot setting using attention maps from a fixed head-layer across all RNA sequences. For secondary-structure-prediction evaluation, median and mean ± standard deviation of individual overall F1 scores are reported to capture overall predictive quality and consistency. Functional classification accuracy is assessed using the overlap ratio (OR) on two datasets, Rfam and ArchiveII, with lower values indicating better class separability.

| Model | F1-score (Secondary Structure)↑ | | | Functional Classifications OR↓ | |
| --- | --- | --- | --- | --- | --- |
| | Median | Mean± Std | P-value | Rfam | ArchiveII |
| UTR_LM | 0.022 | 0.029 ± 0.032 | 8.00E-29 | 0.202 ± 0.001 | 0.626 ± 0.000 |
| SpliceBERT | 0.034 | 0.043 ± 0.039 | 2.24E-29 | 0.240 ± 0.001 | 0.578 ± 0.001 |
| BiRNA-BERT (BPE) | - | - | - | 0.206 ± 0.001 | 0.626 ± 0.003 |
| BiRNA-BERT (NUC) | - | - | - | 0.161 ± 0.000 | 0.454 ± 0.002 |
| RNA-FM | 0.180 | 0.267 ± 0.239 | 1.92E-18 | 0.070 ± 0.000 | 0.375 ± 0.002 |
| RNAErnie | 0.238 | 0.288 ± 0.223 | 3.24E-17 | 0.156 ± 0.000 | 0.514 ± 0.000 |
| RiNALMo (micro) | 0.256 | 0.268 ± 0.184 | 4.45E-19 | 0.168 ± 0.000 | 0.486 ± 0.002 |
| ERNIE-RNA | 0.259 | 0.300 ± 0.234 | 2.74E-15 | 0.089 ± 0.000 | 0.493 ± 0.002 |
| ProtRNA | 0.298 | 0.333 ± 0.236 | 2.93E-17 | 0.081 ± 0.000 | 0.361 ± 0.000 |
| RiNALMo (mega) | 0.315 | 0.342 ± 0.225 | 8.70E-17 | 0.146 ± 0.000 | 0.587 ± 0.001 |
| RiNALMo (giga) | 0.404 | 0.414 ± 0.255 | 2.00E-11 | 0.116 ± 0.000 | 0.600 ± 0.002 |
| AIDO.RNA (650M) | 0.438 | 0.402 ± 0.234 | 3.45E-12 | 0.104 ± 0.000 | 0.395 ± 0.001 |
| RNA-km | 0.473 | 0.492 ± 0.236 | 1.49E-04 | 0.197 ± 0.001 | 0.683 ± 0.001 |
| MP-RNA | 0.507 | 0.507 ± 0.201 | 1.19E-02 | 0.235 ± 0.001 | 0.545 ± 0.001 |
| AIDO.RNA (1.6B) | 0.579 | 0.508 ± 0.263 | 3.43E-04 | 0.110 ± 0.000 | 0.445 ± 0.000 |
| RESM-150M (ours) | 0.596 | 0.521 ± 0.261 | 4.96E-05 | 0.072 ± 0.000 | **0.334 ± 0.002** |
| RNA-MSM | 0.631 | 0.582 ± 0.217 | 9.50E-01 | - | - |
| RESM-650M (ours) | **0.637** | **0.584 ± 0.248** | - | **0.065 ± 0.000** | 0.417 ± 0.000 |

**Supplementary Table 3.**
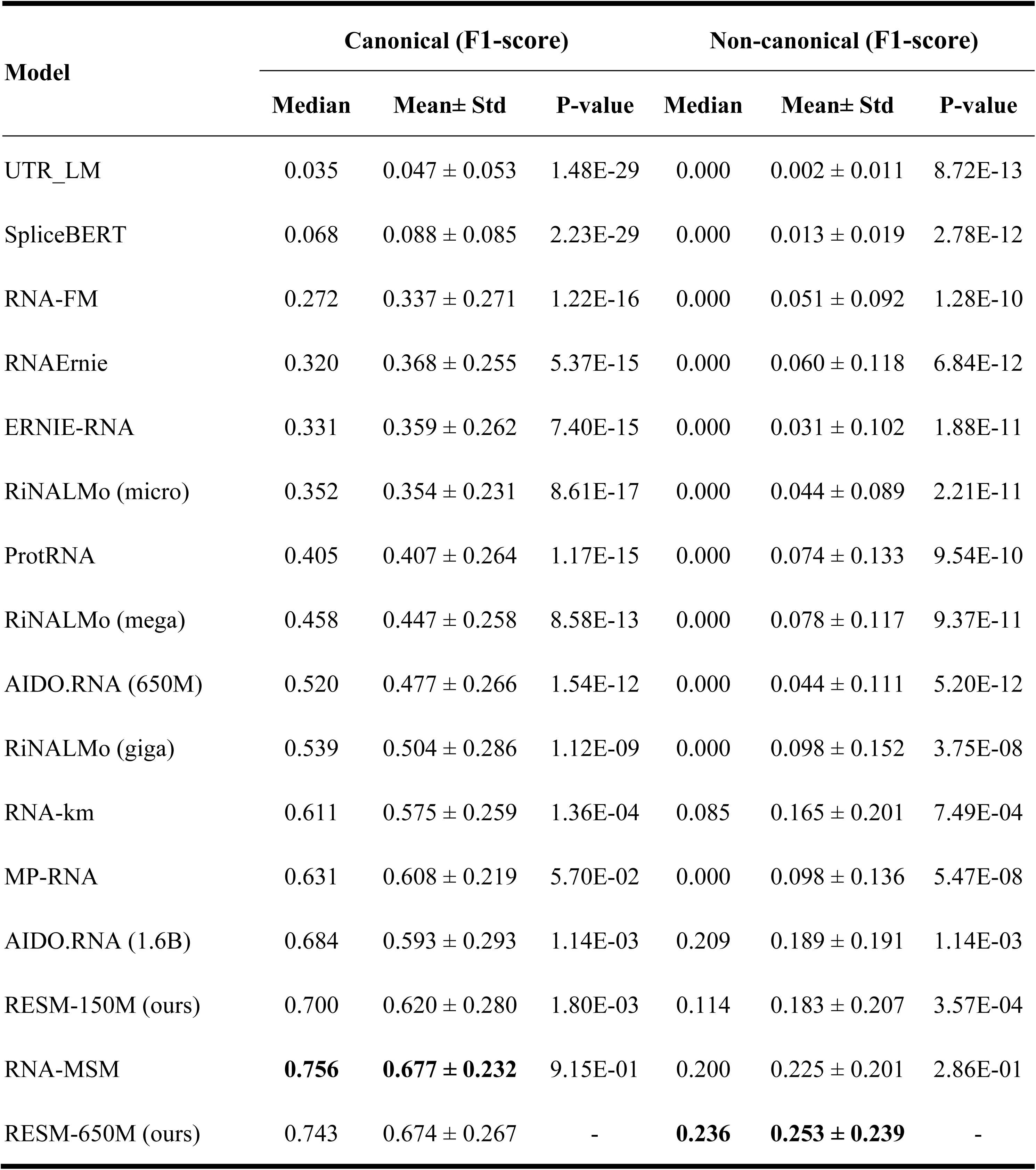
Zero-shot RNA secondary structure prediction. Performance on the TS70 test set, evaluated separately on canonical and non-canonical base pairs. The table reports the median and mean ± standard deviation of the F1 scores for each model, along with corresponding P-values from paired comparisons with RESM-650M in this work.

| Model | Canonical (F1-score) |  |  | Non-canonical (F1-score) |  |  |
| --- | --- | --- | --- | --- | --- | --- |
| | Median | Mean± Std | P-value | Median | Mean± Std | P-value |
| UTR_LM | 0.035 | 0.047 ± 0.053 | 1.48E-29 | 0.000 | 0.002 ± 0.011 | 8.72E-13 |
| SpliceBERT | 0.068 | 0.088 ± 0.085 | 2.23E-29 | 0.000 | 0.013 ± 0.019 | 2.78E-12 |
| RNA-FM | 0.272 | 0.337 ± 0.271 | 1.22E-16 | 0.000 | 0.051 ± 0.092 | 1.28E-10 |
| RNAErnie | 0.320 | 0.368 ± 0.255 | 5.37E-15 | 0.000 | 0.060 ± 0.118 | 6.84E-12 |
| ERNIE-RNA | 0.331 | 0.359 ± 0.262 | 7.40E-15 | 0.000 | 0.031 ± 0.102 | 1.88E-11 |
| RiNALMo (micro) | 0.352 | 0.354 ± 0.231 | 8.61E-17 | 0.000 | 0.044 ± 0.089 | 2.21E-11 |
| ProtRNA | 0.405 | 0.407 ± 0.264 | 1.17E-15 | 0.000 | 0.074 ± 0.133 | 9.54E-10 |
| RiNALMo (mega) | 0.458 | 0.447 ± 0.258 | 8.58E-13 | 0.000 | 0.078 ± 0.117 | 9.37E-11 |
| AIDO.RNA (650M) | 0.520 | 0.477 ± 0.266 | 1.54E-12 | 0.000 | 0.044 ± 0.111 | 5.20E-12 |
| RiNALMo (giga) | 0.539 | 0.504 ± 0.286 | 1.12E-09 | 0.000 | 0.098 ± 0.152 | 3.75E-08 |
| RNA-km | 0.611 | 0.575 ± 0.259 | 1.36E-04 | 0.085 | 0.165 ± 0.201 | 7.49E-04 |
| MP-RNA | 0.631 | 0.608 ± 0.219 | 5.70E-02 | 0.000 | 0.098 ± 0.136 | 5.47E-08 |
| AIDO.RNA (1.6B) | 0.684 | 0.593 ± 0.293 | 1.14E-03 | 0.209 | 0.189 ± 0.191 | 1.14E-03 |
| RESM-150M (ours) | 0.700 | 0.620 ± 0.280 | 1.80E-03 | 0.114 | 0.183 ± 0.207 | 3.57E-04 |
| RNA-MSM | **0.756** | **0.677 ± 0.232** | 9.15E-01 | 0.200 | 0.225 ± 0.201 | 2.86E-01 |
| RESM-650M (ours) | 0.743 | 0.674 ± 0.267 | - | **0.236** | **0.253 ± 0.239** | - |

**Supplementary Table 4.** Performance of supervised RNA secondary-structure prediction using attention-based representations on the TS70 dataset, as compared to other models, according to the F1 score. . Supervised prediction was performed using attention maps aggregated from all layers and attention heads of each model as input to a downstream task-specific network. Model performance was evaluated across the dataset in terms of median F1 score and mean ± standard deviation of individual F1 scores for each RNA. P-values indicate the statistical significance of paired comparisons with RESM-basepair-650M.

| Model | F1-score |  | P-value |
| --- | --- | --- | --- |
| | Median | Mean ± Std | |
| $\wedge$Rfold | 0.252 | 0.326 ± 0.275 | 2.54E-16 |
| *ProtRNA | 0.372 | 0.382 ± 0.265 | 5.29E-17 |
| *ERNIE-RNA | 0.386 | 0.428 ± 0.281 | 4.31E-13 |
| $\wedge$RNADIFFfold | 0.432 | 0.469 ± 0.242 | 7.34E-12 |
| *RiNALMo (giga) | 0.533 | 0.506 ± 0.246 | 5.03E-12 |
| *AIDO.RNA (1.6B) | 0.580 | 0.523 ± 0.242 | 6.13E-11 |
| RESM-zero-shot-150M (ours) | 0.596 | 0.521 ± 0.261 | 1.80E-12 |
| *RNAErnie | 0.601 | 0.546 ± 0.260 | 3.77E-06 |
| †RNAstructure | 0.606 | 0.571 ± 0.271 | 4.99E-04 |
| †CONTRAFold | 0.628 | 0.576 ± 0.265 | 3.99E-04 |
| RESM-zero-shot-650M (ours) | 0.637 | 0.584 ± 0.248 | 5.27E-08 |
| †EternaFold | 0.638 | 0.592 ± 0.245 | 1.15E-03 |
| $\wedge$KnotFold | 0.651 | 0.568 ± 0.248 | 1.31E-06 |
| †LinearFold | 0.653 | 0.580 ± 0.273 | 1.09E-03 |
| †RNAfold | 0.656 | 0.586 ± 0.273 | 3.27E-03 |
| <u>$\wedge$BPfold</u> | 0.667 | 0.628 ± 0.207 | 5.68E-03 |
| <u>^UFold</u> | 0.667 | $0.606 \pm 0.223$ | 6.56E-04 |
| *RNA-FM | 0.679 | $0.634 \pm 0.242$ | 1.34E-02 |
| <u>^MXfold2</u> | 0.696 | $0.648 \pm 0.217$ | 7.99E-02 |
| <u>^SPOT-RNA</u> | 0.699 | $0.629 \pm 0.231$ | 5.56E-03 |
| *RNA-MSM | 0.721 | $0.662 \pm 0.211$ | 1.49E-01 |
| <u>^SPOT-RNA2</u> | 0.743 | $0.678 \pm 0.211$ | 5.36E-01 |
| RESM-basepair-650M (ours) | 0.746 | **$0.701 \pm 0.180$** | - |
| RESM-basepair-150M (ours) | **0.748** | $0.686 \pm 0.190$ | 2.20E-01 |

Note: Models with “†” denote thermodynamic or probabilistic methods; models with “^” denote deep learning methods; models with “\*” denote RNA language models with trained RNA secondary-structure prediction header, whose prediction leader networks and training weights are available from the original reported literature. “RESM-zero-shot-150M” and “RESM-zero-shot-650M” show the zero-shot results, consistent with the results of “RESM-150M” and “RESM-650M” in Supplementary Table 2, respectively. Models with an underlined name have training set overlap with TS70. And limited by the sequence requirements of Rfold, Ufold and SPOT-RNA2 methods, they only show the results of 66, 66, and 67 PDBs from the TS70 dataset, respectively.

**Supplementary Table 5.**
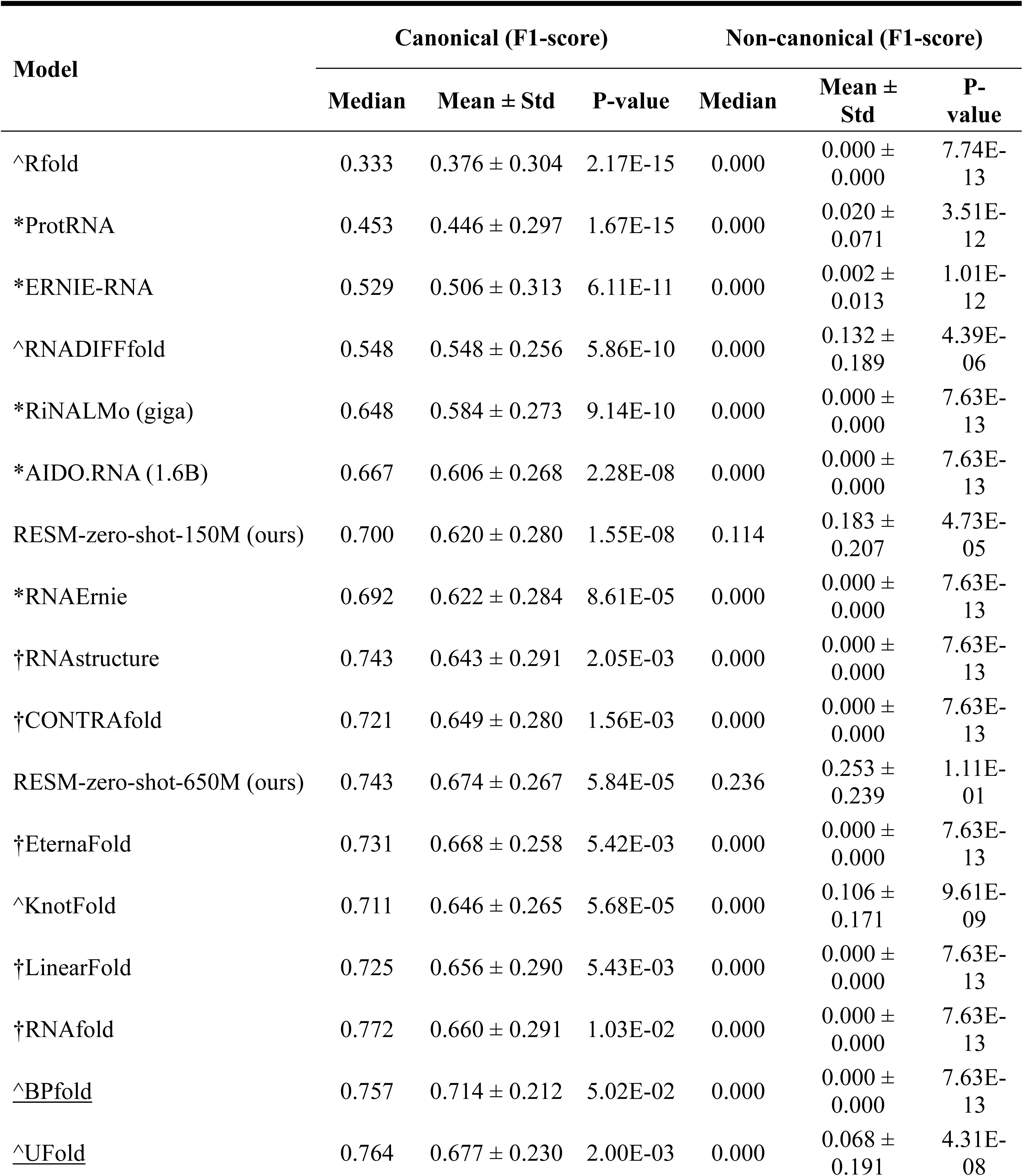
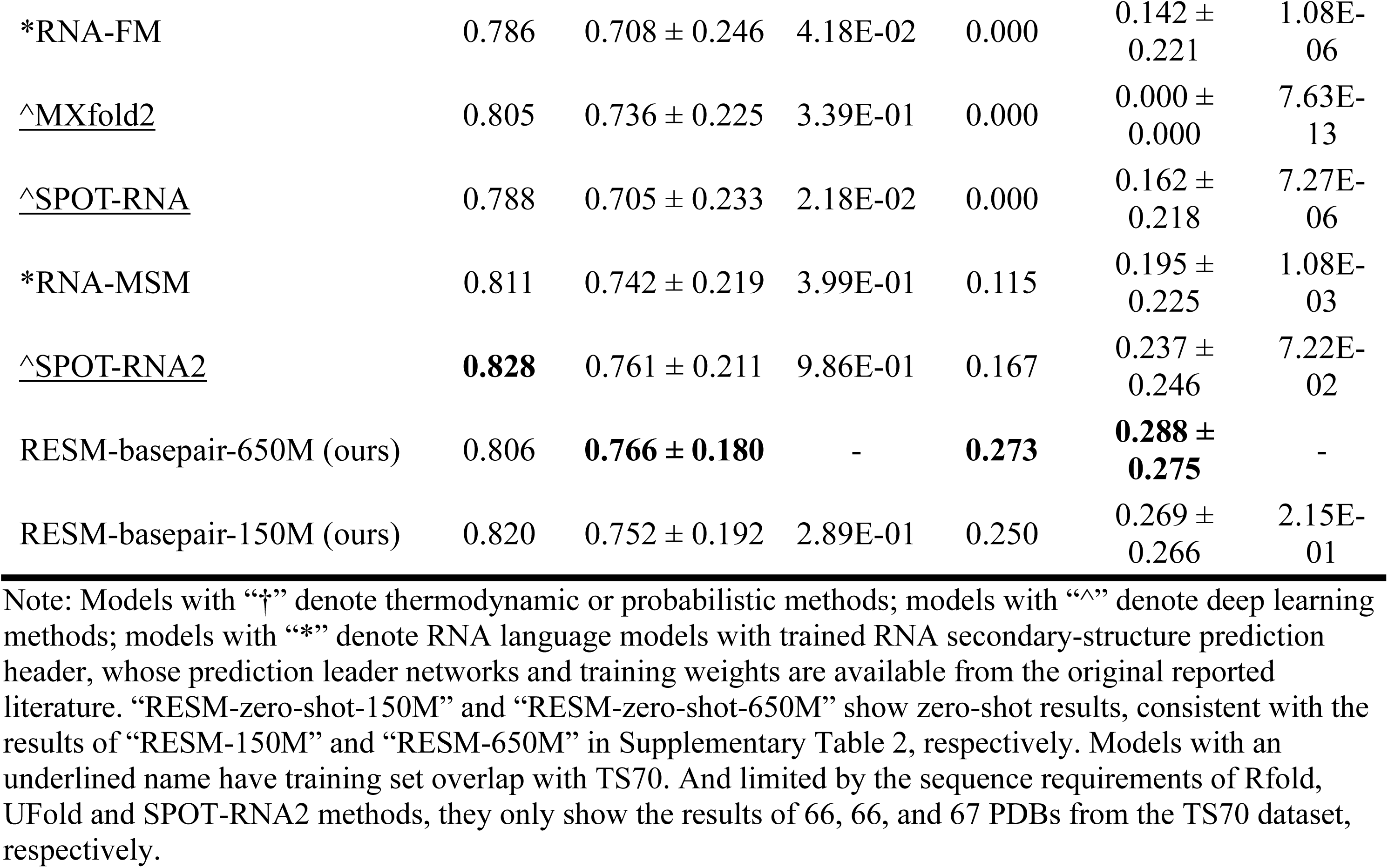
Performance of supervised RNA secondary-structure prediction using attention-based representations on the TS70 dataset, as compared to other models, according to canonical and non-canonical base pairs, separately. Results are reported separately for canonical and non-canonical base pairs. F1 scores are presented as the median and the mean ± standard deviation. P-values indicate the statistical significance of paired comparisons with RESM-basepair-650M.

| Model | Canonical (F1-score) |  |  | Non-canonical (F1-score) |  |  |
| --- | --- | --- | --- | --- | --- | --- |
| | Median | Mean ± Std | P-value | Median | Mean ± Std | P-value |
| $\wedge$Rfold | 0.333 | 0.376 ± 0.304 | 2.17E-15 | 0.000 | 0.000 ± 0.000 | 7.74E-13 |
| *ProtRNA | 0.453 | 0.446 ± 0.297 | 1.67E-15 | 0.000 | 0.020 ± 0.071 | 3.51E-12 |
| *ERNIE-RNA | 0.529 | 0.506 ± 0.313 | 6.11E-11 | 0.000 | 0.002 ± 0.013 | 1.01E-12 |
| $\wedge$RNADIFFfold | 0.548 | 0.548 ± 0.256 | 5.86E-10 | 0.000 | 0.132 ± 0.189 | 4.39E-06 |
| *RiNALMo (giga) | 0.648 | 0.584 ± 0.273 | 9.14E-10 | 0.000 | 0.000 ± 0.000 | 7.63E-13 |
| *AIDO.RNA (1.6B) | 0.667 | 0.606 ± 0.268 | 2.28E-08 | 0.000 | 0.000 ± 0.000 | 7.63E-13 |
| RESM-zero-shot-150M (ours) | 0.700 | 0.620 ± 0.280 | 1.55E-08 | 0.114 | 0.183 ± 0.207 | 4.73E-05 |
| *RNAErnie | 0.692 | 0.622 ± 0.284 | 8.61E-05 | 0.000 | 0.000 ± 0.000 | 7.63E-13 |
| †RNAstructure | 0.743 | 0.643 ± 0.291 | 2.05E-03 | 0.000 | 0.000 ± 0.000 | 7.63E-13 |
| †CONTRAFold | 0.721 | 0.649 ± 0.280 | 1.56E-03 | 0.000 | 0.000 ± 0.000 | 7.63E-13 |
| RESM-zero-shot-650M (ours) | 0.743 | 0.674 ± 0.267 | 5.84E-05 | 0.236 | 0.253 ± 0.239 | 1.11E-01 |
| †EternaFold | 0.731 | 0.668 ± 0.258 | 5.42E-03 | 0.000 | 0.000 ± 0.000 | 7.63E-13 |
| $\wedge$KnotFold | 0.711 | 0.646 ± 0.265 | 5.68E-05 | 0.000 | 0.106 ± 0.171 | 9.61E-09 |
| †LinearFold | 0.725 | 0.656 ± 0.290 | 5.43E-03 | 0.000 | 0.000 ± 0.000 | 7.63E-13 |
| †RNAfold | 0.772 | 0.660 ± 0.291 | 1.03E-02 | 0.000 | 0.000 ± 0.000 | 7.63E-13 |
| <u>$\wedge$BPfold</u> | 0.757 | 0.714 ± 0.212 | 5.02E-02 | 0.000 | 0.000 ± 0.000 | 7.63E-13 |
| <u>$\wedge$UFold</u> | 0.764 | 0.677 ± 0.230 | 2.00E-03 | 0.000 | 0.068 ± 0.191 | 4.31E-08 |
| *RNA-FM | 0.786 | 0.708 ± 0.246 | 4.18E-02 | 0.000 | 0.142 ± 0.221 | 1.08E-06 |
| <u>^MXfold2</u> | 0.805 | 0.736 ± 0.225 | 3.39E-01 | 0.000 | 0.000 ± 0.000 | 7.63E-13 |
| <u>^SPOT-RNA</u> | 0.788 | 0.705 ± 0.233 | 2.18E-02 | 0.000 | 0.162 ± 0.218 | 7.27E-06 |
| *RNA-MSM | 0.811 | 0.742 ± 0.219 | 3.99E-01 | 0.115 | 0.195 ± 0.225 | 1.08E-03 |
| <u>^SPOT-RNA2</u> | **0.828** | 0.761 ± 0.211 | 9.86E-01 | 0.167 | 0.237 ± 0.246 | 7.22E-02 |
| RESM-basepair-650M (ours) | 0.806 | **0.766 ± 0.180** | - | **0.273** | **0.288 ± 0.275** | - |
| RESM-basepair-150M (ours) | 0.820 | 0.752 ± 0.192 | 2.89E-01 | 0.250 | 0.269 ± 0.266 | 2.15E-01 |

Note: Models with “†” denote thermodynamic or probabilistic methods; models with “^” denote deep learning methods; models with “\*” denote RNA language models with trained RNA secondary-structure prediction header, whose prediction leader networks and training weights are available from the original reported literature. “RESM-zero-shot-150M” and “RESM-zero-shot-650M” show zero-shot results, consistent with the results of “RESM-150M” and “RESM-650M” in Supplementary Table 2, respectively. Models with an underlined name have training set overlap with TS70. And limited by the sequence requirements of Rfold, Ufold and SPOT-RNA2 methods, they only show the results of 66, 66, and 67 PDBs from the TS70 dataset, respectively.

**Supplementary Table 6.**
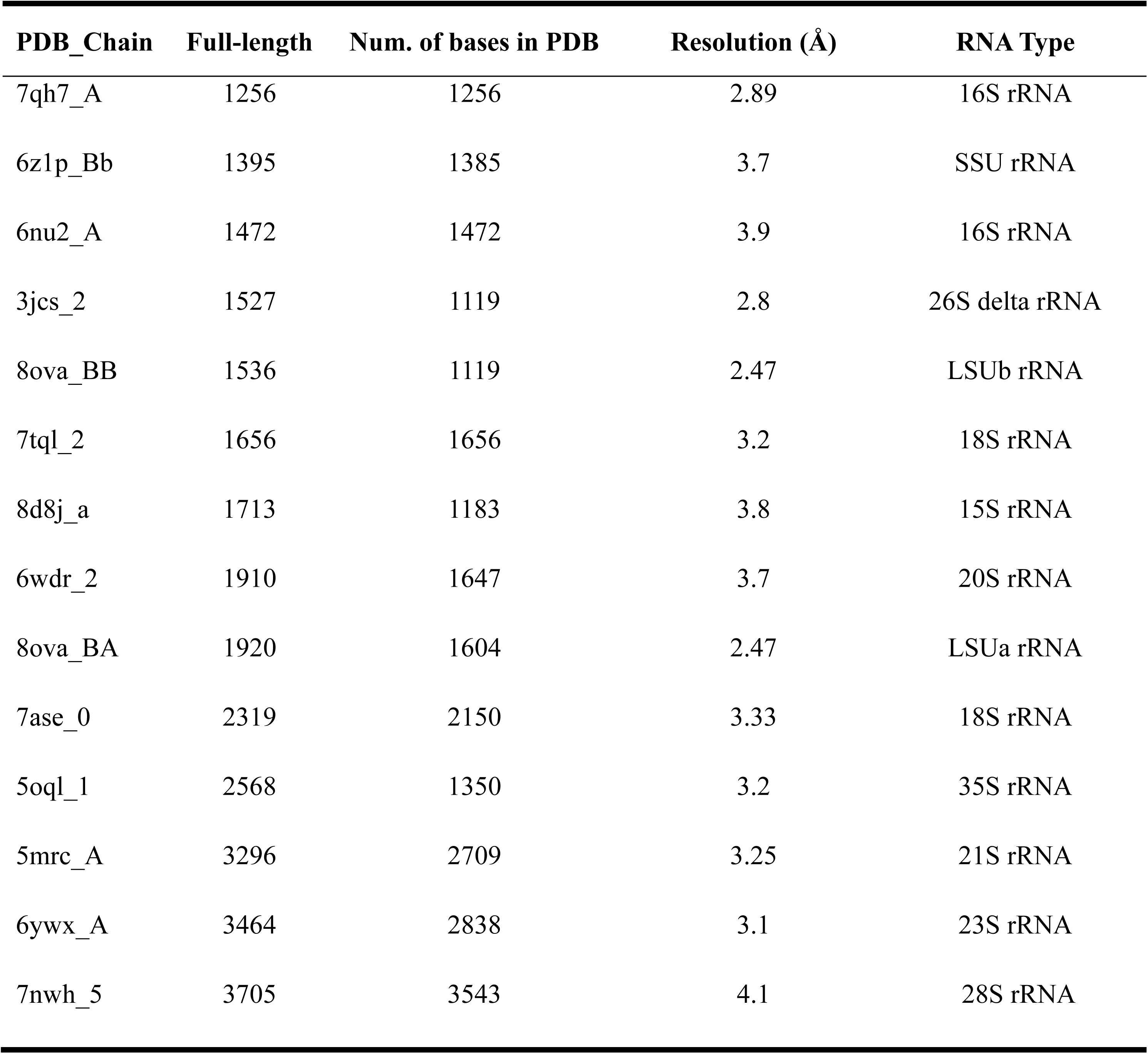
The long RNA dataset. Detailed information of 14 non-redundant long RNA structures (from 1,256 to 3,705 nucleotides) used for evaluating RESM’s and RESM-basepair’s generalization capability beyond their training length limit of 1,024. In detail, RNA chains with a sequence length greater than 1,024 and a base count greater than 500 (bases with a determined structure) in the PDB structure were first collected from the RNA3DB structure database. Subsequently, based on sequence homology (identity < 80%) and PDB structural similarity (TMscore < 0.45). When predicting secondary structures, the full-length RNA sequences were input into each method.

**Supplementary Table 7.**
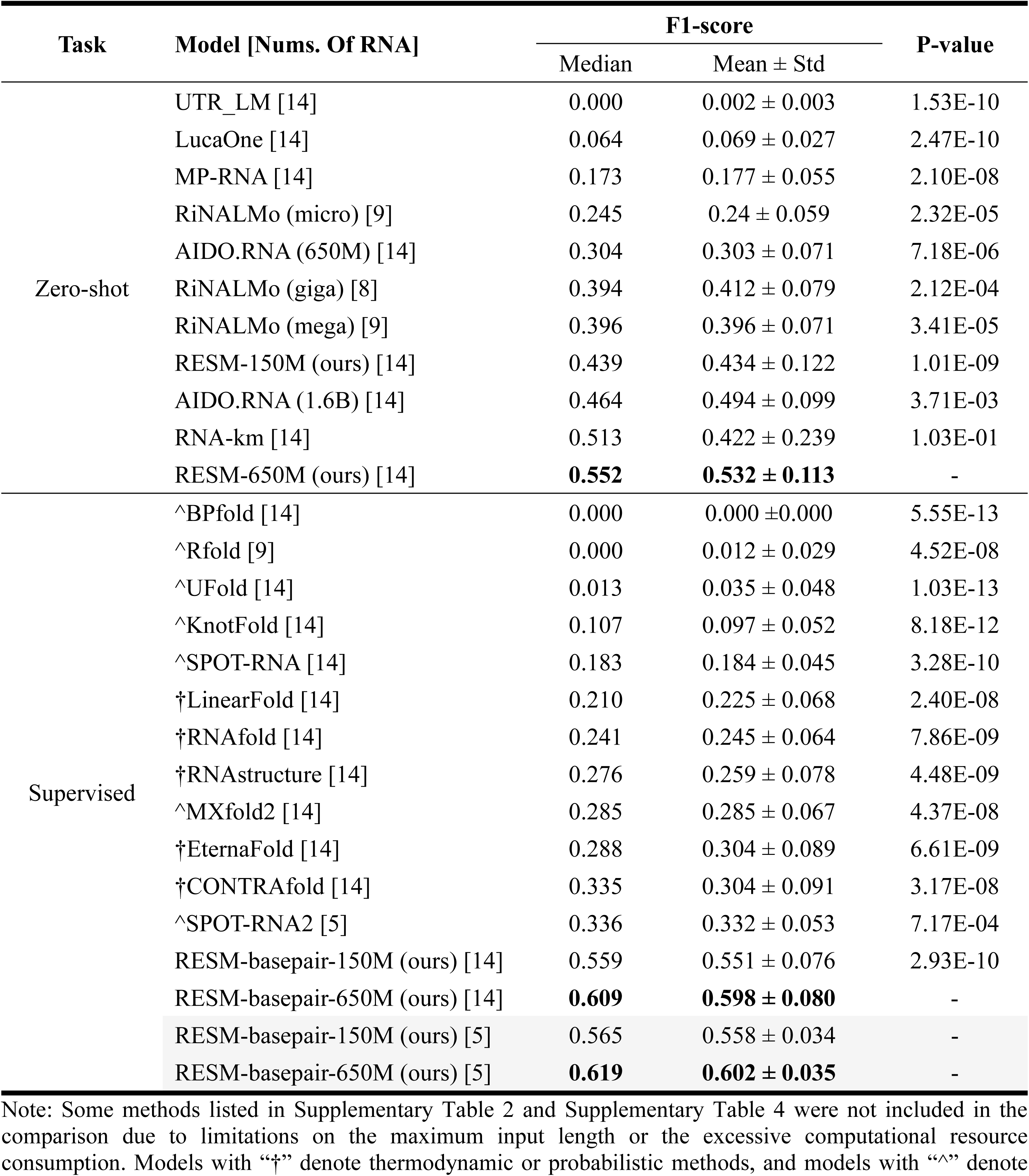
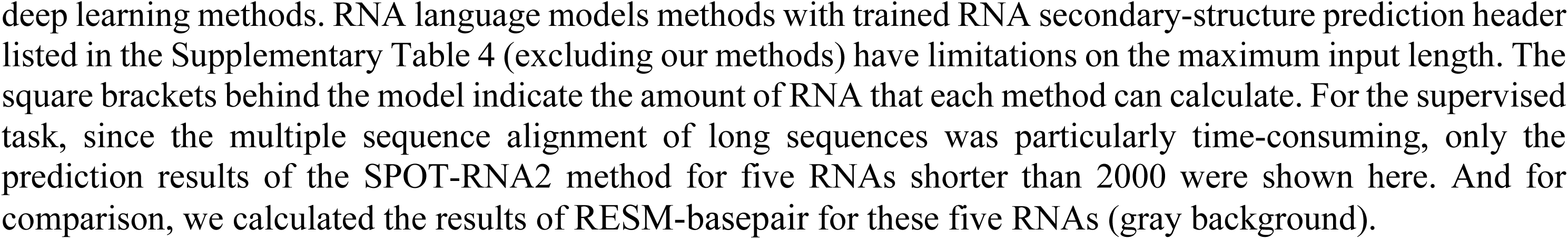
Performance of RNA secondary-structure prediction on long RNA dataset, according to the F1 score. The evaluation contained zero-shot prediction and supervised prediction, whose processes were consistent with Supplementary Table 2 and Supplementary Table 4, respectively. And in the zero-shot prediction process, the threshold was first evaluated on the VL46 dataset and then tested on the long RNA dataset. P-values were calculated based on the results of RESM-650M (zero-shot prediction) or RESM-basepair-650M (supervised prediction).

| Task | Model [Nums. Of RNA] | F1-score |  | P-value |
| --- | --- | --- | --- | --- |
| | | Median | Mean ± Std | |
| Zero-shot | UTR_LM [14] | 0.000 | 0.002 ± 0.003 | 1.53E-10 |
| | LucaOne [14] | 0.064 | 0.069 ± 0.027 | 2.47E-10 |
| | MP-RNA [14] | 0.173 | 0.177 ± 0.055 | 2.10E-08 |
| | RiNALMo (micro) [9] | 0.245 | 0.24 ± 0.059 | 2.32E-05 |
| | AIDO.RNA (650M) [14] | 0.304 | 0.303 ± 0.071 | 7.18E-06 |
| | RiNALMo (giga) [8] | 0.394 | 0.412 ± 0.079 | 2.12E-04 |
| | RiNALMo (mega) [9] | 0.396 | 0.396 ± 0.071 | 3.41E-05 |
| | RESM-150M (ours) [14] | 0.439 | 0.434 ± 0.122 | 1.01E-09 |
| | AIDO.RNA (1.6B) [14] | 0.464 | 0.494 ± 0.099 | 3.71E-03 |
| | RNA-km [14] | 0.513 | 0.422 ± 0.239 | 1.03E-01 |
|  | RESM-650M (ours) [14] | **0.552** | **0.532 ± 0.113** | - |
| Supervised | ^BPfold [14] | 0.000 | 0.000 ± 0.000 | 5.55E-13 |
| | ^Rfold [9] | 0.000 | 0.012 ± 0.029 | 4.52E-08 |
| | ^UFold [14] | 0.013 | 0.035 ± 0.048 | 1.03E-13 |
| | ^KnotFold [14] | 0.107 | 0.097 ± 0.052 | 8.18E-12 |
| | ^SPOT-RNA [14] | 0.183 | 0.184 ± 0.045 | 3.28E-10 |
| | †LinearFold [14] | 0.210 | 0.225 ± 0.068 | 2.40E-08 |
| | †RNAfold [14] | 0.241 | 0.245 ± 0.064 | 7.86E-09 |
| | †RNAstructure [14] | 0.276 | 0.259 ± 0.078 | 4.48E-09 |
| | ^MXfold2 [14] | 0.285 | 0.285 ± 0.067 | 4.37E-08 |
| | †EternaFold [14] | 0.288 | 0.304 ± 0.089 | 6.61E-09 |
| | †CONTRAFold [14] | 0.335 | 0.304 ± 0.091 | 3.17E-08 |
| | ^SPOT-RNA2 [5] | 0.336 | 0.332 ± 0.053 | 7.17E-04 |
| | RESM-basepair-150M (ours) [14] | 0.559 | 0.551 ± 0.076 | 2.93E-10 |
|  | RESM-basepair-650M (ours) [14] | **0.609** | **0.598 ± 0.080** | - |
| | RESM-basepair-150M (ours) [5] | 0.565 | 0.558 ± 0.034 | - |
|  | RESM-basepair-650M (ours) [5] | **0.619** | **0.602 ± 0.035** | - |

Note: Some methods listed in Supplementary Table 2 and Supplementary Table 4 were not included in the comparison due to limitations on the maximum input length or the excessive computational resource consumption. Models with “†” denote thermodynamic or probabilistic methods, and models with “^” denote

**Supplementary Table 8.**
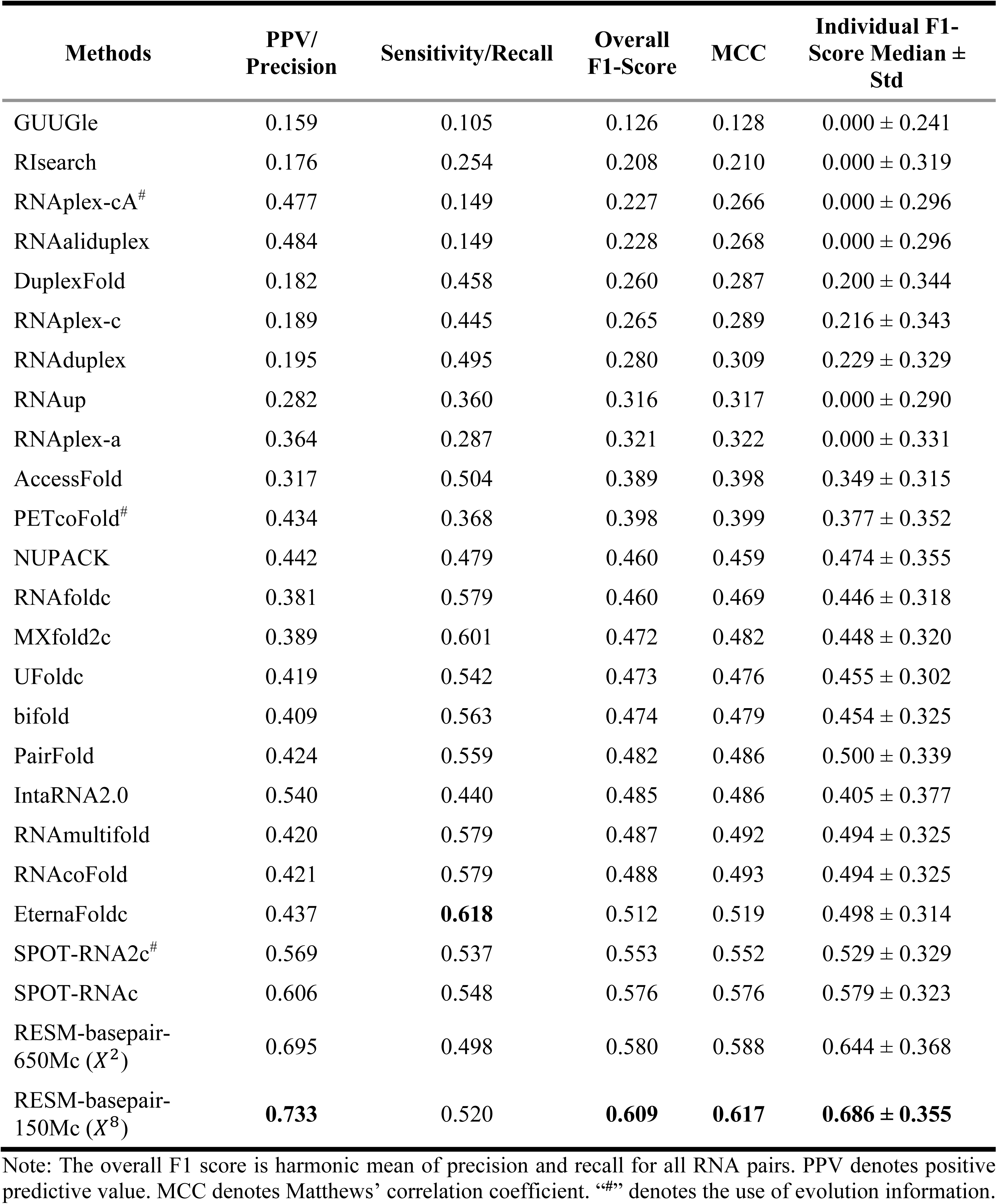
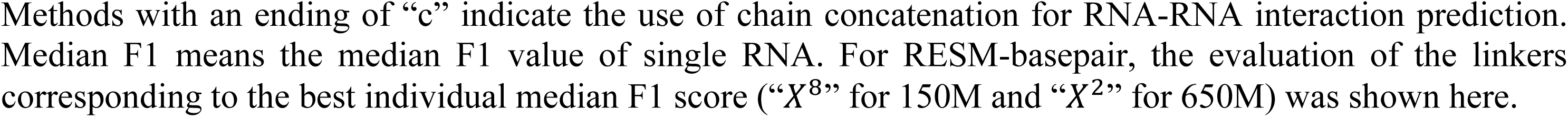
Performance on zero-shot RNA–RNA interaction prediction. Comparison of 23 predictors of inter-RNA base pairs on 64 complexes of RNA structures unseen by SPOT-RNA and SPOT-RNA2. All single-chain structures in the test set have the structural similarity score TM-Score < 0.3 compared to the monomeric structures used in training and validating SPOT-RNA and SPOT-RNA2 methods.

| Methods | PPV/<br>Precision | Sensitivity/Recall | Overall<br>F1-Score | MCC | Individual F1-Score Median ±<br>Std |
| --- | --- | --- | --- | --- | --- |
| GUUGle | 0.159 | 0.105 | 0.126 | 0.128 | 0.000 ± 0.241 |
| RIsearch | 0.176 | 0.254 | 0.208 | 0.210 | 0.000 ± 0.319 |
| RNAplex-cA<sup>#</sup> | 0.477 | 0.149 | 0.227 | 0.266 | 0.000 ± 0.296 |
| RNAaliduplex | 0.484 | 0.149 | 0.228 | 0.268 | 0.000 ± 0.296 |
| DuplexFold | 0.182 | 0.458 | 0.260 | 0.287 | 0.200 ± 0.344 |
| RNAplex-c | 0.189 | 0.445 | 0.265 | 0.289 | 0.216 ± 0.343 |
| RNAduplex | 0.195 | 0.495 | 0.280 | 0.309 | 0.229 ± 0.329 |
| RNAup | 0.282 | 0.360 | 0.316 | 0.317 | 0.000 ± 0.290 |
| RNAplex-a | 0.364 | 0.287 | 0.321 | 0.322 | 0.000 ± 0.331 |
| AccessFold | 0.317 | 0.504 | 0.389 | 0.398 | 0.349 ± 0.315 |
| PETcoFold<sup>#</sup> | 0.434 | 0.368 | 0.398 | 0.399 | 0.377 ± 0.352 |
| NUPACK | 0.442 | 0.479 | 0.460 | 0.459 | 0.474 ± 0.355 |
| RNAfoldc | 0.381 | 0.579 | 0.460 | 0.469 | 0.446 ± 0.318 |
| MXfold2c | 0.389 | 0.601 | 0.472 | 0.482 | 0.448 ± 0.320 |
| UFoldc | 0.419 | 0.542 | 0.473 | 0.476 | 0.455 ± 0.302 |
| bifold | 0.409 | 0.563 | 0.474 | 0.479 | 0.454 ± 0.325 |
| PairFold | 0.424 | 0.559 | 0.482 | 0.486 | 0.500 ± 0.339 |
| IntaRNA2.0 | 0.540 | 0.440 | 0.485 | 0.486 | 0.405 ± 0.377 |
| RNAmultifold | 0.420 | 0.579 | 0.487 | 0.492 | 0.494 ± 0.325 |
| RNAcoFold | 0.421 | 0.579 | 0.488 | 0.493 | 0.494 ± 0.325 |
| EternaFoldc | 0.437 | **0.618** | 0.512 | 0.519 | 0.498 ± 0.314 |
| SPOT-RNA2c<sup>#</sup> | 0.569 | 0.537 | 0.553 | 0.552 | 0.529 ± 0.329 |
| SPOT-RNAc | 0.606 | 0.548 | 0.576 | 0.576 | 0.579 ± 0.323 |
| RESM-basepair-650Mc ($X^2$) | 0.695 | 0.498 | 0.580 | 0.588 | 0.644 ± 0.368 |
| RESM-basepair-150Mc ($X^8$) | **0.733** | 0.520 | **0.609** | **0.617** | **0.686 ± 0.355** |

Note: The overall F1 score is harmonic mean of precision and recall for all RNA pairs. PPV denotes positive predictive value. MCC denotes Matthews' correlation coefficient. “<sup>#</sup>” denotes the use of evolution information.
Methods with an ending of “c” indicate the use of chain concatenation for RNA-RNA interaction prediction. Median F1 means the median F1 value of single RNA. For RESM-basepair, the evaluation of the linkers corresponding to the best individual median F1 score (“X<sup>8</sup>” for 150M and “X<sup>2</sup>” for 650M) was shown here.

**Supplementary Table 9.** Performance on zero-shot RNA–RNA interaction prediction by RESM-basepair-150M with and without linker sequences. Performance is reported using precision (PPV), recall (sensitivity), overall F1 score, Matthews correlation coefficient (MCC), and the median ± standard deviation of individual F1 scores across samples. “None” indicates no linker was used between input RNA pairs. Each 𝑋^𝑛^ denotes a linker sequence composed by concatenating *n* repetitions of “X”.

| Linker | PPV/Precision | Sensitivity/Recall | Overall F1-Score | MCC | Individual F1-Score Median ± Std |
| --- | --- | --- | --- | --- | --- |
| *None* | 0.687 | 0.523 | 0.594 | 0.599 | 0.623 ± 0.345 |
| $X^1$ | 0.698 | 0.520 | 0.596 | 0.601 | 0.628 ± 0.356 |
| $X^2$ | 0.702 | 0.518 | 0.596 | 0.602 | 0.653 ± 0.358 |
| $X^3$ | 0.707 | 0.523 | 0.601 | 0.608 | 0.641 ± 0.360 |
| $X^4$ | 0.728 | **0.534** | **0.616** | 0.623 | 0.653 ± 0.344 |
| $X^5$ | 0.730 | 0.517 | 0.605 | 0.614 | 0.653 ± 0.356 |
| $X^6$ | 0.731 | 0.533 | **0.616** | 0.624 | 0.651 ± 0.344 |
| $X^7$ | **0.743** | 0.527 | **0.616** | **0.625** | 0.664 ± 0.351 |
| $X^8$ | 0.733 | 0.520 | 0.609 | 0.617 | **0.686 ± 0.355** |
| $X^9$ | 0.738 | 0.513 | 0.605 | 0.615 | 0.679 ± 0.364 |
| $X^{10}$ | 0.735 | 0.516 | 0.606 | 0.615 | 0.679 ± 0.359 |
| $X^{11}$ | 0.736 | 0.518 | 0.608 | 0.617 | 0.653 ± 0.358 |
| $X^{12}$ | 0.735 | 0.521 | 0.610 | 0.619 | 0.637 ± 0.353 |

**Supplementary Table 10.** Performance of zero-shot RNA–RNA interaction prediction by RESM-basepair-650M with and without linker sequences. Performance metrics include precision (PPV), recall (sensitivity), overall F1 score, Matthews correlation coefficient (MCC), and the median ± standard deviation of individual F1 scores across test samples. “None” indicates that no linker was used between paired input RNAs. Each linker type “𝑋^𝑛^” represents a sequence of *n* repetitions of the token “X”. The results demonstrate that moderate-length linkers (e.g., 𝑋^4^) can improve individual F1 scores, while longer linkers (e.g., 𝑋^12^) yield the best overall F1 score and MCC.

| Linker | PPV/Precision | Sensitivity/Recall | Overall F1-Score | MCC | Individual F1-Score Median ± Std |
| --- | --- | --- | --- | --- | --- |
| *None* | 0.668 | 0.499 | 0.571 | 0.577 | 0.636 ± 0.364 |
| $X^1$ | 0.689 | 0.498 | 0.578 | 0.585 | 0.636 ± 0.369 |
| $X^2$ | 0.695 | 0.498 | 0.580 | 0.588 | **0.644 ± 0.368** |
| $X^3$ | 0.684 | 0.500 | 0.578 | 0.584 | 0.639 ± 0.371 |
| $X^4$ | 0.680 | 0.501 | 0.577 | 0.583 | **0.644 ± 0.373** |
| $X^5$ | 0.672 | 0.501 | 0.574 | 0.580 | 0.637 ± 0.367 |
| $X^6$ | 0.685 | 0.498 | 0.577 | 0.583 | 0.639 ± 0.370 |
| $X^7$ | 0.692 | 0.499 | 0.580 | 0.587 | 0.636 ± 0.374 |
| $X^8$ | 0.694 | 0.499 | 0.581 | 0.588 | 0.628 ± 0.365 |
| $X^9$ | 0.701 | **0.504** | 0.586 | 0.594 | 0.630 ± 0.359 |
| $X^{10}$ | 0.711 | 0.498 | 0.586 | 0.595 | 0.620 ± 0.358 |
| $X^{11}$ | **0.725** | 0.497 | 0.590 | 0.600 | 0.602 ± 0.357 |
| $X^{12}$ | **0.725** | 0.502 | **0.593** | **0.603** | 0.603 ± 0.352 |

**Supplementary Table 11.**
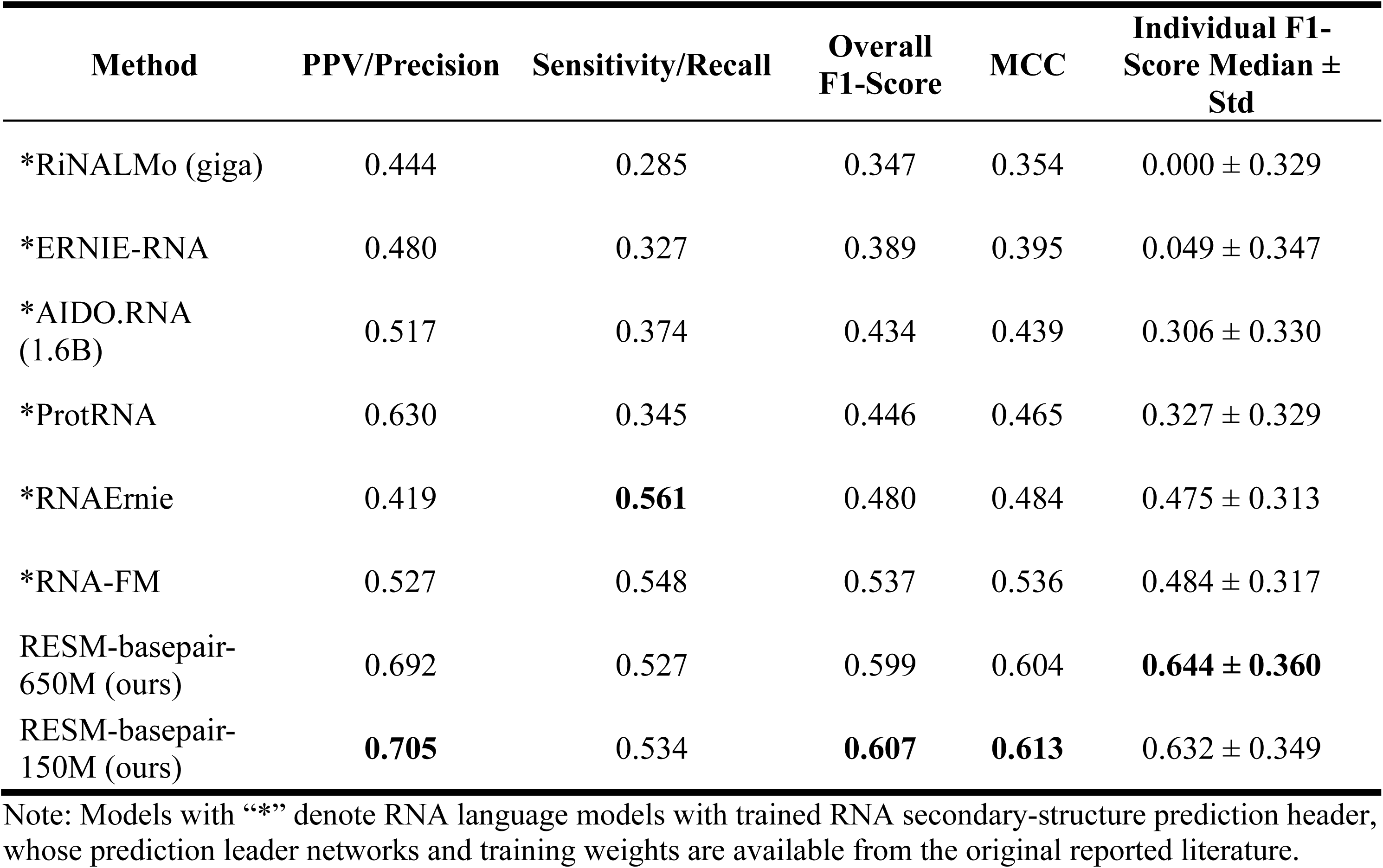
Performance of zero-shot RNA–RNA interaction prediction of RNA language models with secondary-structure prediction heads. Models were evaluated on 62 RNA complexes, excluding 2 complexes with sequences exceeding 510 nucleotides. All linker sequences were set to *None* in this evaluation.

| Method | PPV/Precision | Sensitivity/Recall | Overall F1-Score | MCC | Individual F1-Score Median ± Std |
| --- | --- | --- | --- | --- | --- |
| *RiNALMo (giga) | 0.444 | 0.285 | 0.347 | 0.354 | 0.000 ± 0.329 |
| *ERNIE-RNA | 0.480 | 0.327 | 0.389 | 0.395 | 0.049 ± 0.347 |
| *AIDO.RNA (1.6B) | 0.517 | 0.374 | 0.434 | 0.439 | 0.306 ± 0.330 |
| *ProtRNA | 0.630 | 0.345 | 0.446 | 0.465 | 0.327 ± 0.329 |
| *RNAErnie | 0.419 | **0.561** | 0.480 | 0.484 | 0.475 ± 0.313 |
| *RNA-FM | 0.527 | 0.548 | 0.537 | 0.536 | 0.484 ± 0.317 |
| RESM-basepair-650M (ours) | 0.692 | 0.527 | 0.599 | 0.604 | **0.644 ± 0.360** |
| RESM-basepair-150M (ours) | **0.705** | 0.534 | **0.607** | **0.613** | 0.632 ± 0.349 |

Note: Models with “\*” denote RNA language models with trained RNA secondary-structure prediction header, whose prediction leader networks and training weights are available from the original reported literature.

**Supplementary Table 12.**
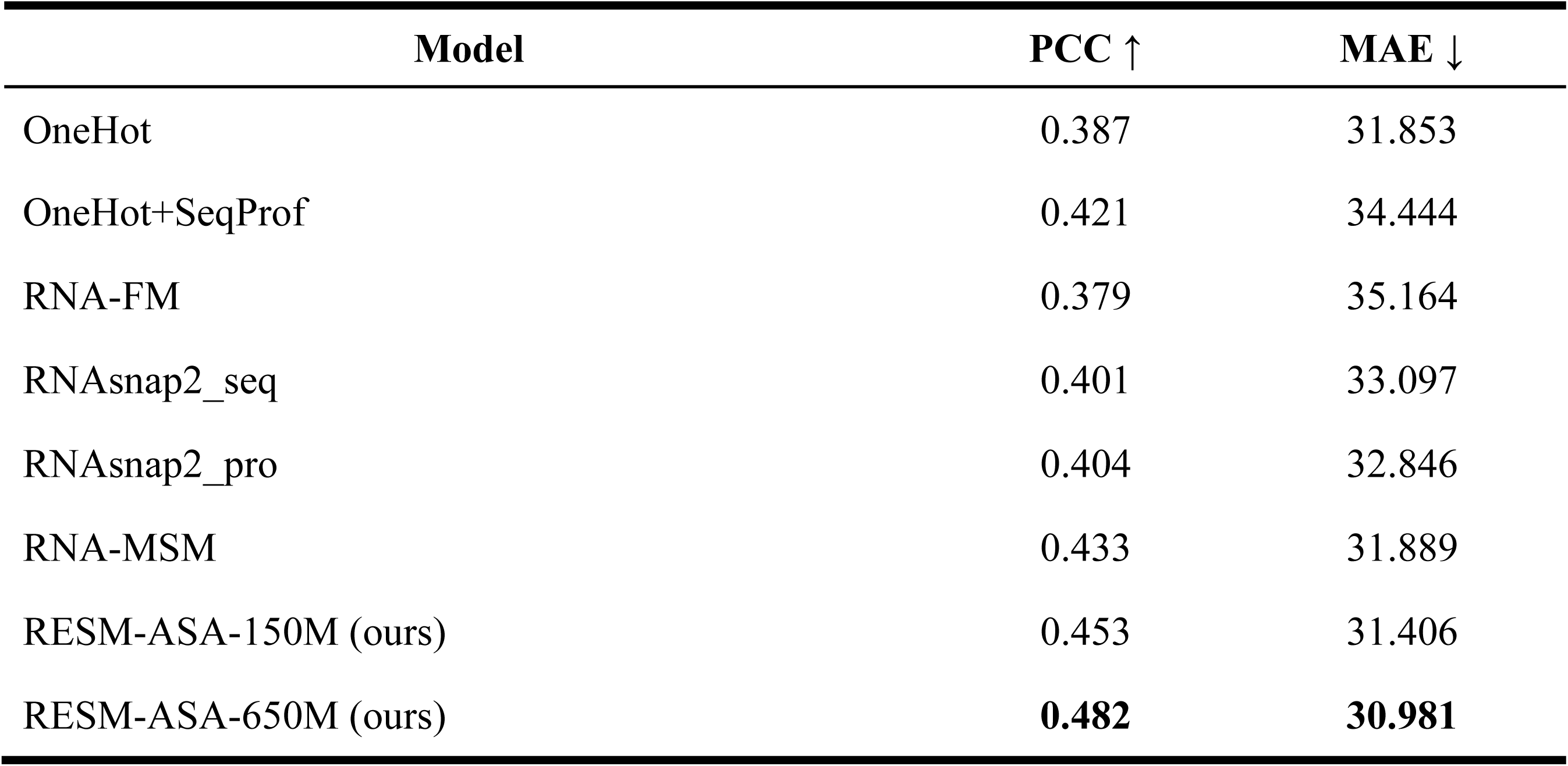
Supervised RNA solvent accessibility prediction on the RNA-MSM benchmark by RESM-ASA as compared to other methods. Pearson correlation coefficient (PCC, higher is better) and mean absolute error (MAE in Å^2^, lower is better) are reported for each model. RESM-650M achieves the best performance, outperforming RNA-FM, RNA-MSM, RNAsnap2, SeqProf and one-hot encoding approaches

**Supplementary Table 13.**
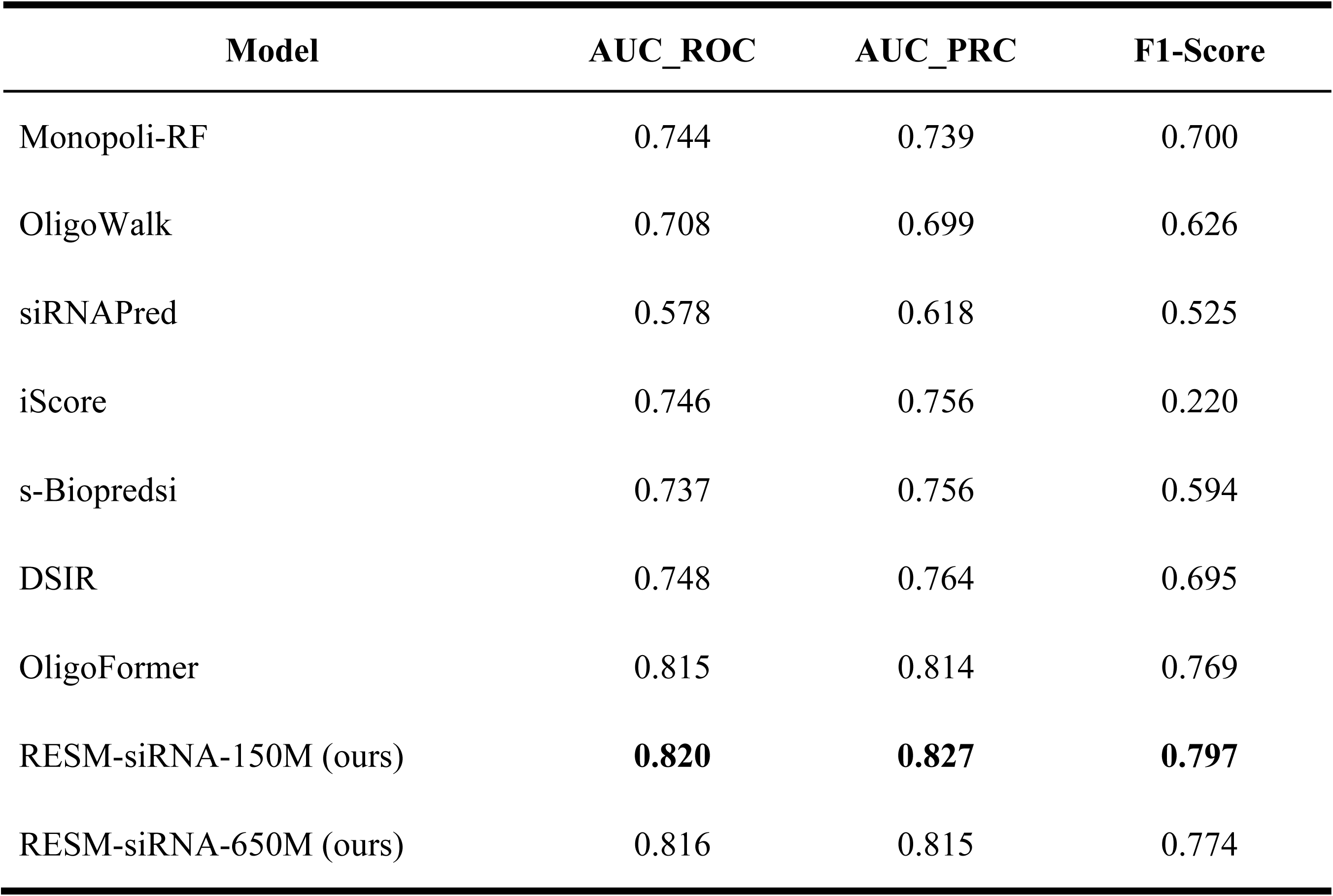
Performance on supervised siRNA activity prediction across benchmark models. Performance was evaluated using three metrics: area under the ROC curve (AUC_ROC), area under the precision-recall curve (AUC_PRC), and F1 score. RESM-siRNA-150M achieves the highest performance across all metrics, surpassing previous methods as shown.

**Supplementary Table 14.**
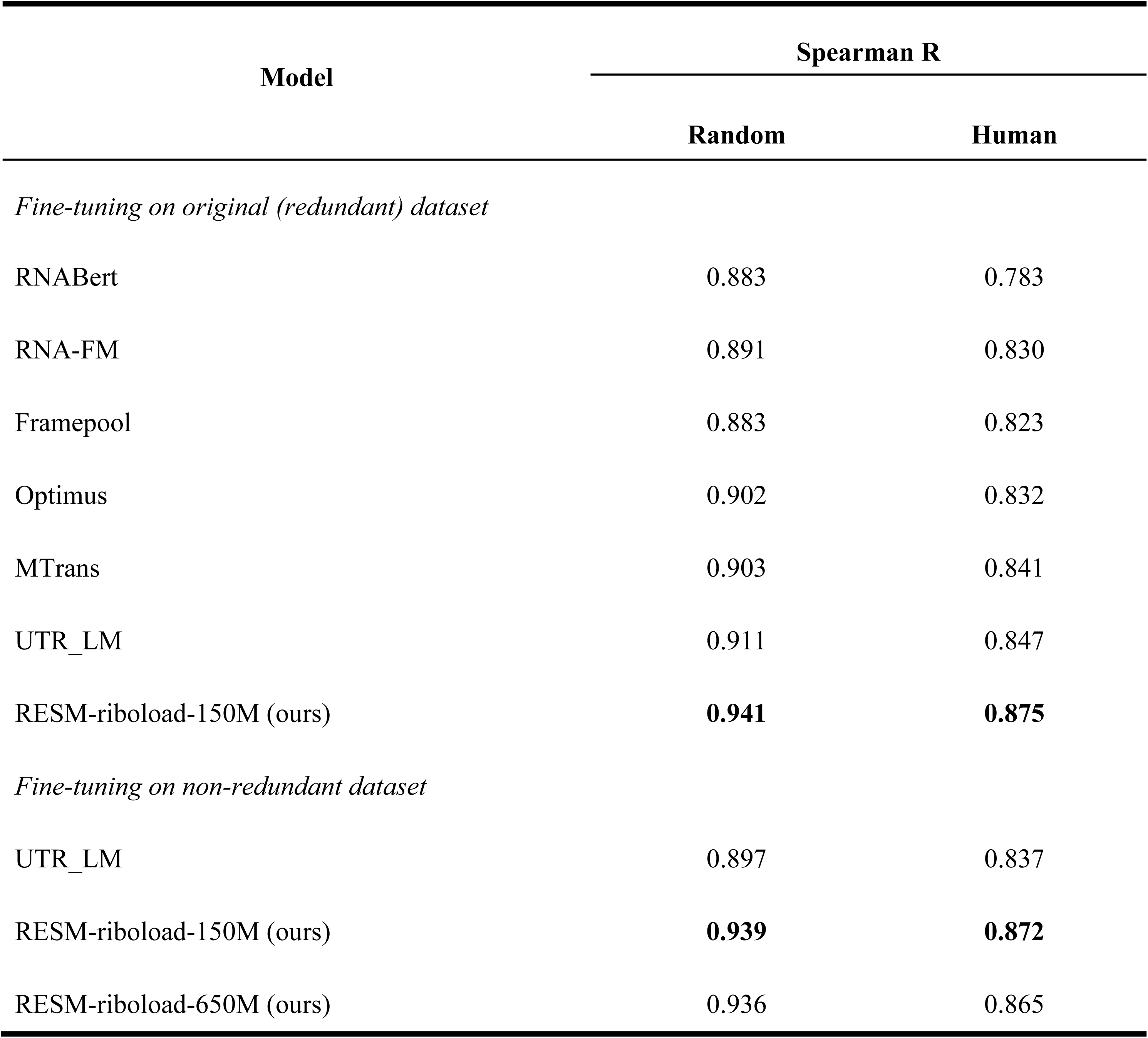
Performance on supervised ribosome loading prediction across random and human-derived 5’UTRs. Spearman’s rank correlation coefficients were reported for each model on two test sets: synthetic random 5’UTR sequences (“Random”) and human-derived 5’UTRs (“Human”). Upper section shows results from models fine-tuned on the non-redundant dataset, while the lower section compares performance when fine-tuned on the original (redundant) dataset. RESM-riboload-150M achieves the highest performance across both settings, with further gains observed when training on the non-redundant dataset.

**Supplementary Table 15.**
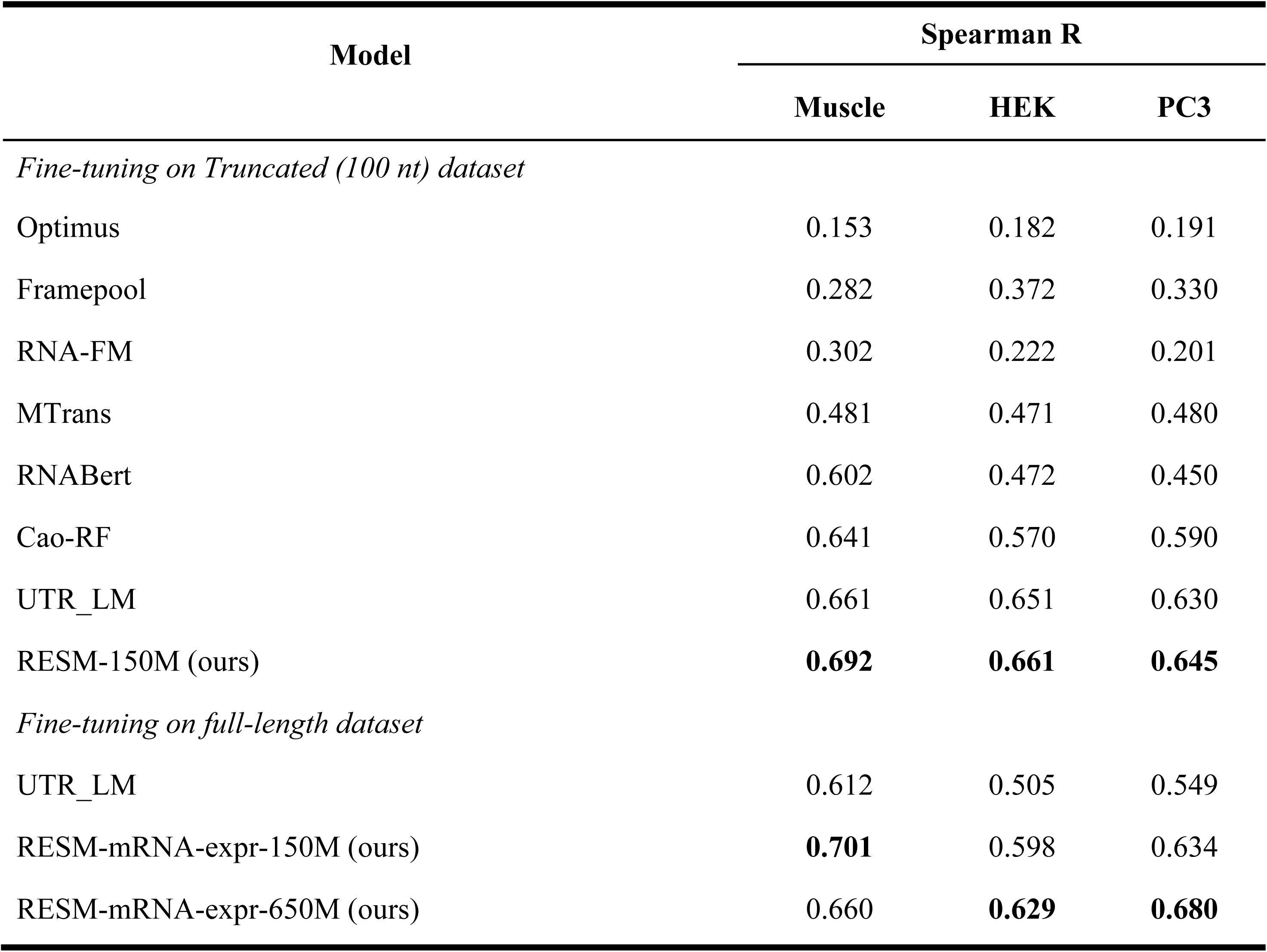
Performance on supervised mRNA expression-level prediction across human tissues and cell lines by RESM-mRNAexpr. Spearman’s rank correlation coefficients were reported for each model on three datasets: human skeletal muscle, prostate cancer (PC3) cells, and human embryonic kidney (HEK) 293T cells. The upper section presents results from models fine-tuned on truncated 5’UTR sequences (first 100 nucleotides), while the lower section reports performance when full-length RNA sequences were used as input.

### Pretraining details

For model pretraining, we adopted the AdamW optimizer with a learning rate of 1e-3, weight decay of 1e-4, and β₁ = 0.9, β₂ = 0.98. To stabilize early-stage training, we employed a warmup linear learning rate scheduler with 1,600 warmup steps, followed by a linear decay schedule. All models were trained using masked language modeling (MLM) as the training objective.

We followed the ESM-2 architecture exactly, adapting its design to the RNA domain without altering core architectural components. The masking strategy was consistent with standard MLM settings: 15% of tokens were randomly masked during training. Importantly, instead of using an RNA-specific vocabulary for prediction, we adopted the protein vocabulary from ESM-2 as the decoding target.

The RESM-150M model was trained for approximately 460k steps on 8 NVIDIA Tesla A100 80GB GPUs, while the larger RESM-650M model was trained for approximately 100k steps on 256 Huawei Ascend D910B GPUs to accommodate its greater parameter count and computational demand.

### Downstream task details

#### Supervised secondary structure prediction and Zero-shot RNA-RNA interaction prediction

**Supplementary Table 16.**
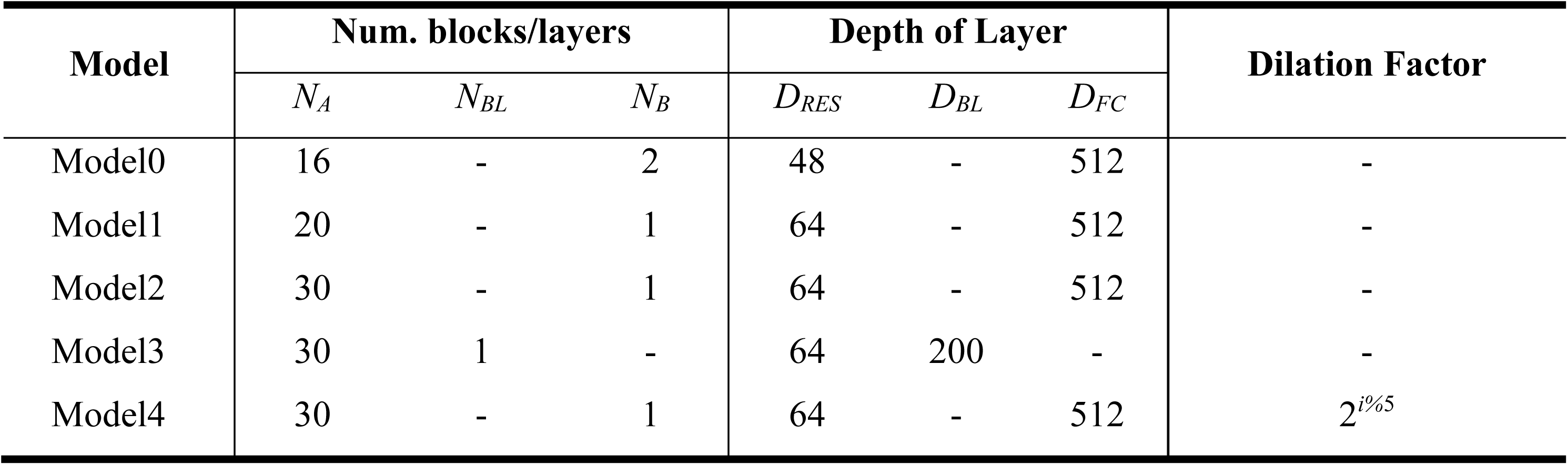
The model hyperparameters for supervised secondary-structure prediction used in RESM-150M-basepair or RESM-650M-basepair ensembles. Each model variant shares the same architectural design as SPOT-RNA. *N_A_* denotes the number of the ResNet blocks; *N_BL_* is the number of 2D-LSTM layers; *N_B_*, represents the number of the fully connected blocks; *D_RES_*, *D_BL_* and *D_FC_* refer to the layer depths in ResNet blocks, 2D-LSTM cells (number of nodes per direction), and fully connected layers, respectively. The dilation factor is specified where applicable. Model4 incorporates a dilation factor of 2*^i%^*^5^, following the dilated convolution strategy described in the original SPOT-RNA framework.

**Supplementary Table 16.**
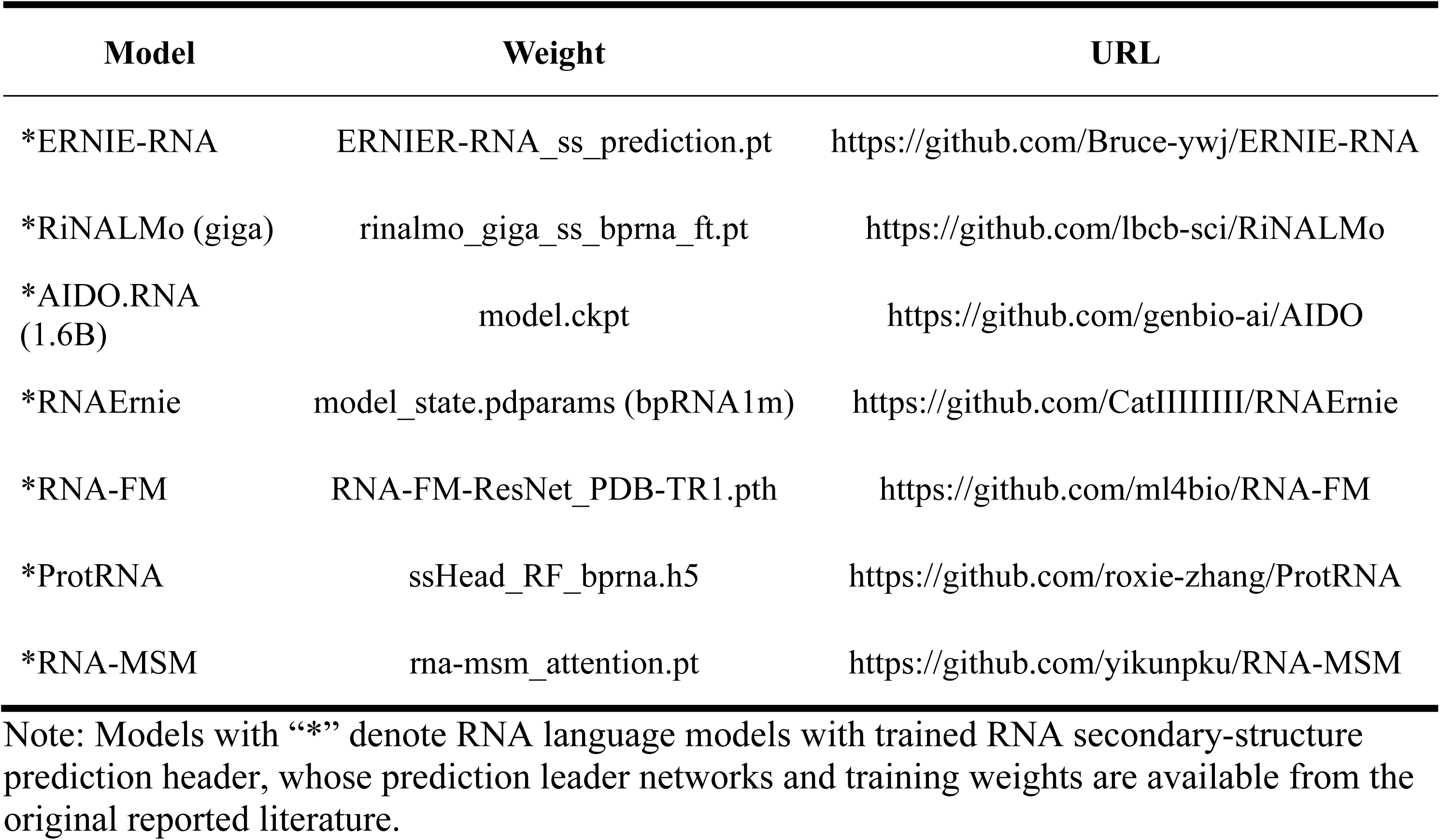
The locations of model weights for supervised secondary-structure prediction heads of compared RNA language models.

**Supplementary Fig. 1.**
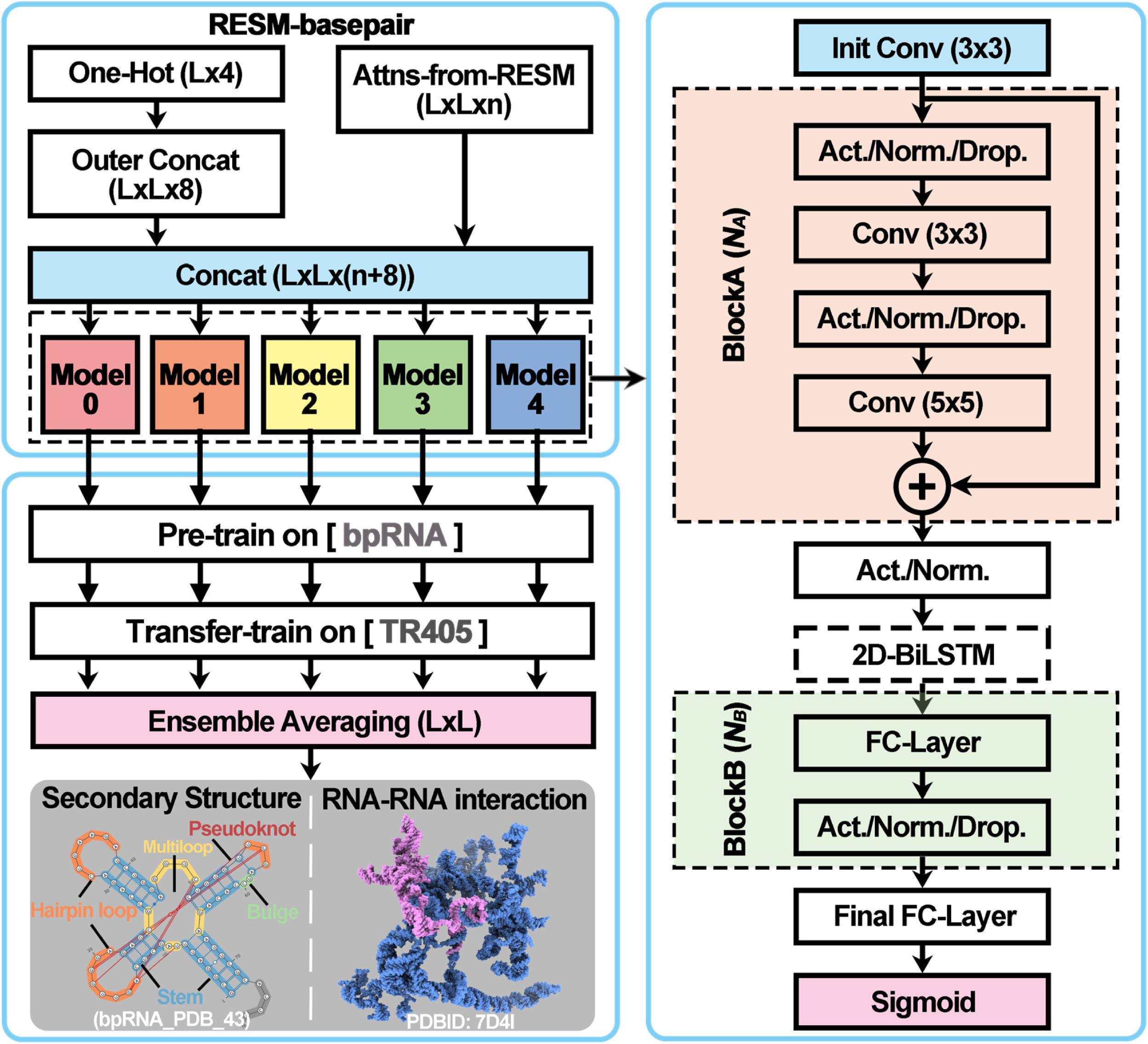
Generalized model architecture and training process of RESM-basepair. The full RESM-basepair model consists of five independently trained (including pre-training on the bpRNA dataset and transfer training on the TR405 dataset) models with similar network structure (shown on the right), whose outputs are ensemble averaged to obtain the final prediction. The input is made of two parts, one is the one-hot encoding feature with dimension 𝐿 × 4 and the other is RESM attention maps with dimension 𝐿 × 𝐿 × 𝑟, where 𝐿 is the sequence length of the target RNA and 𝑟 is the number of attention maps, which is 600 for the RESM-150M model and 660 for the RESM-650M model. “Outer Concat” indicates the outer concatenation operation; “Conv” indicates the convolutional layer; “Act.” indicates the activation function; “Norm.” indicates the normalization function; “Drop.” indicates the dropout layer; “FC” indicates the fully-connected layer and “Sigmoid” indicates the sigmoid function. The module “BlockA” consists of 𝑁_𝐴_ blocks and the module “BlockB” consists of 𝑁_𝐵_ blocks. The “2D-BiLSTM” layer is only used in “Model 3”.

#### Supervised RNA solvent accessibility prediction

**Supplementary Fig. 2.**
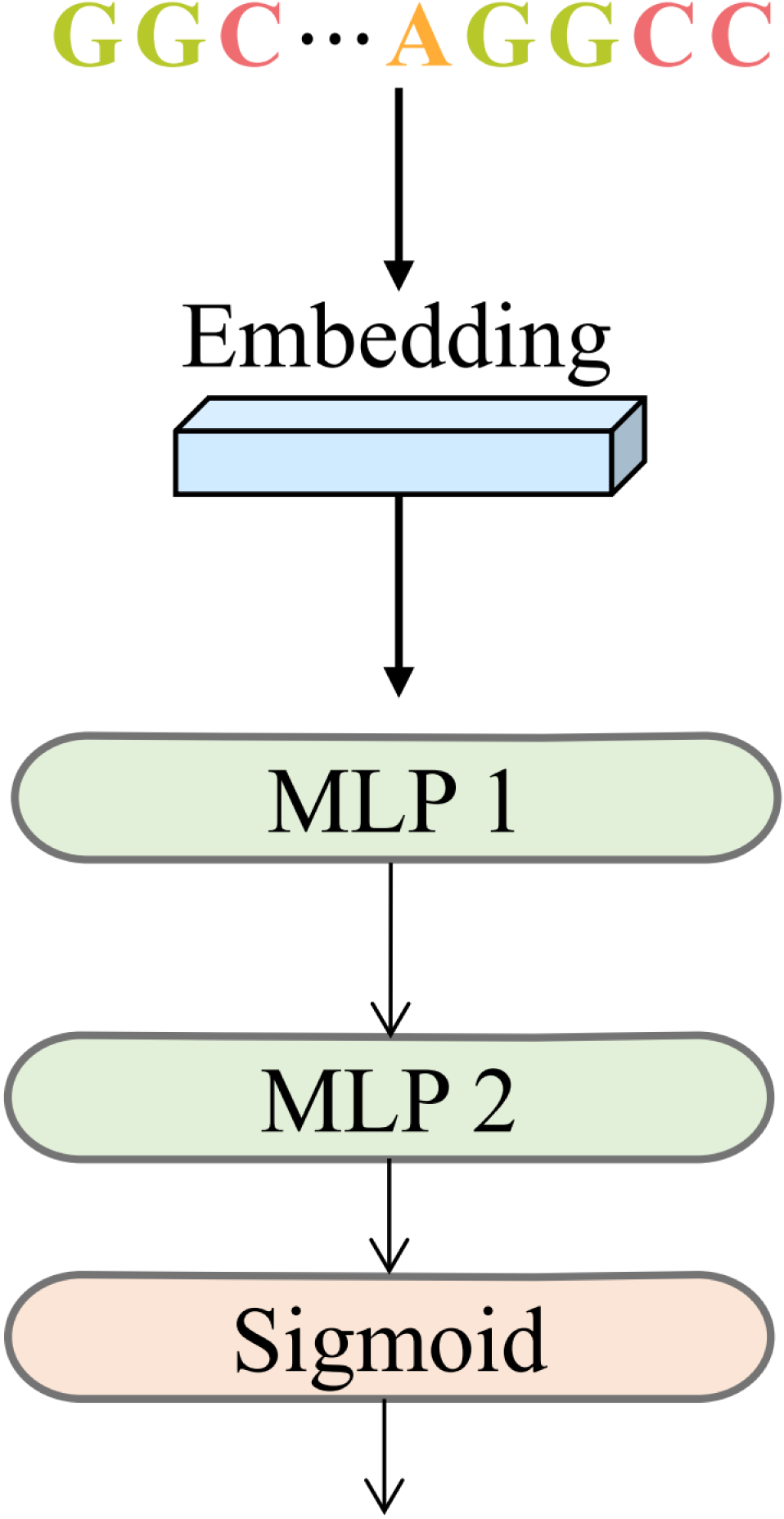
Model architecture of the supervised RNA solvent-accessibility prediction (RESM-ASA). Final-layer embeddings from the pre-trained RESM model are passed through a two-layer multilayer perceptron (MLP), followed by a sigmoid activation to predict nucleotide-wise relative solvent accessibility (RSA) values ranging from 0 to 1. Compared to the more complex architecture used in RNA-MSM, this model adopts a simpler and more efficient design while maintaining high predictive performance. The predicted relative solvent-accessibility values are subsequently converted to absolute solvent-accessible surface area (ASA) using predefined normalization factors.

#### Supervised siRNA activity prediction

**Supplementary Fig. 3.**
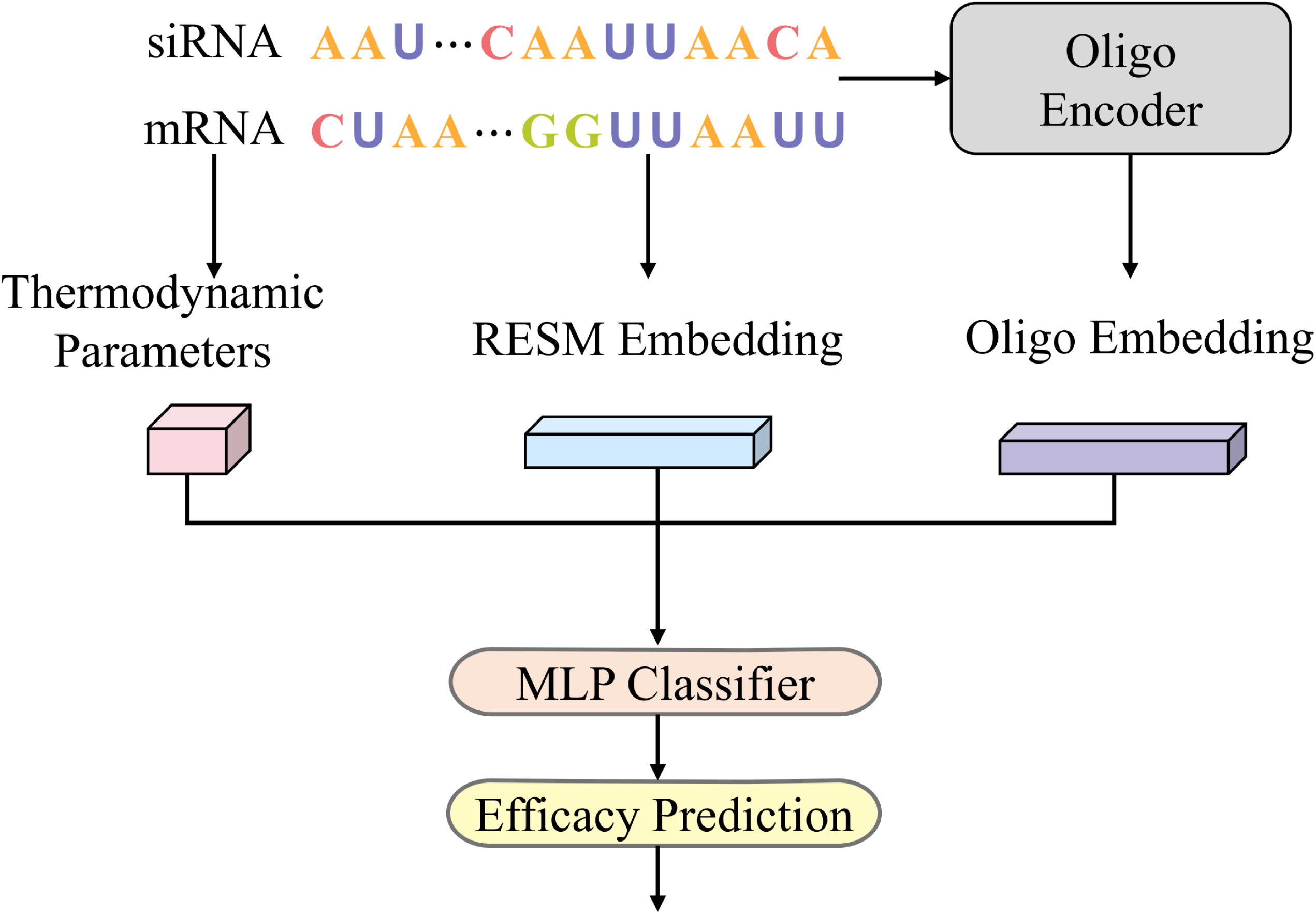
Architecture of the supervised siRNA activity prediction model (RESM-siRNA). The framework integrates three parallel feature extraction modules: (1) the RESM Embedding Module encodes both siRNA sequences and flanking mRNA regions using final-layer embeddings from the pre-trained RESM model; (2) the Oligo Encoder processes input sequences using a hybrid architecture consisting of 2D convolutions, pooling layers, a BiLSTM, and a transformer module to capture spatial and sequential interaction patterns; and (3) a thermodynamic feature module extracts siRNA–mRNA pairing properties. The outputs of all modules are concatenated and fed into a multilayer perceptron (MLP) classifier to predict siRNA efficacy scores ranging from 0 to 1. This model adopts the overall design of OligoFormer but replaces RNA-FM with RESM for embedding generation, enabling improved representation learning for RNA-guided targeting tasks.

#### Supervised ribosome loading and expression levels prediction in mRNA

**Supplementary Fig. 4.**
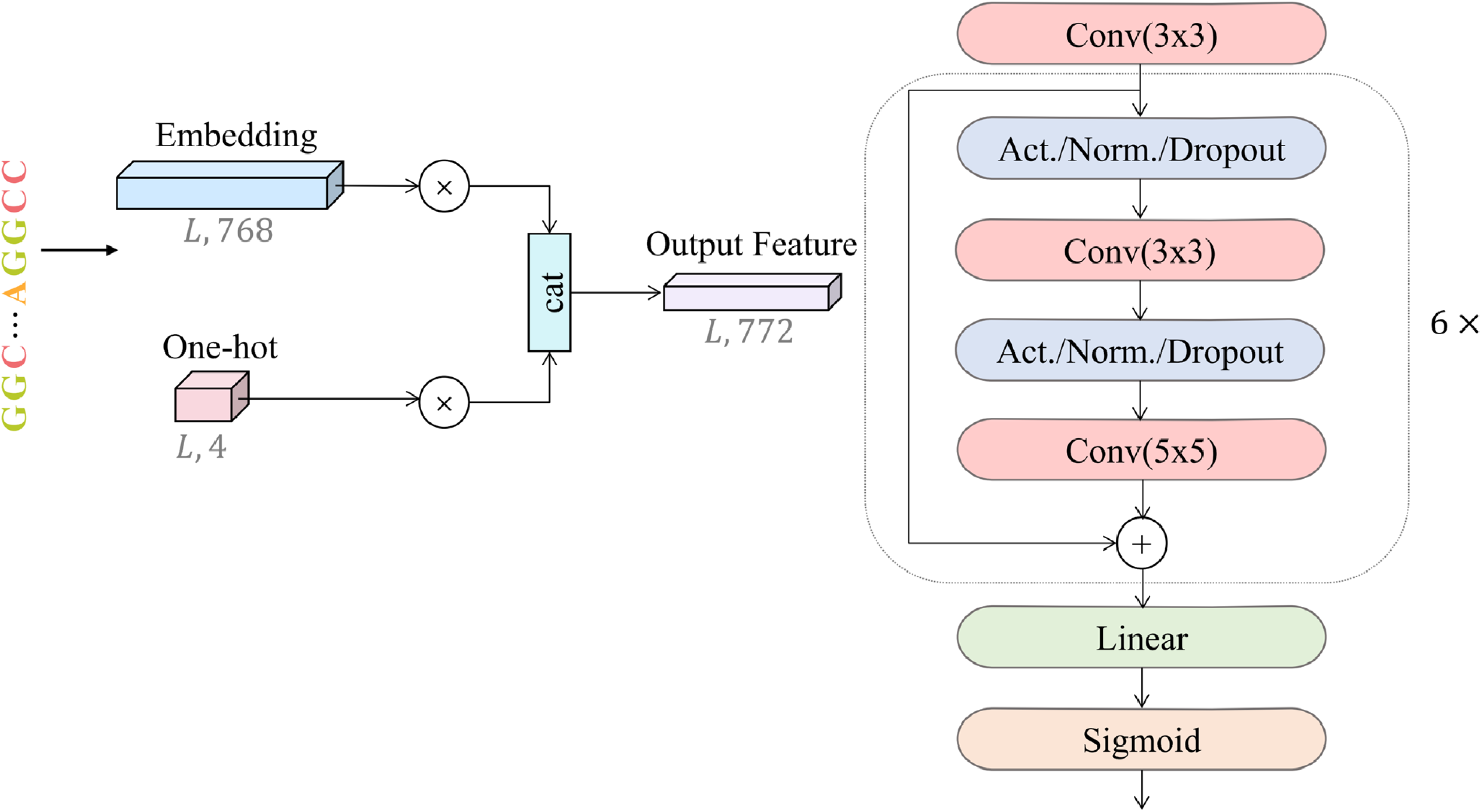
Model architecture for prediction of ribosome loading and mRNA expression levels (RESM-riboload and RESM-mRNAexpr). The model integrates one-hot encoded RNA sequences and final-layer embeddings from the pre-trained RESM model. These features are concatenated and projected into a unified representation, which is passed to a ResNet1D prediction head consisting of six residual blocks. Each residual block contains multiple 1D convolutional layers with kernel sizes of 3 and 5, along with activation, normalization, and dropout. A final linear layer and sigmoid activation are used for regression output. This architecture is utilized consistently across both ribosome loading and expression level prediction tasks, enabling robust evaluation of the model’s generalization capabilities across diverse biological datasets.

