## Supplementary material for "RESM: Capturing sequence and structure encoding of RNAs by mapped transfer learning from ESM (evolutionary scale modeling) protein language model": https://github.com/yikunpku/RESM

### Data details

**Supplementary Table 1. Datasets used for RESM pre-training and downstream tasks.** Overview of datasets and sample sizes across all tasks. RESM was pretrained on RNAcentral and evaluated on zero-shot and supervised benchmarks covering structure, function, interaction, and regulatory prediction tasks.

| Task | Dataset | Number |
|---|---|---|
| Pre-training | RNAcentral 18.0 | 23,510,939 |
| Zero-shot RNA functional classification | Rfam 14.10 | 24,607 |
| | ArchiveII | 3,864 |
| Zero-shot secondary structure prediction | RNA-MSM Dataset | 515 |
| Supervised secondary structure prediction | RNA-MSM Dataset | 515 |
| | bpRNA-1m Dataset | 12,114 |
| | Long RNA Dataset | 14 |
| RNA-RNA interaction prediction | RRI Dataset | 64 |
| Supervised RNA solvent accessibility prediction | RNA-MSM Dataset | 515 |
| Supervised siRNA Activity prediction | Huesken dataset | 2,361 |
| | Mixset dataset | 472 |
| Supervised ribosome loading | Redundant GSM4084997 Dataset | 91,519 |
| | Non-redundant GSM4084997 Dataset | 85,572 |
| Expression levels prediction | Muscle | 1,257 |
| | HEK | 14,410 |
| | PC3 | 12,579 |

### Results

**Supplementary Table 2. Zero-shot evaluation on RNA secondary structure and functional classification tasks.** Models were evaluated under a unified zero-shot setting using attention maps from a fixed head-layer across all RNA sequences. For secondary-structure-prediction evaluation, median and mean ± standard deviation of individual overall F1 scores are reported to capture overall predictive quality and consistency. Functional classification accuracy is assessed using the overlap ratio (OR) on two datasets, Rfam and ArchiveII, with lower values indicating better class separability.

| **Model** | **F1-score (Secondary Structure)↑** | | | **Functional Classifications OR↓** | |
|---|---|---|---|---|---|
| | **Median** | **Mean± Std** | **P-value** | **Rfam** | **ArchiveII** |
| UTR_LM | 0.022 | 0.029 ± 0.032 | 8.00E-29 | 0.202 ± 0.001 | 0.626 ± 0.000 |
| SpliceBERT | 0.034 | 0.043 ± 0.039 | 2.24E-29 | 0.240 ±0.001 | 0.578 ± 0.001 |
| BiRNA-BERT (BPE) | - | - | - | 0.206 ± 0.001 | 0.626 ± 0.003 |
| BiRNA-BERT (NUC) | - | - | - | 0.161 ± 0.000 | 0.454 ± 0.002 |
| RNA-FM | 0.180 | 0.267 ± 0.239 | 1.92E-18 | 0.070 ± 0.000 | 0.375 ± 0.002 |
| RNAErnie | 0.238 | 0.288 ± 0.223 | 3.24E-17 | 0.156 ± 0.000 | 0.514 ± 0.000 |
| RiNALMo (micro) | 0.256 | 0.268 ± 0.184 | 4.45E-19 | 0.168 ± 0.000 | 0.486 ± 0.002 |
| ERNIE-RNA | 0.259 | 0.300 ± 0.234 | 2.74E-15 | 0.089 ± 0.000 | 0.493 ± 0.002 |
| ProtRNA | 0.298 | 0.333 ± 0.236 | 2.93E-17 | 0.081 ± 0.000 | 0.361 ± 0.000 |
| RiNALMo (mega) | 0.315 | 0.342 ± 0.225 | 8.70E-17 | 0.146 ± 0.000 | 0.587 ± 0.001 |
| RiNALMo (giga) | 0.404 | 0.414 ± 0.255 | 2.00E-11 | 0.116 ± 0.000 | 0.600 ± 0.002 |
| AIDO.RNA (650M) | 0.438 | 0.402 ± 0.234 | 3.45E-12 | 0.104 ± 0.000 | 0.395 ± 0.001 |
| RNA-km | 0.473 | 0.492 ± 0.236 | 1.49E-04 | 0.197 ± 0.001 | 0.683 ± 0.001 |
| MP-RNA | 0.507 | 0.507 ± 0.201 | 1.19E-02 | 0.235 ± 0.001 | 0.545 ± 0.001 |
| AIDO.RNA (1.6B) | 0.579 | 0.508 ± 0.263 | 3.43E-04 | 0.110 ± 0.000 | 0.445 ± 0.000 |
| RESM-150M (ours) | 0.596 | 0.521 ± 0.261 | 4.96E-05 | 0.072 ± 0.000 | **0.334 ± 0.002** |
| RNA-MSM | 0.631 | 0.582 ± 0.217 | 9.50E-01 | - | - |
| RESM-650M (ours) | **0.637** | **0.584 ± 0.248** | **-** | **0.065 ± 0.000** | 0.417 ± 0.000 |

**Supplementary Table 3. Zero-shot RNA secondary structure prediction.** Performance on the TS70 test set, evaluated separately on canonical and non-canonical base pairs. The table reports the median and mean ± standard deviation of the F1 scores for each model, along with corresponding P-values from paired comparisons with RESM-650M in this work.

| **Model** | **Canonical (F1-score)** | | | **Non-canonical (F1-score)** | | |
|---|---|---|---|---|---|---|
| | **Median** | **Mean± Std** | **P-value** | **Median** | **Mean± Std** | **P-value** |
| UTR_LM | 0.035 | 0.047 ± 0.053 | 1.48E-29 | 0.000 | 0.002 ± 0.011 | 8.72E-13 |
| SpliceBERT | 0.068 | 0.088 ± 0.085 | 2.23E-29 | 0.000 | 0.013 ± 0.019 | 2.78E-12 |
| RNA-FM | 0.272 | 0.337 ± 0.271 | 1.22E-16 | 0.000 | 0.051 ± 0.092 | 1.28E-10 |
| RNAErnie | 0.320 | 0.368 ± 0.255 | 5.37E-15 | 0.000 | 0.060 ± 0.118 | 6.84E-12 |
| ERNIE-RNA | 0.331 | 0.359 ± 0.262 | 7.40E-15 | 0.000 | 0.031 ± 0.102 | 1.88E-11 |
| RiNALMo (micro) | 0.352 | 0.354 ± 0.231 | 8.61E-17 | 0.000 | 0.044 ± 0.089 | 2.21E-11 |
| ProtRNA | 0.405 | 0.407 ± 0.264 | 1.17E-15 | 0.000 | 0.074 ± 0.133 | 9.54E-10 |
| RiNALMo (mega) | 0.458 | 0.447 ± 0.258 | 8.58E-13 | 0.000 | 0.078 ± 0.117 | 9.37E-11 |
| AIDO.RNA (650M) | 0.520 | 0.477 ± 0.266 | 1.54E-12 | 0.000 | 0.044 ± 0.111 | 5.20E-12 |
| RiNALMo (giga) | 0.539 | 0.504 ± 0.286 | 1.12E-09 | 0.000 | 0.098 ± 0.152 | 3.75E-08 |
| RNA-km | 0.611 | 0.575 ± 0.259 | 1.36E-04 | 0.085 | 0.165 ± 0.201 | 7.49E-04 |
| MP-RNA | 0.631 | 0.608 ± 0.219 | 5.70E-02 | 0.000 | 0.098 ± 0.136 | 5.47E-08 |
| AIDO.RNA (1.6B) | 0.684 | 0.593 ± 0.293 | 1.14E-03 | 0.209 | 0.189 ± 0.191 | 1.14E-03 |
| RESM-150M (ours) | 0.700 | 0.620 ± 0.280 | 1.80E-03 | 0.114 | 0.183 ± 0.207 | 3.57E-04 |
| RNA-MSM | **0.756** | **0.677 ± 0.232** | 9.15E-01 | 0.200 | 0.225 ± 0.201 | 2.86E-01 |
| RESM-650M (ours) | 0.743 | 0.674 ± 0.267 | - | **0.236** | **0.253 ± 0.239** | - |

**Supplementary Table 4. Performance of supervised RNA secondary-structure prediction using attention-based representations on the TS70 dataset, as compared to other models, according to the F1 score**. Supervised prediction was performed using attention maps aggregated from all layers and attention heads of each model as input to a downstream task-specific network. Model performance was evaluated across the dataset in terms of median F1 score and mean ± standard deviation of individual F1 scores for each RNA. P-values indicate the statistical significance of paired comparisons with RESM-basepair-650M.

| **Model** | **F1-score** | | **P-value** |
|---|---|---|---|
| | **Median** | **Mean ± Std** | |
| ^Rfold | 0.252 | 0.326 ± 0.275 | 2.54E-16 |
| *ProtRNA | 0.372 | 0.382 ± 0.265 | 5.29E-17 |
| *ERNIE-RNA | 0.386 | 0.428 ± 0.281 | 4.31E-13 |
| ^RNADIFFfold | 0.432 | 0.469 ± 0.242 | 7.34E-12 |
| *RiNALMo (giga) | 0.533 | 0.506 ± 0.246 | 5.03E-12 |
| *AIDO.RNA (1.6B) | 0.580 | 0.523 ± 0.242 | 6.13E-11 |
| RESM-zero-shot-150M (ours) | 0.596 | 0.521 ± 0.261 | 1.80E-12 |
| *RNAErnie | 0.601 | 0.546 ± 0.260 | 3.77E-06 |
| †RNAstructure | 0.606 | 0.571 ± 0.271 | 4.99E-04 |
| †CONTRAfold | 0.628 | 0.576 ± 0.265 | 3.99E-04 |
| RESM-zero-shot-650M (ours) | 0.637 | 0.584 ± 0.248 | 5.27E-08 |
| †EternaFold | 0.638 | 0.592 ± 0.245 | 1.15E-03 |
| ^KnotFold | 0.651 | 0.568 ± 0.248 | 1.31E-06 |
| †LinearFold | 0.653 | 0.580 ± 0.273 | 1.09E-03 |
| †RNAfold | 0.656 | 0.586 ± 0.273 | 3.27E-03 |
| ^BPfold | 0.667 | 0.628 ± 0.207 | 5.68E-03 |

| | | | |
|---|---|---|---|
| ^UFold | 0.667 | 0.606 ± 0.223 | 6.56E-04 |
| *RNA-FM | 0.679 | 0.634 ± 0.242 | 1.34E-02 |
| ^MXfold2 | 0.696 | 0.648 ± 0.217 | 7.99E-02 |
| ^SPOT-RNA | 0.699 | 0.629 ± 0.231 | 5.56E-03 |
| *RNA-MSM | 0.721 | 0.662 ± 0.211 | 1.49E-01 |
| ^SPOT-RNA2 | 0.743 | 0.678 ± 0.211 | 5.36E-01 |
| RESM-basepair-650M (ours) | 0.746 | **0.701 ± 0.180** | - |
| RESM-basepair-150M (ours) | **0.748** | 0.686 ± 0.190 | 2.20E-01 |

Note: Models with "†" denote thermodynamic or probabilistic methods; models with "^" denote deep learning methods; models with "*" denote RNA language models with trained RNA secondary-structure prediction header, whose prediction leader networks and training weights are available from the original reported literature. "RESM-zero-shot-150M" and "RESM-zero-shot-650M" show the zero-shot results, consistent with the results of "RESM-150M" and "RESM-650M" in Supplementary Table 2, respectively. Models with an underlined name have training set overlap with TS70. And limited by the sequence requirements of Rfold, UFold and SPOT-RNA2 methods, they only show the results of 66, 66, and 67 PDBs from the TS70 dataset, respectively.

**Supplementary Table 5. Performance of supervised RNA secondary-structure prediction using attention-based representations on the TS70 dataset, as compared to other models, according to canonical and non-canonical base pairs, separately.** Results are reported separately for canonical and non-canonical base pairs. F1 scores are presented as the median and the mean ± standard deviation. P-values indicate the statistical significance of paired comparisons with RESM-basepair-650M.

| Model | Canonical (F1-score) | | | Non-canonical (F1-score) | | |
|---|---|---|---|---|---|---|
| | Median | Mean ± Std | P-value | Median | Mean ± Std | P-value |
| ^Rfold | 0.333 | 0.376 ± 0.304 | 2.17E-15 | 0.000 | 0.000 ± 0.000 | 7.74E-13 |
| *ProtRNA | 0.453 | 0.446 ± 0.297 | 1.67E-15 | 0.000 | 0.020 ± 0.071 | 3.51E-12 |
| *ERNIE-RNA | 0.529 | 0.506 ± 0.313 | 6.11E-11 | 0.000 | 0.002 ± 0.013 | 1.01E-12 |
| ^RNADIFFfold | 0.548 | 0.548 ± 0.256 | 5.86E-10 | 0.000 | 0.132 ± 0.189 | 4.39E-06 |
| *RiNALMo (giga) | 0.648 | 0.584 ± 0.273 | 9.14E-10 | 0.000 | 0.000 ± 0.000 | 7.63E-13 |
| *AIDO.RNA (1.6B) | 0.667 | 0.606 ± 0.268 | 2.28E-08 | 0.000 | 0.000 ± 0.000 | 7.63E-13 |
| RESM-zero-shot-150M (ours) | 0.700 | 0.620 ± 0.280 | 1.55E-08 | 0.114 | 0.183 ± 0.207 | 4.73E-05 |
| *RNAErnie | 0.692 | 0.622 ± 0.284 | 8.61E-05 | 0.000 | 0.000 ± 0.000 | 7.63E-13 |
| †RNAstructure | 0.743 | 0.643 ± 0.291 | 2.05E-03 | 0.000 | 0.000 ± 0.000 | 7.63E-13 |
| †CONTRAfold | 0.721 | 0.649 ± 0.280 | 1.56E-03 | 0.000 | 0.000 ± 0.000 | 7.63E-13 |
| RESM-zero-shot-650M (ours) | 0.743 | 0.674 ± 0.267 | 5.84E-05 | 0.236 | 0.253 ± 0.239 | 1.11E-01 |
| †EternaFold | 0.731 | 0.668 ± 0.258 | 5.42E-03 | 0.000 | 0.000 ± 0.000 | 7.63E-13 |
| ^KnotFold | 0.711 | 0.646 ± 0.265 | 5.68E-05 | 0.000 | 0.106 ± 0.171 | 9.61E-09 |
| †LinearFold | 0.725 | 0.656 ± 0.290 | 5.43E-03 | 0.000 | 0.000 ± 0.000 | 7.63E-13 |
| †RNAfold | 0.772 | 0.660 ± 0.291 | 1.03E-02 | 0.000 | 0.000 ± 0.000 | 7.63E-13 |
| ^BPfold | 0.757 | 0.714 ± 0.212 | 5.02E-02 | 0.000 | 0.000 ± 0.000 | 7.63E-13 |
| ^UFold | 0.764 | 0.677 ± 0.230 | 2.00E-03 | 0.000 | 0.068 ± 0.191 | 4.31E-08 |

| | | | | | | |
|---|---|---|---|---|---|---|
| *RNA-FM | 0.786 | 0.708 ± 0.246 | 4.18E-02 | 0.000 | 0.142 ± 0.221 | 1.08E-06 |
| ^MXfold2 | 0.805 | 0.736 ± 0.225 | 3.39E-01 | 0.000 | 0.000 ± 0.000 | 7.63E-13 |
| ^SPOT-RNA | 0.788 | 0.705 ± 0.233 | 2.18E-02 | 0.000 | 0.162 ± 0.218 | 7.27E-06 |
| *RNA-MSM | 0.811 | 0.742 ± 0.219 | 3.99E-01 | 0.115 | 0.195 ± 0.225 | 1.08E-03 |
| ^SPOT-RNA2 | **0.828** | 0.761 ± 0.211 | 9.86E-01 | 0.167 | 0.237 ± 0.246 | 7.22E-02 |
| RESM-basepair-650M (ours) | 0.806 | **0.766 ± 0.180** | - | **0.273** | **0.288 ± 0.275** | - |
| RESM-basepair-150M (ours) | 0.820 | 0.752 ± 0.192 | 2.89E-01 | 0.250 | 0.269 ± 0.266 | 2.15E-01 |

Note: Models with “†” denote thermodynamic or probabilistic methods; models with “^” denote deep learning methods; models with “*” denote RNA language models with trained RNA secondary-structure prediction header, whose prediction leader networks and training weights are available from the original reported literature. “RESM-zero-shot-150M” and “RESM-zero-shot-650M” show zero-shot results, consistent with the results of “RESM-150M” and “RESM-650M” in Supplementary Table 2, respectively. Models with an underlined name have training set overlap with TS70. And limited by the sequence requirements of Rfold, UFold and SPOT-RNA2 methods, they only show the results of 66, 66, and 67 PDBs from the TS70 dataset, respectively.

**Supplementary Table 6. The long RNA dataset.** Detailed information of 14 non-redundant long RNA structures (from 1,256 to 3,705 nucleotides) used for evaluating RESM's and RESM-basepair's generalization capability beyond their training length limit of 1,024. In detail, RNA chains with a sequence length greater than 1,024 and a base count greater than 500 (bases with a determined structure) in the PDB structure were first collected from the RNA3DB structure database. Subsequently, based on sequence homology (identity $< 80\%$) and PDB structural similarity (TMscore $< 0.45$). When predicting secondary structures, the full-length RNA sequences were input into each method.

| **PDB_Chain** | **Full-length** | **Num. of bases in PDB** | **Resolution (Å)** | **RNA Type** |
|---|---|---|---|---|
| 7qh7_A | 1256 | 1256 | 2.89 | 16S rRNA |
| 6z1p_Bb | 1395 | 1385 | 3.7 | SSU rRNA |
| 6nu2_A | 1472 | 1472 | 3.9 | 16S rRNA |
| 3jcs_2 | 1527 | 1119 | 2.8 | 26S delta rRNA |
| 8ova_BB | 1536 | 1119 | 2.47 | LSUb rRNA |
| 7tql_2 | 1656 | 1656 | 3.2 | 18S rRNA |
| 8d8j_a | 1713 | 1183 | 3.8 | 15S rRNA |
| 6wdr_2 | 1910 | 1647 | 3.7 | 20S rRNA |
| 8ova_BA | 1920 | 1604 | 2.47 | LSUa rRNA |
| 7ase_0 | 2319 | 2150 | 3.33 | 18S rRNA |
| 5oql_1 | 2568 | 1350 | 3.2 | 35S rRNA |
| 5mrc_A | 3296 | 2709 | 3.25 | 21S rRNA |
| 6ywx_A | 3464 | 2838 | 3.1 | 23S rRNA |
| 7nwh_5 | 3705 | 3543 | 4.1 | 28S rRNA |

**Supplementary Table 7. Performance of RNA secondary-structure prediction on long RNA dataset, according to the F1 score.** The evaluation contained zero-shot prediction and supervised prediction, whose processes were consistent with Supplementary Table 2 and Supplementary Table 4, respectively. And in the zero-shot prediction process, the threshold was first evaluated on the VL46 dataset and then tested on the long RNA dataset. P-values were calculated based on the results of RESM-650M (zero-shot prediction) or RESM-basepair-650M (supervised prediction).

| Task | Model [Nums. Of RNA] | F1-score | | P-value |
|---|---|---|---|---|
| | | Median | Mean ± Std | |
| Zero-shot | UTR_LM [14] | 0.000 | 0.002 ± 0.003 | 1.53E-10 |
| | LucaOne [14] | 0.064 | 0.069 ± 0.027 | 2.47E-10 |
| | MP-RNA [14] | 0.173 | 0.177 ± 0.055 | 2.10E-08 |
| | RiNALMo (micro) [9] | 0.245 | 0.24 ± 0.059 | 2.32E-05 |
| | AIDO.RNA (650M) [14] | 0.304 | 0.303 ± 0.071 | 7.18E-06 |
| | RiNALMo (giga) [8] | 0.394 | 0.412 ± 0.079 | 2.12E-04 |
| | RiNALMo (mega) [9] | 0.396 | 0.396 ± 0.071 | 3.41E-05 |
| | RESM-150M (ours) [14] | 0.439 | 0.434 ± 0.122 | 1.01E-09 |
| | AIDO.RNA (1.6B) [14] | 0.464 | 0.494 ± 0.099 | 3.71E-03 |
| | RNA-km [14] | 0.513 | 0.422 ± 0.239 | 1.03E-01 |
| | RESM-650M (ours) [14] | **0.552** | **0.532 ± 0.113** | - |
| Supervised | ^BPfold [14] | 0.000 | 0.000 ±0.000 | 5.55E-13 |
| | ^Rfold [9] | 0.000 | 0.012 ± 0.029 | 4.52E-08 |
| | ^UFold [14] | 0.013 | 0.035 ± 0.048 | 1.03E-13 |
| | ^KnotFold [14] | 0.107 | 0.097 ± 0.052 | 8.18E-12 |
| | ^SPOT-RNA [14] | 0.183 | 0.184 ± 0.045 | 3.28E-10 |
| | †LinearFold [14] | 0.210 | 0.225 ± 0.068 | 2.40E-08 |
| | †RNAfold [14] | 0.241 | 0.245 ± 0.064 | 7.86E-09 |
| | †RNAstructure [14] | 0.276 | 0.259 ± 0.078 | 4.48E-09 |
| | ^MXfold2 [14] | 0.285 | 0.285 ± 0.067 | 4.37E-08 |
| | †EternaFold [14] | 0.288 | 0.304 ± 0.089 | 6.61E-09 |
| | †CONTRAfold [14] | 0.335 | 0.304 ± 0.091 | 3.17E-08 |
| | ^SPOT-RNA2 [5] | 0.336 | 0.332 ± 0.053 | 7.17E-04 |
| | RESM-basepair-150M (ours) [14] | 0.559 | 0.551 ± 0.076 | 2.93E-10 |
| | RESM-basepair-650M (ours) [14] | **0.609** | **0.598 ± 0.080** | - |
| | RESM-basepair-150M (ours) [5] | 0.565 | 0.558 ± 0.034 | - |
| | RESM-basepair-650M (ours) [5] | **0.619** | **0.602 ± 0.035** | - |

Note: Some methods listed in Supplementary Table 2 and Supplementary Table 4 were not included in the comparison due to limitations on the maximum input length or the excessive computational resource consumption. Models with "†" denote thermodynamic or probabilistic methods, and models with "^" denote deep learning methods. RNA language models methods with trained RNA secondary-structure prediction header listed in the Supplementary Table 4 (excluding our methods) have limitations on the maximum input length. The square brackets behind the model indicate the amount of RNA that each method can calculate. For the supervised task, since the multiple sequence alignment of long sequences was particularly time-consuming, only the prediction results of the SPOT-RNA2 method for five RNAs shorter than 2000 were shown here. And for comparison, we calculated the results of RESM-basepair for these five RNAs (gray background).

**Supplementary Table 8. Performance on zero-shot RNA–RNA interaction prediction.** Comparison of 23 predictors of inter-RNA base pairs on 64 complexes of RNA structures unseen by SPOT-RNA and SPOT-RNA2. All single-chain structures in the test set have the structural similarity score TM-Score < 0.3 compared to the monomeric structures used in training and validating SPOT-RNA and SPOT-RNA2 methods.

| **Methods** | **PPV/ Precision** | **Sensitivity/Recall** | **Overall F1-Score** | **MCC** | **Individual F1-Score Median ± Std** |
|---|---|---|---|---|---|
| GUUGle | 0.159 | 0.105 | 0.126 | 0.128 | 0.000 ± 0.241 |
| RIsearch | 0.176 | 0.254 | 0.208 | 0.210 | 0.000 ± 0.319 |
| RNAplex-cA# | 0.477 | 0.149 | 0.227 | 0.266 | 0.000 ± 0.296 |
| RNAaliduplex | 0.484 | 0.149 | 0.228 | 0.268 | 0.000 ± 0.296 |
| DuplexFold | 0.182 | 0.458 | 0.260 | 0.287 | 0.200 ± 0.344 |
| RNAplex-c | 0.189 | 0.445 | 0.265 | 0.289 | 0.216 ± 0.343 |
| RNAduplex | 0.195 | 0.495 | 0.280 | 0.309 | 0.229 ± 0.329 |
| RNAup | 0.282 | 0.360 | 0.316 | 0.317 | 0.000 ± 0.290 |
| RNAplex-a | 0.364 | 0.287 | 0.321 | 0.322 | 0.000 ± 0.331 |
| AccessFold | 0.317 | 0.504 | 0.389 | 0.398 | 0.349 ± 0.315 |
| PETcoFold# | 0.434 | 0.368 | 0.398 | 0.399 | 0.377 ± 0.352 |
| NUPACK | 0.442 | 0.479 | 0.460 | 0.459 | 0.474 ± 0.355 |
| RNAfoldc | 0.381 | 0.579 | 0.460 | 0.469 | 0.446 ± 0.318 |
| MXfold2c | 0.389 | 0.601 | 0.472 | 0.482 | 0.448 ± 0.320 |
| UFoldc | 0.419 | 0.542 | 0.473 | 0.476 | 0.455 ± 0.302 |
| bifold | 0.409 | 0.563 | 0.474 | 0.479 | 0.454 ± 0.325 |
| PairFold | 0.424 | 0.559 | 0.482 | 0.486 | 0.500 ± 0.339 |
| IntaRNA2.0 | 0.540 | 0.440 | 0.485 | 0.486 | 0.405 ± 0.377 |
| RNAmultifold | 0.420 | 0.579 | 0.487 | 0.492 | 0.494 ± 0.325 |
| RNAcoFold | 0.421 | 0.579 | 0.488 | 0.493 | 0.494 ± 0.325 |
| EternaFoldc | 0.437 | **0.618** | 0.512 | 0.519 | 0.498 ± 0.314 |
| SPOT-RNA2c# | 0.569 | 0.537 | 0.553 | 0.552 | 0.529 ± 0.329 |
| SPOT-RNAc | 0.606 | 0.548 | 0.576 | 0.576 | 0.579 ± 0.323 |
| RESM-basepair-650Mc ($X^2$) | 0.695 | 0.498 | 0.580 | 0.588 | 0.644 ± 0.368 |
| RESM-basepair-150Mc ($X^8$) | **0.733** | 0.520 | **0.609** | **0.617** | **0.686 ± 0.355** |

Note: The overall F1 score is harmonic mean of precision and recall for all RNA pairs. PPV denotes positive predictive value. MCC denotes Matthews' correlation coefficient. "#" denotes the use of evolution information.

Methods with an ending of “c” indicate the use of chain concatenation for RNA-RNA interaction prediction. Median F1 means the median F1 value of single RNA. For RESM-basepair, the evaluation of the linkers corresponding to the best individual median F1 score (“$X^8$” for 150M and “$X^2$” for 650M) was shown here.

**Supplementary Table 9. Performance on zero-shot RNA–RNA interaction prediction by RESM-basepair-150M with and without linker sequences.** Performance is reported using precision (PPV), recall (sensitivity), overall F1 score, Matthews correlation coefficient (MCC), and the median ± standard deviation of individual F1 scores across samples. "None" indicates no linker was used between input RNA pairs. Each $X^n$ denotes a linker sequence composed by concatenating *n* repetitions of "X".

| Linker | PPV/Precision | Sensitivity/Recall | Overall F1-Score | MCC | Individual F1-Score Median ± Std |
|---|---|---|---|---|---|
| *None* | 0.687 | 0.523 | 0.594 | 0.599 | 0.623 ± 0.345 |
| $X^1$ | 0.698 | 0.520 | 0.596 | 0.601 | 0.628 ± 0.356 |
| $X^2$ | 0.702 | 0.518 | 0.596 | 0.602 | 0.653 ± 0.358 |
| $X^3$ | 0.707 | 0.523 | 0.601 | 0.608 | 0.641 ± 0.360 |
| $X^4$ | 0.728 | **0.534** | **0.616** | 0.623 | 0.653 ± 0.344 |
| $X^5$ | 0.730 | 0.517 | 0.605 | 0.614 | 0.653 ± 0.356 |
| $X^6$ | 0.731 | 0.533 | **0.616** | 0.624 | 0.651 ± 0.344 |
| $X^7$ | **0.743** | 0.527 | **0.616** | **0.625** | 0.664 ± 0.351 |
| $X^8$ | 0.733 | 0.520 | 0.609 | 0.617 | **0.686** ± **0.355** |
| $X^9$ | 0.738 | 0.513 | 0.605 | 0.615 | 0.679 ± 0.364 |
| $X^{10}$ | 0.735 | 0.516 | 0.606 | 0.615 | 0.679 ± 0.359 |
| $X^{11}$ | 0.736 | 0.518 | 0.608 | 0.617 | 0.653 ± 0.358 |
| $X^{12}$ | 0.735 | 0.521 | 0.610 | 0.619 | 0.637 ± 0.353 |

**Supplementary Table 10. Performance of zero-shot RNA–RNA interaction prediction by RESM-basepair-650M with and without linker sequences.** Performance metrics include precision (PPV), recall (sensitivity), overall F1 score, Matthews correlation coefficient (MCC), and the median ± standard deviation of individual F1 scores across test samples. “None” indicates that no linker was used between paired input RNAs. Each linker type “$X^{n}$” represents a sequence of *n* repetitions of the token “X”. The results demonstrate that moderate-length linkers (e.g., $X^{4}$) can improve individual F1 scores, while longer linkers (e.g., $X^{12}$) yield the best overall F1 score and MCC.

| Linker | PPV/Precision | Sensitivity/Recall | Overall F1-Score | MCC | Individual F1-Score Median ± Std |
|---|---|---|---|---|---|
| *None* | 0.668 | 0.499 | 0.571 | 0.577 | 0.636 ± 0.364 |
| $X^{1}$ | 0.689 | 0.498 | 0.578 | 0.585 | 0.636 ± 0.369 |
| $X^{2}$ | 0.695 | 0.498 | 0.580 | 0.588 | **0.644** ± **0.368** |
| $X^{3}$ | 0.684 | 0.500 | 0.578 | 0.584 | 0.639 ± 0.371 |
| $X^{4}$ | 0.680 | 0.501 | 0.577 | 0.583 | **0.644** ± **0.373** |
| $X^{5}$ | 0.672 | 0.501 | 0.574 | 0.580 | 0.637 ± 0.367 |
| $X^{6}$ | 0.685 | 0.498 | 0.577 | 0.583 | 0.639 ± 0.370 |
| $X^{7}$ | 0.692 | 0.499 | 0.580 | 0.587 | 0.636 ± 0.374 |
| $X^{8}$ | 0.694 | 0.499 | 0.581 | 0.588 | 0.628 ± 0.365 |
| $X^{9}$ | 0.701 | **0.504** | 0.586 | 0.594 | 0.630 ± 0.359 |
| $X^{10}$ | 0.711 | 0.498 | 0.586 | 0.595 | 0.620 ± 0.358 |
| $X^{11}$ | **0.725** | 0.497 | 0.590 | 0.600 | 0.602 ± 0.357 |
| $X^{12}$ | **0.725** | 0.502 | **0.593** | **0.603** | 0.603 ± 0.352 |

**Supplementary Table 11. Performance of zero-shot RNA–RNA interaction prediction of RNA language models with secondary-structure prediction heads.** Models were evaluated on 62 RNA complexes, excluding 2 complexes with sequences exceeding 510 nucleotides. All linker sequences were set to *None* in this evaluation.

| Method | PPV/Precision | Sensitivity/Recall | Overall F1-Score | MCC | Individual F1-Score Median ± Std |
|---|---|---|---|---|---|
| *RiNALMo (giga) | 0.444 | 0.285 | 0.347 | 0.354 | 0.000 ± 0.329 |
| *ERNIE-RNA | 0.480 | 0.327 | 0.389 | 0.395 | 0.049 ± 0.347 |
| *AIDO.RNA (1.6B) | 0.517 | 0.374 | 0.434 | 0.439 | 0.306 ± 0.330 |
| *ProtRNA | 0.630 | 0.345 | 0.446 | 0.465 | 0.327 ± 0.329 |
| *RNAErnie | 0.419 | **0.561** | 0.480 | 0.484 | 0.475 ± 0.313 |
| *RNA-FM | 0.527 | 0.548 | 0.537 | 0.536 | 0.484 ± 0.317 |
| RESM-basepair-650M (ours) | 0.692 | 0.527 | 0.599 | 0.604 | **0.644 ± 0.360** |
| RESM-basepair-150M (ours) | **0.705** | 0.534 | **0.607** | **0.613** | 0.632 ± 0.349 |

Note: Models with "*" denote RNA language models with trained RNA secondary-structure prediction header, whose prediction leader networks and training weights are available from the original reported literature.

**Supplementary Table 12**. **Supervised RNA solvent accessibility prediction on the RNA-MSM benchmark by RESM-ASA as compared to other methods.** Pearson correlation coefficient (PCC, higher is better) and mean absolute error (MAE in Å$^2$, lower is better) are reported for each model. RESM-650M achieves the best performance, outperforming RNA-FM, RNA-MSM, RNAsnap2, SeqProf and one-hot encoding approaches.

| Model | PCC ↑ | MAE ↓ |
|---|---|---|
| OneHot | 0.387 | 31.853 |
| OneHot+SeqProf | 0.421 | 34.444 |
| RNA-FM | 0.379 | 35.164 |
| RNAsnap2_seq | 0.401 | 33.097 |
| RNAsnap2_pro | 0.404 | 32.846 |
| RNA-MSM | 0.433 | 31.889 |
| RESM-ASA-150M (ours) | 0.453 | 31.406 |
| RESM-ASA-650M (ours) | **0.482** | **30.981** |

**Supplementary Table 13. Performance on supervised siRNA activity prediction across benchmark models.** Performance was evaluated using three metrics: area under the ROC curve (AUC_ROC), area under the precision-recall curve (AUC_PRC), and F1 score. RESM-siRNA-150M achieves the highest performance across all metrics, surpassing previous methods as shown.

| Model | AUC_ROC | AUC_PRC | F1-Score |
|---|---|---|---|
| Monopoli-RF | 0.744 | 0.739 | 0.700 |
| OligoWalk | 0.708 | 0.699 | 0.626 |
| siRNAPred | 0.578 | 0.618 | 0.525 |
| iScore | 0.746 | 0.756 | 0.220 |
| s-Biopredsi | 0.737 | 0.756 | 0.594 |
| DSIR | 0.748 | 0.764 | 0.695 |
| OligoFormer | 0.815 | 0.814 | 0.769 |
| RESM-siRNA-150M (ours) | **0.820** | **0.827** | **0.797** |
| RESM-siRNA-650M (ours) | 0.816 | 0.815 | 0.774 |

**Supplementary Table 14. Performance on supervised ribosome loading prediction across random and human-derived 5'UTRs.** Spearman's rank correlation coefficients were reported for each model on two test sets: synthetic random 5'UTR sequences ("Random") and human-derived 5'UTRs ("Human"). Upper section shows results from models fine-tuned on the non-redundant dataset, while the lower section compares performance when fine-tuned on the original (redundant) dataset. RESM-riboload-150M achieves the highest performance across both settings, with further gains observed when training on the non-redundant dataset.

| Model | Spearman R | |
|---|---|---|
| | Random | Human |
| *Fine-tuning on original (redundant) dataset* | | |
| RNABert | 0.883 | 0.783 |
| RNA-FM | 0.891 | 0.830 |
| Framepool | 0.883 | 0.823 |
| Optimus | 0.902 | 0.832 |
| MTrans | 0.903 | 0.841 |
| UTR_LM | 0.911 | 0.847 |
| RESM-riboload-150M (ours) | **0.941** | **0.875** |
| *Fine-tuning on non-redundant dataset* | | |
| UTR_LM | 0.897 | 0.837 |
| RESM-riboload-150M (ours) | **0.939** | **0.872** |
| RESM-riboload-650M (ours) | 0.936 | 0.865 |

**Supplementary Table 15. Performance on supervised mRNA expression-level prediction across human tissues and cell lines by RESM-mRNAexpr.** Spearman's rank correlation coefficients were reported for each model on three datasets: human skeletal muscle, prostate cancer (PC3) cells, and human embryonic kidney (HEK) 293T cells. The upper section presents results from models fine-tuned on truncated 5'UTR sequences (first 100 nucleotides), while the lower section reports performance when full-length RNA sequences were used as input.

| Model | Spearman R | | |
|---|---|---|---|
| | **Muscle** | **HEK** | **PC3** |
| *Fine-tuning on Truncated (100 nt) dataset* | | | |
| Optimus | 0.153 | 0.182 | 0.191 |
| Framepool | 0.282 | 0.372 | 0.330 |
| RNA-FM | 0.302 | 0.222 | 0.201 |
| MTrans | 0.481 | 0.471 | 0.480 |
| RNABert | 0.602 | 0.472 | 0.450 |
| Cao-RF | 0.641 | 0.570 | 0.590 |
| UTR_LM | 0.661 | 0.651 | 0.630 |
| RESM-150M (ours) | **0.692** | **0.661** | **0.645** |
| *Fine-tuning on full-length dataset* | | | |
| UTR_LM | 0.612 | 0.505 | 0.549 |
| RESM-mRNA-expr-150M (ours) | **0.701** | 0.598 | 0.634 |
| RESM-mRNA-expr-650M (ours) | 0.660 | **0.629** | **0.680** |

#### Pretraining details

For model pretraining, we adopted the AdamW optimizer with a learning rate of 1e-3, weight decay of 1e-4, and $\beta_1 = 0.9$, $\beta_2 = 0.98$. To stabilize early-stage training, we employed a warmup linear learning rate scheduler with 1,600 warmup steps, followed by a linear decay schedule. All models were trained using masked language modeling (MLM) as the training objective.

We followed the ESM-2 architecture exactly, adapting its design to the RNA domain without altering core architectural components. The masking strategy was consistent with standard MLM settings: 15% of tokens were randomly masked during training. Importantly, instead of using an RNA-specific vocabulary for prediction, we adopted the protein vocabulary from ESM-2 as the decoding target.

The RESM-150M model was trained for approximately 460k steps on 8 NVIDIA Tesla A100 80GB GPUs, while the larger RESM-650M model was trained for approximately 100k steps on 256 Huawei Ascend D910B GPUs to accommodate its greater parameter count and computational demand.

### Downstream task details

##### Supervised secondary structure prediction and Zero-shot RNA-RNA interaction prediction

**Supplementary Table 16.** The model hyperparameters for supervised secondary-structure prediction used in RESM-150M-basepair or RESM-650M-basepair ensembles. Each model variant shares the same architectural design as SPOT-RNA. $N_A$ denotes the number of the ResNet blocks; $N_{BL}$ is the number of 2D-LSTM layers; $N_B$, represents the number of the fully connected blocks; $D_{RES}$, $D_{BL}$ and $D_{FC}$ refer to the layer depths in ResNet blocks, 2D-LSTM cells (number of nodes per direction), and fully connected layers, respectively. The dilation factor is specified where applicable. Model4 incorporates a dilation factor of $2^{i\%5}$, following the dilated convolution strategy described in the original SPOT-RNA framework.

| Model | Num. blocks/layers | | | Depth of Layer | | | Dilation Factor |
|---|---|---|---|---|---|---|---|
| | $N_A$ | $N_{BL}$ | $N_B$ | $D_{RES}$ | $D_{BL}$ | $D_{FC}$ | |
| Model0 | 16 | - | 2 | 48 | - | 512 | - |
| Model1 | 20 | - | 1 | 64 | - | 512 | - |
| Model2 | 30 | - | 1 | 64 | - | 512 | - |
| Model3 | 30 | 1 | - | 64 | 200 | - | - |
| Model4 | 30 | - | 1 | 64 | - | 512 | $2^{i\%5}$ |

**Supplementary Table 16.** The locations of model weights for supervised secondary-structure prediction heads of compared RNA language models.

| Model | Weight | URL |
|---|---|---|
| *ERNIE-RNA | ERNIER-RNA_ss_prediction.pt | https://github.com/Bruce-ywj/ERNIE-RNA |
| *RiNALMo (giga) | rinalmo_giga_ss_bprna_ft.pt | https://github.com/lbcb-sci/RiNALMo |
| *AIDO.RNA (1.6B) | model.ckpt | https://github.com/genbio-ai/AIDO |
| *RNAErnie | model_state.pdparams (bpRNA1m) | https://github.com/CatIIIIIIII/RNAErnie |
| *RNA-FM | RNA-FM-ResNet_PDB-TR1.pth | https://github.com/ml4bio/RNA-FM |
| *ProtRNA | ssHead_RF_bprna.h5 | https://github.com/roxie-zhang/ProtRNA |
| *RNA-MSM | rna-msm_attention.pt | https://github.com/yikunpku/RNA-MSM |

Note: Models with "*" denote RNA language models with trained RNA secondary-structure prediction header, whose prediction leader networks and training weights are available from the original reported literature.

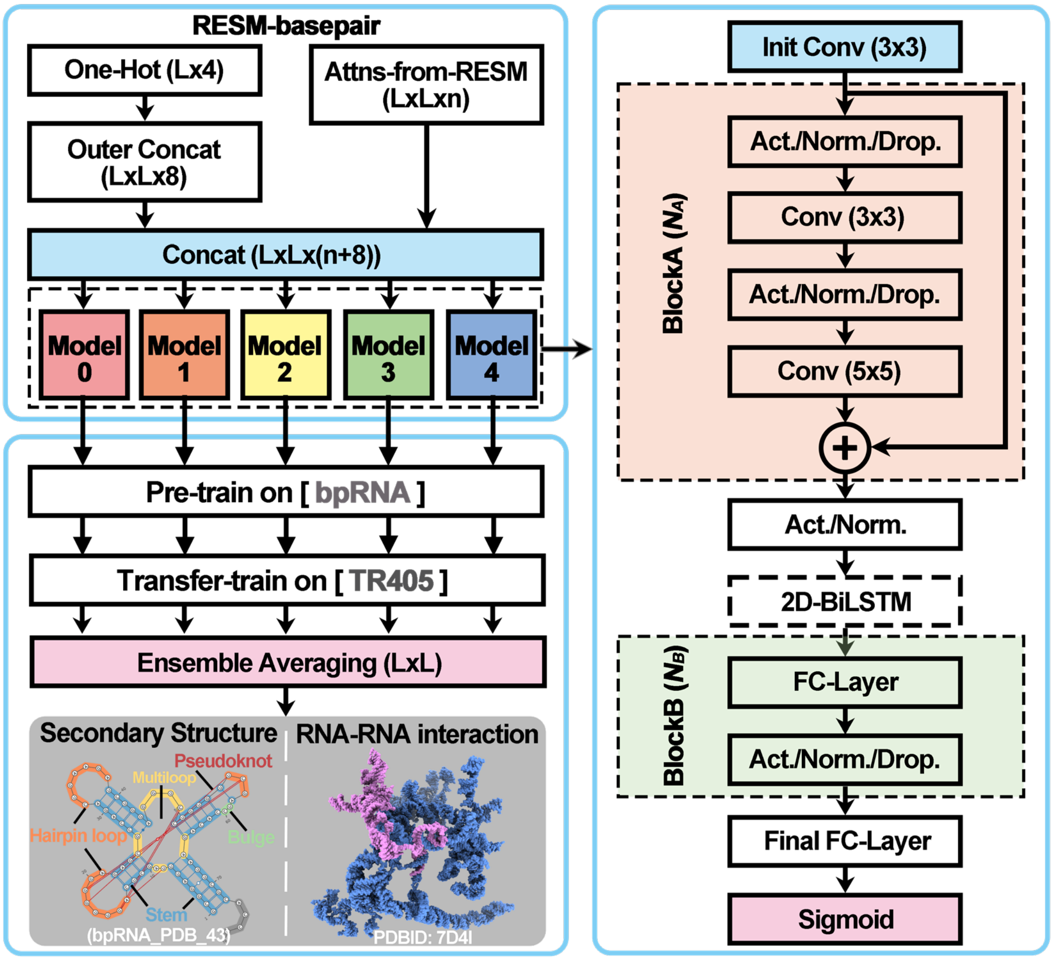


**Supplementary Fig. 1.** Generalized model architecture and training process of RESM-basepair. The full RESM-basepair model consists of five independently trained (including pre-training on the bpRNA dataset and transfer training on the TR405 dataset) models with similar network structure (shown on the right), whose outputs are ensemble averaged to obtain the final prediction. The input is made of two parts, one is the one-hot encoding feature with dimension $L \times 4$ and the other is RESM attention maps with dimension $L \times L \times n$, where $L$ is the sequence length of the target RNA and $n$ is the number of attention maps, which is 600 for the RESM-150M model and 660 for the RESM-650M model. "Outer Concat" indicates the outer concatenation operation; "Conv" indicates the convolutional layer; "Act." indicates the activation function; "Norm." indicates the normalization function; "Drop." indicates the dropout layer; "FC" indicates the fully-connected layer and "Sigmoid" indicates the sigmoid function. The module "BlockA" consists of $N_A$ blocks and the module "BlockB" consists of $N_B$ blocks. The "2D-BiLSTM" layer is only used in "Model 3".

**Supervised RNA solvent accessibility prediction**

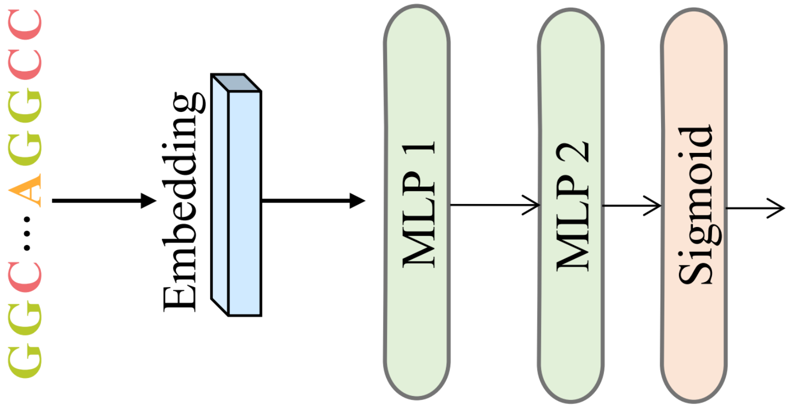


**Supplementary Fig. 2.** Model architecture of the supervised RNA solvent-accessibility prediction (RESM-ASA). Final-layer embeddings from the pre-trained RESM model are passed through a two-layer multilayer perceptron (MLP), followed by a sigmoid activation to predict nucleotide-wise relative solvent accessibility (RSA) values ranging from 0 to 1. Compared to the more complex architecture used in RNA-MSM, this model adopts a simpler and more efficient design while maintaining high predictive performance. The predicted relative solvent-accessibility values are subsequently converted to absolute solvent-accessible surface area (ASA) using predefined normalization factors.

**Supervised siRNA activity prediction**

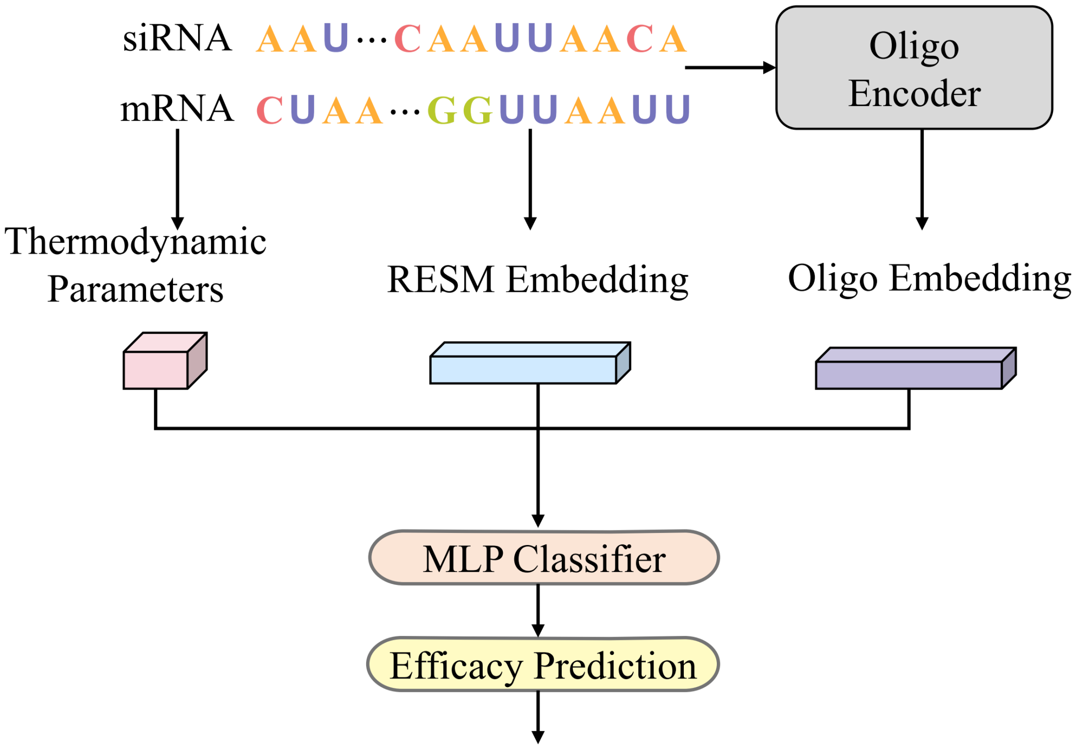

**Supplementary Fig. 3.** Architecture of the supervised siRNA activity prediction model (RESM-siRNA). The framework integrates three parallel feature extraction modules: (1) the RESM Embedding Module encodes both siRNA sequences and flanking mRNA regions using final-layer embeddings from the pre-trained RESM model; (2) the Oligo Encoder processes input sequences using a hybrid architecture consisting of 2D convolutions, pooling layers, a BiLSTM, and a transformer module to capture spatial and sequential interaction patterns; and (3) a thermodynamic feature module extracts siRNA–mRNA pairing properties. The outputs of all modules are concatenated and fed into a multilayer perceptron (MLP) classifier to predict siRNA efficacy scores ranging from 0 to 1. This model adopts the overall design of OligoFormer but replaces RNA-FM with RESM for embedding generation, enabling improved representation learning for RNA-guided targeting tasks.

#### Supervised ribosome loading and expression levels prediction in mRNA

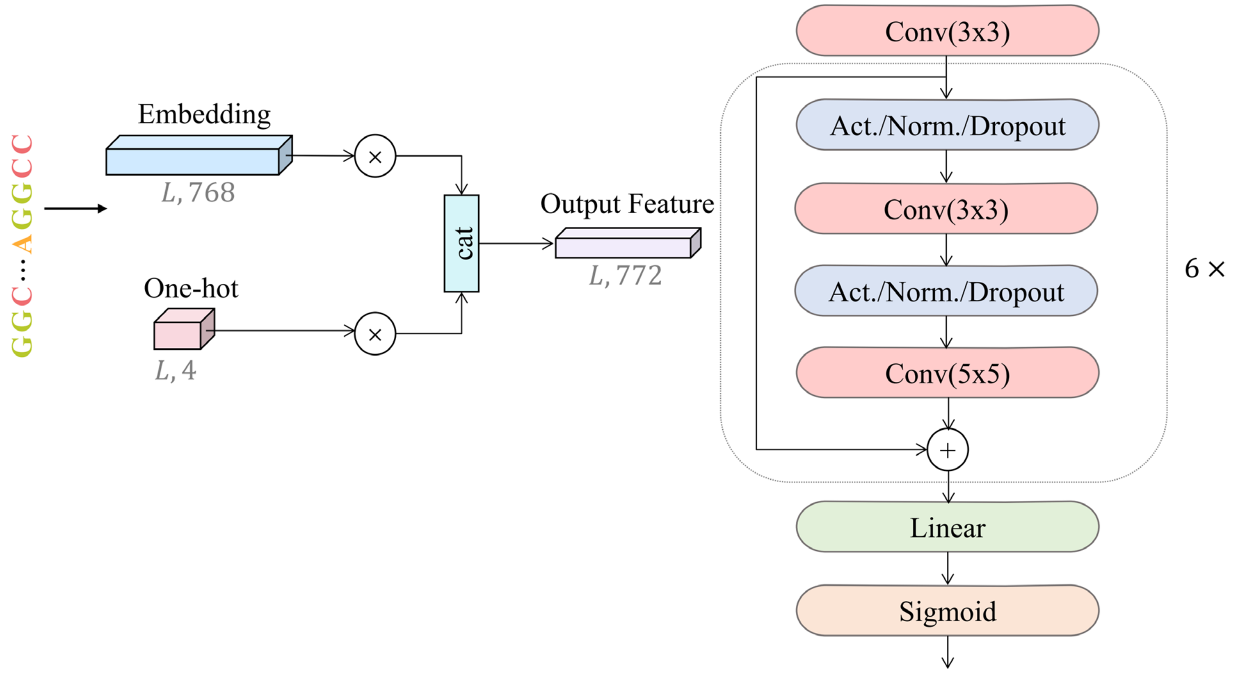


**Supplementary Fig. 4.** Model architecture for prediction of ribosome loading and mRNA expression levels (RESM-riboload and RESM-mRNAexpr). The model integrates one-hot encoded RNA sequences and final-layer embeddings from the pre-trained RESM model. These features are concatenated and projected into a unified representation, which is passed to a ResNet1D prediction head consisting of six residual blocks. Each residual block contains multiple 1D convolutional layers with kernel sizes of 3 and 5, along with activation, normalization, and dropout. A final linear layer and sigmoid activation are used for regression output. This architecture is utilized consistently across both ribosome loading and expression level prediction tasks, enabling robust evaluation of the model's generalization capabilities across diverse biological datasets.
